# Bone adhered soil as a source of target and environmental DNA and proteins

**DOI:** 10.1101/2024.09.10.611648

**Authors:** Toni de-Dios, Biancamaria Bonucci, Rémi Barbieri, Alena Kushniarevich, Eugenia D’Atanasio, Jenna M Dittmar, Craig Cessford, Anu Solnik, John E. Robb, Christina Warinner, Ester Oras, Christiana L. Scheib

## Abstract

In recent years, sediments from cave environments have provided invaluable insights into ancient hominids, as well as past fauna and flora. Unfortunately, locations with favourable conditions for ancient DNA (aDNA) preservation in sediments are scarce. In this study we analysed a set of samples obtained from soil adhered to different human skeletal elements, originating from Neolithic to Medieval sites in England, and performed metagenomics and metaproteomics analysis. From them, we were able to recover aDNA sequences matching the genomes of endogenous gut and oral microbiome bacteria. We also found the presence of genetic data corresponding to animals and plants. In particular we managed to retrieve the partial genome and proteome of a Black Rat (*Rattus rattus*), sharing close genetic affinities to other medieval *Rattus rattus*. Furthermore, we have also been able to reconstruct a partial human genome. The genetic profile of those human sequences matches the one recovered from the original skeletal element. Our results demonstrate that material usually discarded, as it is soil adhering to human remains, can be used to get a glimpse of the environmental conditions at the time of the death of an individual, even in contexts where due to harsh environmental conditions, the skeletal remains themselves are not preserved.

## Introduction

The study of sedimentary ancient DNA (sedaDNA) has profoundly broadened our knowledge about deep human genetic history ^1^. This advance is of crucial importance for certain cases, such as the Neanderthal or the Denisovan, where high quality skeletal samples suitable for ancient DNA (aDNA) studies are scarce or practically non-existent ^2,3^. The application of sedaDNA techniques has allowed the successful reconstruction of ancient environments, providing with inestimable data about past flora and fauna ^4,5^. This has been key to understanding changes in vegetation composition through long periods of times, spanning different environmental and climatological conditions ^6,7^. The use of sedaDNA is not limited to qualitative reconstructions, but also allows for phylogenetic and population genetic analysis of ancient fauna ^8–10^. Furthermore, sedaDNA has been used to retrieve ancient microbial and viral communities ^11–13^. Unfortunately, suitable conditions for sedaDNA preservation are mostly restricted to polar latitudes where soil is frozen in the form of permafrost, to caves where temperatures remain constant all the year around, or to lake sediments ^6,14,15^.

Conversely, palaeoproteomics can be applied in a broader range of environmental conditions ^16,17^, offering complementary and generally better preservation and abundance of biological materials over time ^18–20^. Although proteins provide less genetic data than aDNA, they offer unique and valuable insights into biological evolution, heritable genetic expressions, and functional information of ancient organisms. Palaeoproteomics techniques have facilitated significant discoveries in ancient hominin genetic diversity such as *Homo antecessor* ^21^ and Denisovans ^22^. These methods also help to resolve the taxonomic relationships of various animal species, including extinct birds and mammals ^23–25^. While palaeoproteomics is mainly associated with analysing human or animal remains, adhesives and paint binders, or food residues in pottery^17,26,27^, recent advancements have extended its theoretical application to archaeological sediments ^28^. To the date, ancient soil metaproteomics has only been used successfully to identify and isolate silk proteins from historical periods in specific archaeological contexts ^17,29–31^.

In contrast to aftermention sedaDNA sources, soil adhered to human skeletal remains is a readily available and abundant material that is often discarded during excavation, post-excavation cleaning, or laboratory processing ^32,33^. Bone-adhered soil is extremely persistent in skeleton collections and can be found after meticulous cleaning or years of storage. In some cases, this soil could be the last remaining material that provides environmental data of a site years after its excavation. Despite its abundance, the role of bone-adhered soil as a reliable source of ancient biomolecules is poorly understood. Only few studies have analysed its capability of harnessing authentic ancient microbial DNA or protein sequences ^34^, and none has investigated its potential for endogenous or environmental biomolecules preservation.

Here we present a detailed metagenomic and metaproteomic study of bone-adhered soil from archaeological contexts spanning from the Early Neolithic to the Middle Ages. In it, we demonstrate that this novel source of ancient biological compounds can provide valuable information not otherwise directly retrievable from bones. Additionally, this study opens the door for non-destructive sampling of valuable skeletal remains.

## Results

We targeted 11 individuals from 7 sites in Cambridgeshire, England (Figure 1A, B) We have retrieved a total of 11 samples, from bone-adhered soil originating from temporal bone (PbS), rib (RsS), and vertebrae (VsS) (Figure 1C, Table 1), which after DNA extraction and sequencing, yielded between 15 and 21 million DNA sequences (Supplementary Table S1). An initial metagenomic screening of the samples using Kraken2 ^35^ showed similar abundances of sequences assigned to the domain Bacteria, ranging from 0.56% to 2.31% across samples, while Archaea did not surpass 0.04% in any of the samples. Eukaryotic sequences also display similar proportions among the analysed samples (range 0.08% - 6.34%) (Supplementary Figure 1).

**Figure 1.**
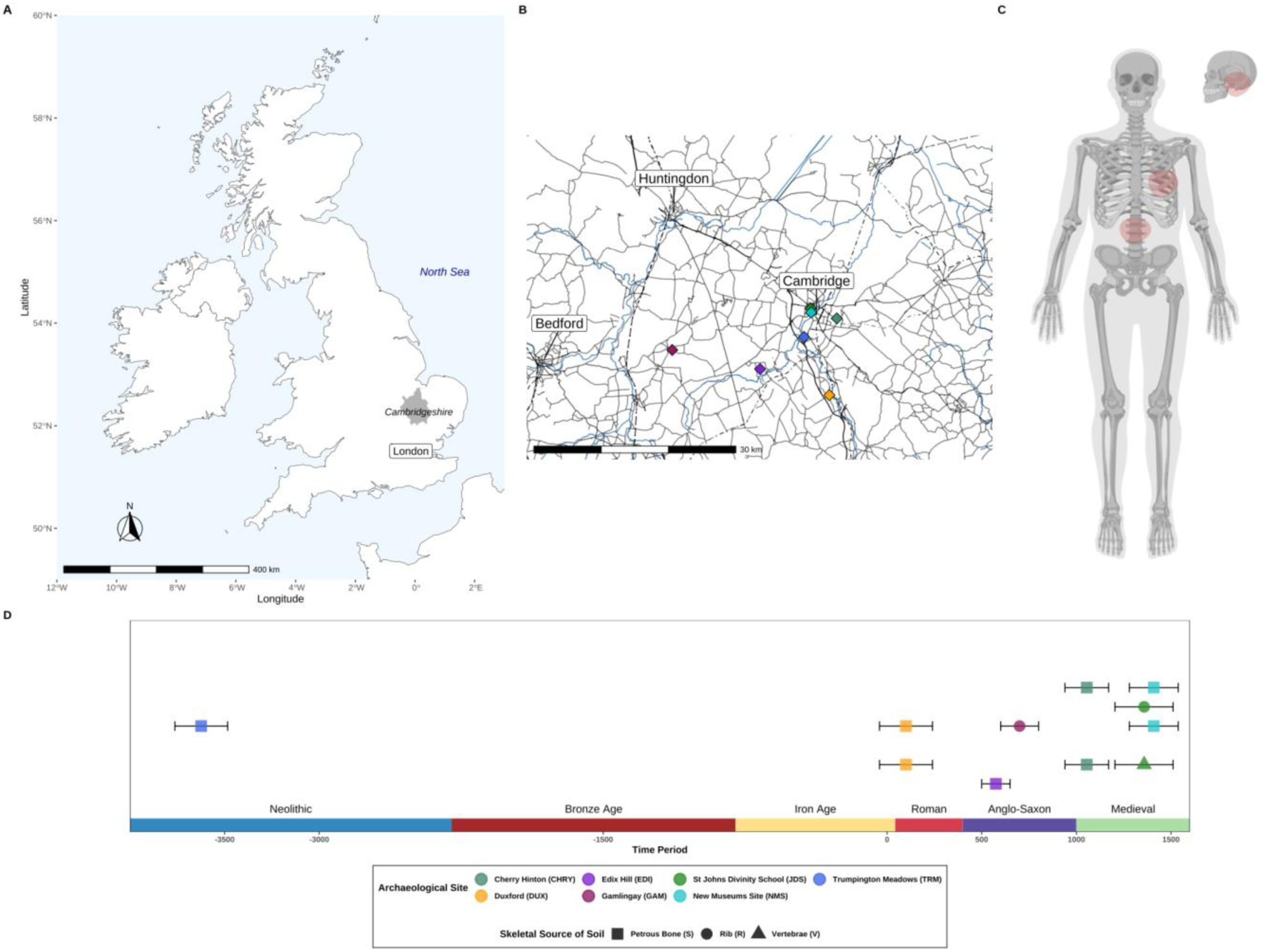
Geographical, chronological and osteological sampling areas used in this study. A) Map of the United Kingdom with the Cambridgeshire area highlighted in grey B) Map of the sites used in this study. C) Highlighted in red, skeletal elements from which the soil was sampled for this study. D) Timeline of the chronological order of the sites used in this study (legend below). Sites are colour coded, skeletal elements designated by shape

**Table 1.** Provenance and summary information for samples used in this study. Individual ID is the project specific number from the After the Plague Project. Mass is the mass of the soil used in the extraction. Some radiocarbon dates for the skeletal elements used have previously been published ^50,51^.

| Lab ID | Individual ID | Site name | Age of site | Skeletal Element | Mass (mg) |
| --- | --- | --- | --- | --- | --- |
| JDS123B | 223 | St John's Divinity School | Medieval | Vertebrae | 6.94 |
| JDS157B | 403 | St John's Divinity School | Medieval | Rib | 3.43 |
| DUX010B | 458 | Duxford | Iron Age / Roman | Petrous | 10.04 |
| DUX012B | 460 | Duxford | Iron Age / Roman | Petrous | 13.75 |
| NMS031B | 509 | New Museums Site | Late Mediaeval | Petrous | 10.76 |
| NMS022B | 518 | New Museums Site | Late Mediaeval | Petrous | 10.7 |
| EDI013B | 559 | Barrington Edix Hill A | Early Mediaeval | Petrous | 8.63 |
| TRM003B | 616 | Trumpington Meadows | Neolithic | Petrous | 15.58 |
| GAM042B | 842 | Gamlingay | Early Mediaeval | Rib | 6.74 |
| CHRY038B | 947 | Cherry Hinton | Medieval | Petrous | 14.68 |
| CHRY051B | 1155 | Cherry Hinton | Medieval | Petrous | 14.05 |

Proteins were also extracted from the soil samples using a protocol modified from ^36,37^. Raw data was analysed using *novor.cloud* and pFind, with searches conducted against Swissprot and cRAP databases ^38^. (see methods). A total of 11,559 and 5,709 peptides were recovered across all samples from 501 and 815 proteins with *novor.cloud* and *pFind* respectively. According to the top hits from each software, the distribution of all identified peptides by source (Human, Bacterial, Non-Human Vertebrates, Plants, Viruses) is nearly consistent between *novor.cloud* and *pFind* (Supplementary Figure S2, Supplementary Table S2). Human peptides are present in each sample. *novor.cloud* identifies human peptides in greater proportion, averaging 57%, compared to *pFind*, which attributes most collagen peptides to non-human vertebrates, averaging 22%. Only four out of 11 samples—DUX012B, TRM003B, GAM042B, and CHRY051B—contain peptides from bacteria, plants, archaea, amoebae, or viruses. Notably, the DUX012B sample has a significant amount of bacterial peptides (87% and 38% with *novor.cloud* and *pFind* respectively) compared to the other samples (Supplementary Figure S3). None of the bacteria identified in the samples could be exclusively (100% identity and coverage) linked to a pathogen or to the human oral, skin, or digestive microbiota.

### Microbial aDNA

After re-screening the samples using KrakenUniq ^39,40^, we generated a bacterial and archaeal assignment dataset based on hits with an E-value above 7 (see methods). The Microbial Source Tracking (MST) analysis of the samples using SourceTracker2 ^41^ showed a heterogeneous landscape in terms of the origin of microbial species, with no clear pattern consistent with the anatomical origin of the sample (Figure 2A, Supplementary Table S3A).

**Figure 2.**
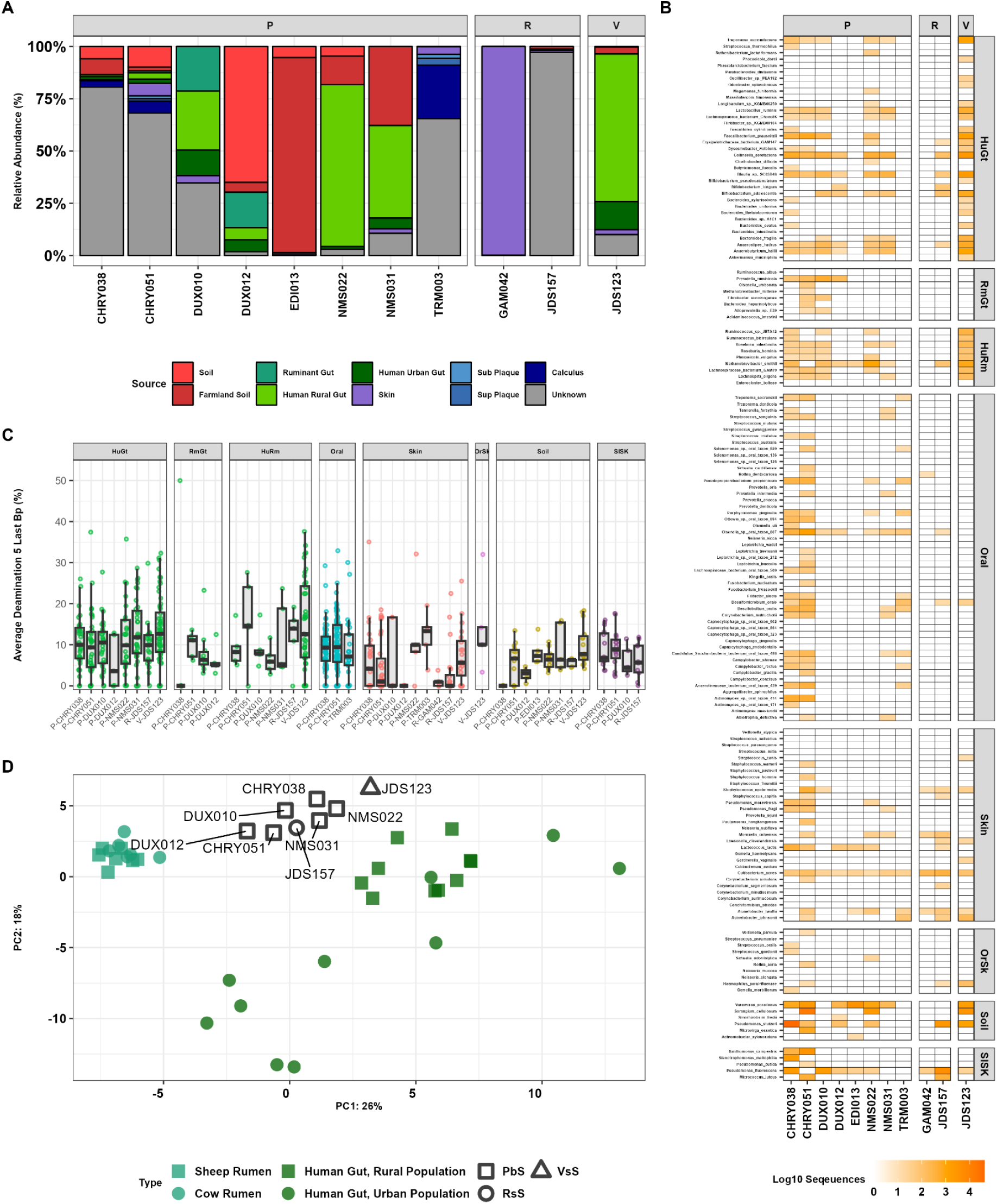
Bacterial composition of the samples in this study. A) Stacked bar chart of MST results. B) Presence of source characteristic bacteria per sample. C) Average terminal damage by each source per sample. D) Principal Component Analysis of gut-associated bacteria in the analysed samples and published gut microbiome data from humans and domesticated ruminants (cow and sheep).

NMS022B, NMS031B and JDS123B display a high abundance of microbial species originating from the human gut microbiome (78.7%, 49.4%, and 84% respectively). DUX10B and DUX12B also exhibit a moderate proportion of human gut microbiota (40.5% and 11.5%), as well as microbial species associated with the rumen of domestic animals (21.3% and 17%). CHRY038B, CHRY051B and TRM003B present traces of human oral bacteria (3.2%, 8.2%, and 30.8%), and the rest of the species present in these samples could not be assigned to any sources used as proxies. Finally, EDI013B, GAM042B and JDS157B seem to be composed in their totality of unique sources: farmland soil, skin and an unknown source respectively (Supplementary Table S3B - S3D). A non-metric multidimensional scaling (nMDS) from the same dataset using Bray-Curtis distances reflects similar tendencies, with most of the samples falling in between the diversity of the used sources, with the exception of TRM003B and GAM042B, which fall as outliers (Supplementary Figure S4).

A closer inspection of the screening data using source-specific species (see Supplementary Data) reveals patterns not obvious using SourceTracker2 alone (Figure 2B). Most of the PbS samples (CHRY, DUX and NMS) contain traces of human gut and/or ruminant gut microbial species. In addition, we have identified the presence of oral bacteria in all PbS samples except for EDI013B (Supplementary Figure S5), including Red Complex Bacteria (*Tannerella forsythia, Treponema denticola* and *Porphyromonas gingivalis*), Sulphur-reducing Bacteria (*Desulfobulbus oralis* and *Desulfomicrobium orale*) and *Olsenella sp. oral taxon 807* (Present in all PbS samples). Finally, we have detected and validated the presence of the pathogenic bacterium *Mycobacterium leprae* sequences in CHRY051B (Supplementary Figure S6, Supplementary Table S4).

Mapping of the sequences assigned to the source-characteristic species and subsequent aDNA damage pattern characterisation reveals the presence of deamination levels compatible with authentic ancient DNA sequences (Figure 2C, Supplementary Figure S7). The damage levels vary depending on the inferred source, but not the skeletal element from which the soil was sampled (Supplementary Table S5), being on average higher in species from the oral and the gut microbiomes (Oral/Skin=13.93% ± 9.61; HumanGut/Rumen=13.27% ± 9.61; HumanGut=11.4% ± 7.55; Oral=9.81% ± 6.24; Rumen=8.5% ± 10.1). On the other hand, sequences originating from possible environmental contamination display lower deamination values (Soil/Skin= 8.1% ± 4.46; Soil=6.57% ± 4.65; Skin=5.57% ± 7.32). Bacterial sequences assigned to skin as a source in NMS022B and TRM003B are exceptions to this trend, both showing an average C to T rate in the last five base pairs of 12.09% and 12.21% respectively, indicating the possibility of being from an endogenous, ancient origin.

Given the abundance of gut-associated microbial sequences in the samples, we decided to further explore the composition by performing a Principal Component Analysis (PCA). This included 79 microbial species found across 33 of the included reference metagenomes (10 Human Urban Gut, 10 Human Rural Gut and 13 Ruminants’ Gut) ^42–45^, and eight of our soil samples (CHRY, DUX, NMS and JDS) (Figure 2D). The reference samples for human urban gut, human rural gut and ruminant gut form three distinct clusters within the PCA (PC1=26%; PC2=18%) (Figures 2D, S8). All soil samples fall within the diversity described by the PCA, forming a cline between samples with higher levels of inferred rumen microbes falling closer to the ruminant gut (CHRY051B and DUX10B) and samples more closely resembling authentic human gut microbiomes from non industrialised populations (NMS022B and JDS0123B). This indicates a certain degree of metagenomic overlap between microbial species abundances originating from humans and domestic animals.

### Non-human eukaryotes metagenomics and metaproteomics

Samples were screened for eukaryotic species presence using a custom pipeline (see methods, Supplementary Figures S9 - S10, Supplementary Table S6), in short: we removed all reads assigned to microbial species and possible contaminants from the dataset as identified by KrakenUniq ^39^, the remaining sequences were re-screened using a database of plastid and mitochondrial DNA from RefSeq ^46^ (Supplementary Figures S11 - S12). We validated sequences according to their E-value with the goal of finding a suitable reference genome. We selected those species with more than three mitochondrial hits per genus and with an E-value above 7. We then performed a set of competitive mappings between the selected reference genomes to determine the best match. Additionally, we calculated edit distance distribution, terminal damage patterns, read length distribution and read coverage of the mapped sequences for each sample and species. Although this was possible for *Fungi* and *Animals*, other organisms such as *Plants* or *Algae* were only classified down to order level due to the high level of homology between retrieved sequences. Despite lower resolution, analysis of terminal damage reveals damage characteristics compatible with those sequences being from an ancient environmental origin (Supplementary figure S13).

After whole genome mapping, we recovered 9,985,882 sequences corresponding to *Homo, Ovis, Bos, Canis, Trichuris, Ascaris, Apodemus* and *Rattus* genera (Figure 3, Table 2) . We also report reads recovered from other species, but these are in very low amounts (Supplementary Figure S14). The sequences display features as length, terminal deamination and edit distance distribution compatible with ancient genomes (Supplementary Figures S15- S17). In the vertebral sample (JDS123B), we have found the presence of sequences belonging to parasitic species (*Ascaris lumbricoides* (roundworm), *Trichuris spp.* (whipworm)). We also find the presence of possible scavengers (*Diptonevra peregrina* (fly), *Hydrotaea spp.* (fly), *Sancassania berlesi* (mite) and *Apodemus sylvaticus* (wood/field mouse)).

**Figure 3.**
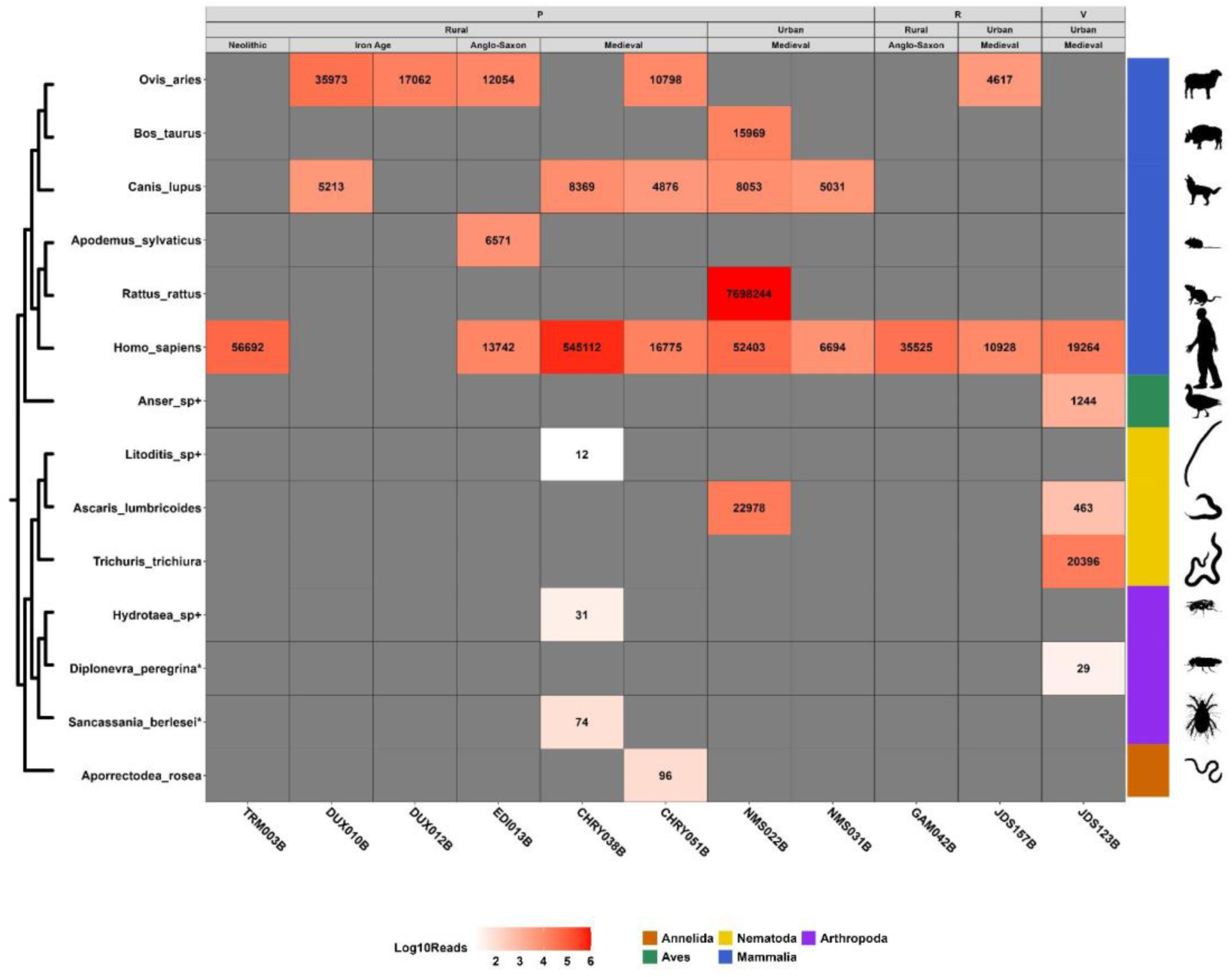
Number of reads from a select panel of animal species present in the samples. Read assignments were determined using a custom screening pipeline described in this study (see methods). Vertical axis is ordered by the taxonomic location of detected species, with some species marked by * (Only mitochondrial reference available) and + (Taxonomic level could not be determined below genus level). Horizontal axis is the sample. Samples are grouped by their sampling origin (P = Petrous Bone Soil, R = Rib Soil, V = Vertebrae Soil), context of site (Rural vs Urban), and site datation (Neolithic, Iron Age, Anglo-Saxon and Medieval). Read numbers are in log10 scale with higher numbers displayed in a darker red.

**Table 2.** Mapping statistics of different animals retrieved. Mapping statistics of the identified animals in the samples, those include number of mapped sequences, average edit distance, average deamination in the last 3 bp of each read and average depth of coverage. Only Animals with whole genome data have been included.

| Individual | Specie | Mapped Reads | Av. Damage last 3 bp | Av. Edit Distance | Average Coverage |
| --- | --- | --- | --- | --- | --- |
| CHRY038B | *Homo sapiens* | 545,112 | 0,138 | 0,62 | 0,0099X |
| CHRY038B | *Canis lupus* | 8,369 | 0,119 | 1,348 | 0,0002X |
| CHRY051B | *Homo sapiens* | 16,775 | 0,109 | 0,608 | 0,0004X |
| CHRY051B | *Canis lupus* | 4,876 | 0,145 | 1,41 | 0,0001X |
| CHRY051B | *Ovis aries* | 10,798 | 0,143 | 1,39 | 0,0003X |
| DUX010B | *Canis_lupus* | 5,213 | 0,075 | 1,214 | 0,0001X |
| DUX010B | *Ovis aries* | 35,973 | 0,086 | 0,89 | 0,0007X |
| DUX012B | *Ovis aries* | 17,062 | 0,067 | 0,918 | 0,0003X |
| EDI013B | *Apodemus sylvaticus* | 6,571 | 0,112 | 1,1472 | 0,0001X |
| EDI013B | *Homo sapiens* | 13,742 | 0,186 | 0,962 | 0,0002X |
| EDI013B | *Ovis aries* | 12,054 | 0,167 | 1,275 | 0,0002X |
| GAM042B | *Homo sapiens* | 35,525 | 0,134 | 0,829 | 0,0005X |
| JDS123B | *Anser spp.* | 1,244 | 0,097 | 2,174 | 0X |
| JDS123B | *Trichuris trichiura* | 20,386 | 0,128 | 1,433 | 0,0152X |
| JDS123B | *Ascaris lumbricoides* | 463 | 0,118 | 1,687 | 0,0001X |
| JDS123B | *Homo sapiens* | 19,264 | 0,159 | 0,878 | 0,0003X |
| JDS157B | *Homo sapiens* | 10,928 | 0,072 | 0,906 | 0,0002X |
| JDS157B | *Ovis aries* | 4,617 | 0,086 | 1,631 | 0,0001X |
| NMS022B | *Rattus rattus* | 7,698,226 | 0,065 | 0,551 | 0,1842X |
| NMS022B | *Ascaris lumbricoides* | 22,978 | 0,142 | 1,149 | 0,0042X |
| NMS022B | *Homo sapiens* | 52,403 | 0,141 | 0,92 | 0,0009X |
| NMS022B | *Canis lupus* | 8,053 | 0,126 | 1,328 | 0,0002X |
| NMS022B | *Bos taurus* | 15,969 | 0,108 | 1,079 | 0,0003X |
| NMS031B | *Homo sapiens* | 6,694 | 0,142 | 0,944 | 0,0001X |
| NMS031B | *Canis lupus* | 5,031 | 0,133 | 1,3267 | 0,0001X |
| TRM003B | *Homo sapiens* | 56,692 | 0,22 | 0,847 | 0,0009X |

Perhaps most interesting, within NMS022B, we found sequences amounting to a depth of 0.1877× for the nuclear and 8.09× for the mitochondrial genome of the black rat *(Rattus rattus)*. By analysing the heterozygosity level of the mitochondrial genome (0 after filtering for potential aDNA damage), we conclude that the sequences most likely come from a unique individual, most probably male given the ratio of X and Y chromosome sequences (1/1, 1 to 2 autosomes) (Supplementary Figure S18). We created a genome-wide SNP dataset of modern and historical *R. rattus* (Supplementary Table S7) and performed a PCA using a published set of ancient ^47^, modern rat genomes ^48^, and our ancient genome (Figure 4A, Supplementary Figure S19). The NMS022B ancient rat displays affinities to other Western and Northern European Medieval and Early Modern rats.. In order to formally validate this result we performed an *f*4 analysis (D statistics) in AdmixTools ^49^, showing that indeed, this particular individual is genetically closer to other Mediaeval black rats (Figure 4C).

**Figure 4.**
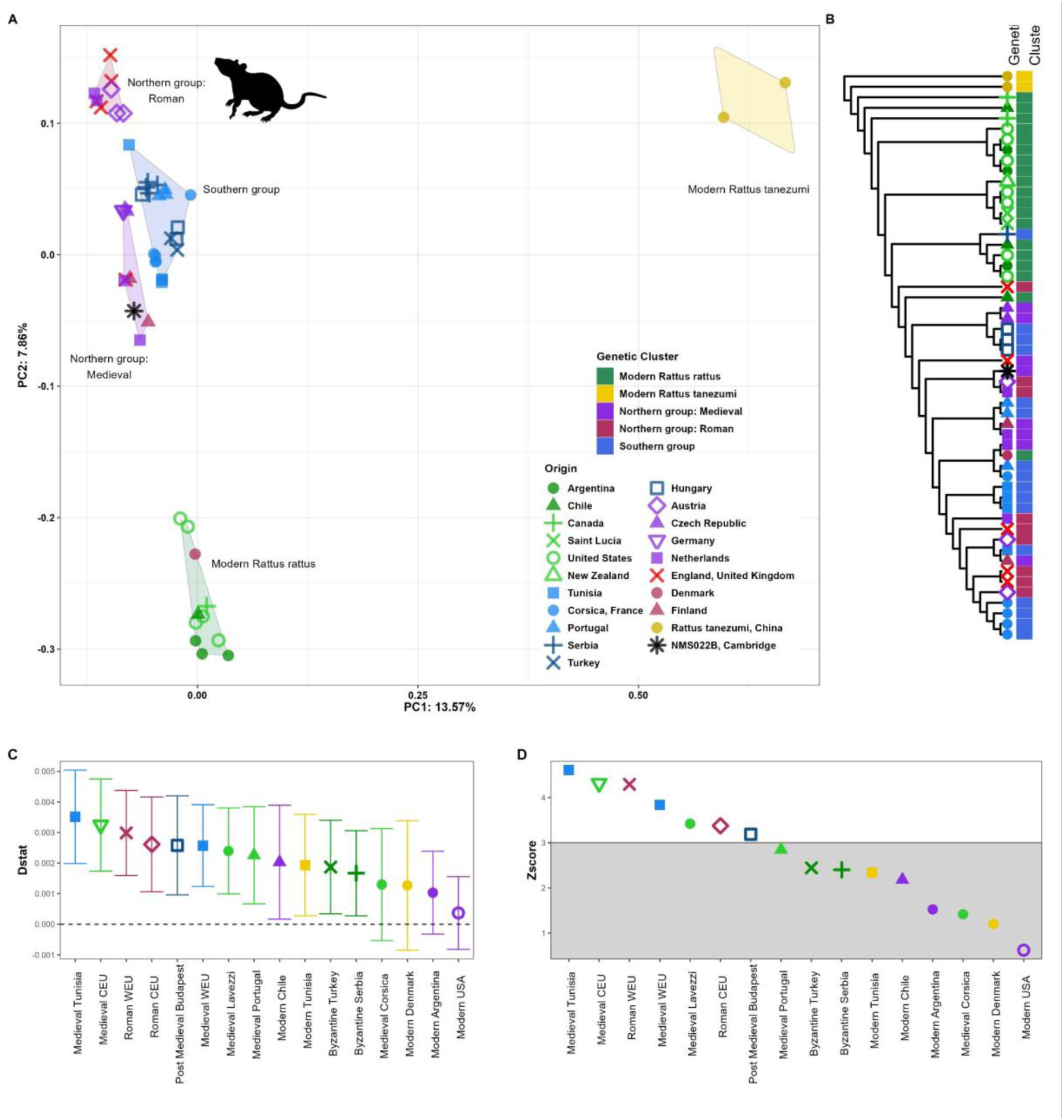
Population genomic analyses of the ancient rat genome recovered from NMS022B. A) PCA of NMS022B and 51 modern and ancient Rats. Cluster names are annotated in figure. B) Mitochondrial Tree. Clusters are annotated in colour-coded format). C) f4-statistics (Dstat) under the test relationship f4(Rattus tanemuzi, NMS022B, Modern_Canada, X) of NMS022B sample and other historical and modern *Rattus rattus* populations. D) Z-scores under the test relationship assessed through block jack-knife resampling.

The protein data also supports the presence of an individual black rat in the sample. The NMS022B sample contains the most unique peptides (2,363 and 1,141) and protein count (96 and 125) with *novor.cloud* and *pFind* respectively. This sample also contains a higher number of peptides assigned to *R. norvegicus* (1,436 and 148) or *M. musculus* (295 and 269) against SwissProt (Figure 5a) which are not found in any of the other samples or the extraction blank.

**Figure 5.**
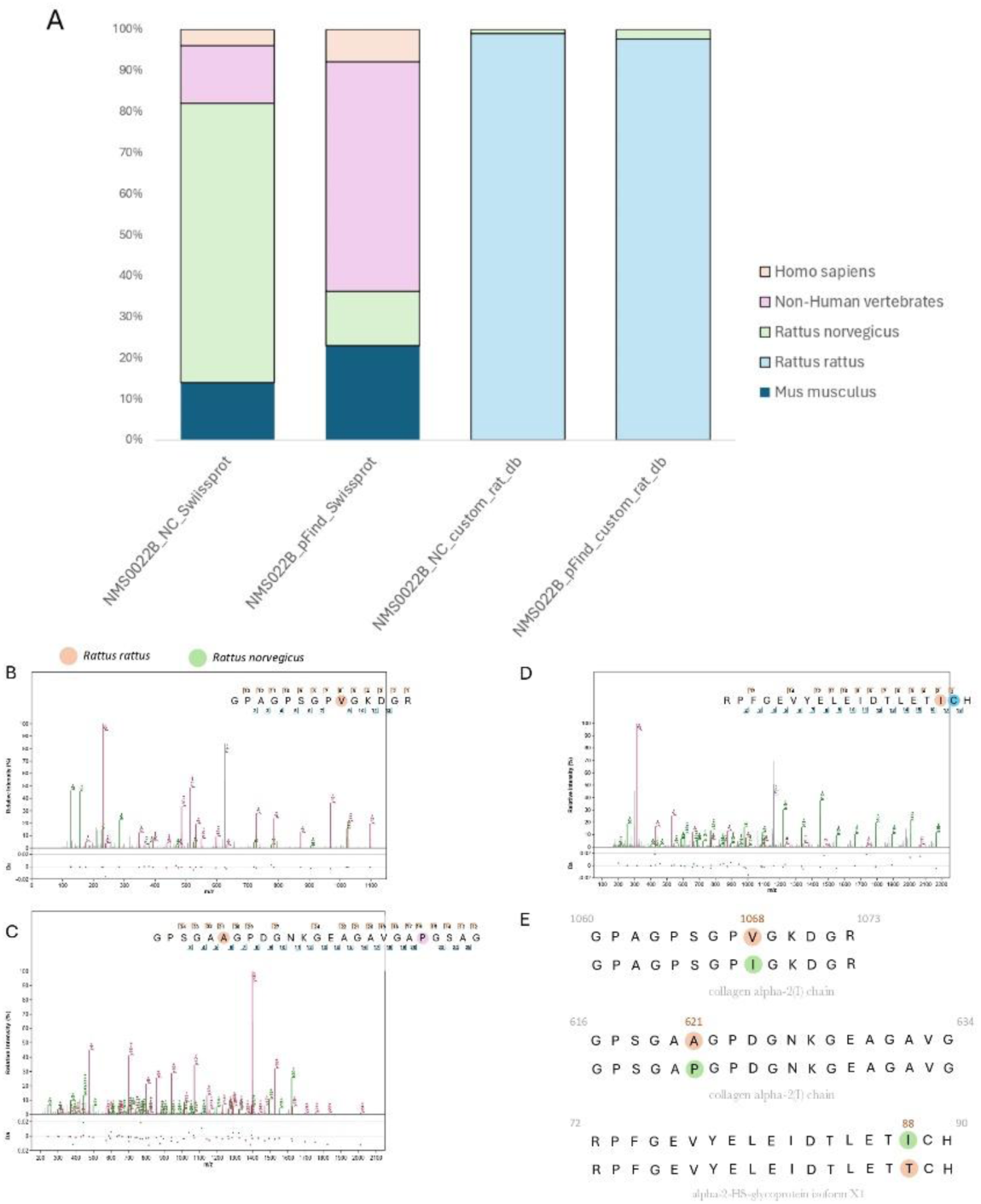
*Rattus rattus* specific peptide identification from sample NMS022B. (a) stacked bar plot showed the relative abundance of peptide origins based on Swiss-Prot identification from *novor.cloud* and *pFind* outputs and our custom rat database. (b) Spectrum from GPAGPSGPVGKDGR (collagen alpha-2(I) chain) identified by *pFind* specific to *Rattus rattus*. (c) Spectrum from GPSGAAGPDGNKGEAGAVGAPGSAG (collagen alpha-2(I) chain) identified by *pFind* specific to *Rattus rattus*, the purple dots indicate hydroxylated proline residues.(d) Spectrum from RPFGEVYELEIDTLETICH (alpha-2-HS-glycoprotein isoform X1) identified by *pFind* specific to *Rattus rattus*. Blue circle represents carbamidomethylation of cysteine. (e) Substitutions in the amino acid sequences from peptides of collagen alpha-2(1) chain (position 1068 and 621) and alpha-2-HS-glycoprotein isoform X1 (positions 88) identified with both *novor.cloud* and *pFind* can be used to distinguish between *Rattus rattus* and *Rattus norvegicus*.

The SwissProt database, which exclusively contained protein sequences of *R. norvegicus*, limited our ability to obtain peptides from other rat species. Consequently, we developed a database incorporating all published protein sequences from *R. rattus* and *R. norvegicus* on NCBI. We identified a total of 2,758 peptides on this occasion, with 2,719 (99%) associated with *R. rattus* and 39 (1%) with *R. norvegicus* (Figure 5a). Our focus was on analyzing specific peptides to distinguish between these two species. In confirming the identity of our peptides as belonging to *R. rattus*, we focused on the GPAGPSGPVGKDGR and GPSGAAGPDGNKGEAGAVGAPGSAG peptide sequences from collagen alpha-2(1) chain, and the RPFGEVYELEIDTLETICH peptide sequence from the alpha-2-HS-glycoprotein isoform XI protein(Figure 5b,c,d,e). This scrutiny enabled us to pinpoint differences on three specific amino acid substitutions allowing the differentiation of *Rattus rattus* from *Rattus norvegicu*s species (Figure 5b,c,d), affirming the reliable identification of ancient proteins from *R. rattus* in medieval Europe. This resulted in a total of 44 proteins originating from different tissues with a coverage above 5% (Supplementary Table S8).

### Human aDNA from soil

Substantial human aDNA sequences were recovered from all samples with the exception of samples from the Duxford site (Supplementary Table S9). The abundance of endogenous human aDNA does not seem to relate to the endogenous abundance found in the original bone ^50,51^ (Figure 6A), while terminal C to T deamination is comparable to the original sample (Figure 6B) . Genetic sex characterisation of the human sequences adjust to that of the original skeletal element sequences (Supplementary Figure S20), although it is important to remark that in most of the cases the endogenous content is extremely low and the results may not be accurate. A special case is sample CHRY038B, which yielded a human genome at a depth of roughly 0.01× (3% of endogenous DNA). Genetic sex and mitochondrial haplotype match those of the original petrous bone (XX, U5b3e) ^50^ (Supplementary Table S9B). The pseudo-haploid-called nuclear genome contained enough sequences to recover 8,580 SNPs from the Human Origins dataset ^52^. We also imputed the genome using the UK Biobank panel ^53^ and projected it against a panel of modern European populations ^52^. We used READv2 ^54^ to compare the original bone sample and soil-adhered library, with the result of ‘IDENTICAL’, both with and without imputation (Supplementary Figure S21, Supplementary Table S10-S11). The imputed sample falls close in the PCA to the original sample (from bone) amongst the samples from Western Europe (Figure 6C, Supplementary Table S11). We also pulled imputed phenotypes to check the concordance between information in the original bone sample and bone-adhered soil (Supplementary Table S13). Applying a genotype dosage (hereby indicated as DS) of 0.1, which is the standard used in imputation, we observed a concordance of approximately 73.4% with 29 genotypes of the total 109 phenotypic-informative markers differing between the two sample sources (Supplementary Table 13).

**Figure 6.**
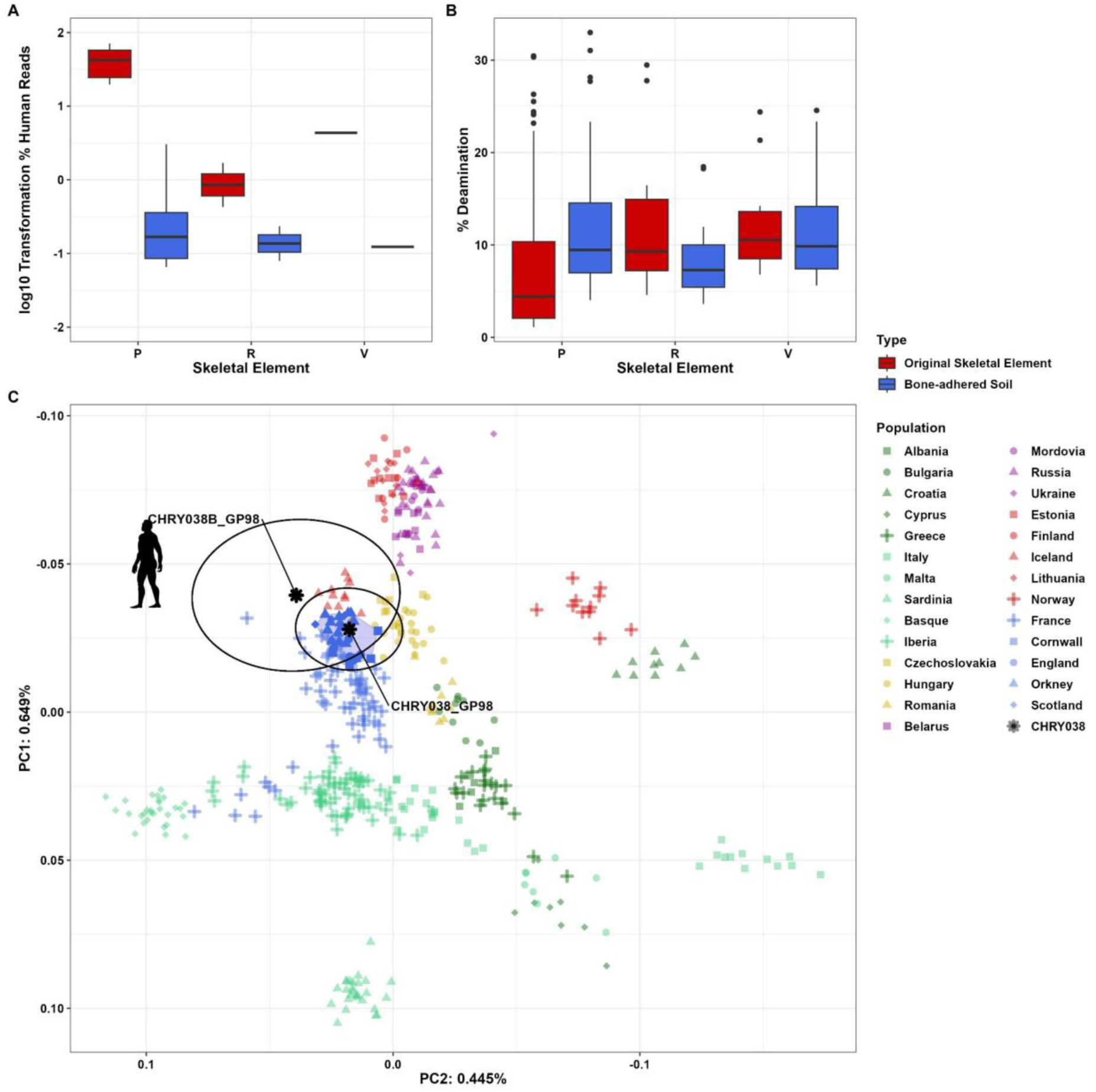
Human DNA content. A) Log10 transformation of the percentage of endogenous human aDNA from original skeletal element versus soil recovered from that element. B) Deamination proportion at the 5 last positions in both reads’ ends from original skeletal element versus soil recovered from that element C)Principal Component Analysis of the human aDNA from the soil sample of CHRY038B and the tooth sample of CHRY038. The PCA was created using smartpca, with CI 95 displayed by black circle using *ellconf: 0.95,* and projecting our ancient samples against 442 modern samples from Europe. Modern British samples are highlighted in the blue polygon.

## Discussion

Our results show that authentic traces of ancient microbiome, endogenous, and environmental ancient biomolecules can be retrieved from bone-adhered soil sampled from archaeological remains. This is the case for both human gut and oral microbiome. Ancient gut microbiomes have been previously reported from paleofeces ^27,55^. Although putatively ancient, the reported gut microbiomes seem to be a mixture of both human and domestic animal gut microbiota, suggesting a high degree of environmental noise. Even if only considering human sequences, we cannot discard contemporaneous contamination due the lack of hygiene measures at the time. Nonetheless, if this was the case, those metagenomes could be representative of the whole population, and the composition is comparable to other ancient gut microbiomes ^55^. In contrast, the simplest explanation for the presence of oral signal in the samples seems to be an endogenous source, leaching from the oral cavity to the inner ear and petrous bone area during the decomposition process ^56^. The presence of oral bacteria can hinder the use of this type of material as environmental controls in oral metagenomic studies. Apart from typical human commensals, we found potentially pathogenic microbes including red complex bacteria and *Mycobacterium leprae*. For the case of *M. leprae,* although that particular individual was not infected with the bacterium, other individuals in the site were tested positive. The presence of *M. leprae* sequences in the surrounding soil could be evidence of DNA leaching during the decomposition process of the body.

Despite most of the samples originating from soil attached to the petrous bone, which is a priori relatively isolated from the exterior media, there is a surprising abundance of environmental sequences. This is the case for plant aDNA, mainly assigned to shrub or grasses phyla (although some potential tree aDNA can be found in the Neolithic and Iron Age sites), and that we can find in most of the samples; this is also confirmed by metaproteomics, where the samples with highest plant aDNA also present the highest amount of plant peptides. Animal sequences are also widespread across all the sample types and sites. Those include common domestic animals such as dog, cow, sheep and a geese species; or scavengers such as the woodmouse or different fly species. Those could have left DNA traces due to proximity to the deposition site or by actively interacting with the body. An interesting case is that the sites where we find *Ovis* aDNA correlates with more rural sites (Duxford, Cherry Hinton and Edix Hill) where sheep grazing during or after the burial period would have been more common. Further evidence supporting this is that samples displaying more *Ovis* aDNA also display a higher content of ruminant microbial sequences. We also found potential traces of human intestinal parasites (pinworms and whipworms) and if this is a target for research, then bone-adhered soil from the pelvic region, i.e. pelvis or vertebrae, could be a good source. Pelvic soil is already being used as a source for parasite studies ^57,58^, but if this is not available because excavation was in the past, bone-ahered soil from stored remains could still be a valuable source.

The paleoproteomics results somewhat mirror those of the aDNA, which served as a reference control. In addition to *Homo sapiens* specific peptides, our samples exhibit a significant abundance of probable peptides from other mammals, likely attributable to *Bos* sp. and *Canis* sp. Furthermore, soil bacteria, environmental fungi, potential parasites, archaea, amoebae, and plant viruses were also detected in three samples (TRM003B, DUX012B, GAM042B). One sample (TRM003B) yielded a high abundance of aDNA sequences and peptides belonging to the *Poaceae* and *Brassicaceae* families. Unfortunately, these are not well-identified (few peptides not only specific to the species of interest), so they cannot be confirmed with proteins alone but could be interpreted as a reflection of the high potential diversity of these bone- adhered soil samples. Palaeoproteomics results confirm the feasibility of isolating and identifying ancient proteins from soil samples using metaproteomic techniques. This allowed us to analyse for the first time soil samples using both proteomics and aDNA techniques, generating consistent results between the two analysed ancient biomolecules.

In the urban context, the finding of a partial genome of a black rat supported with palaeoproteomics in a site containing plague victims ^59,60^ is particularly interesting. How the ancient rat genome and proteome came to be incorporated in the soil inside the petrous bone of this individual is unclear. Analysis of protein origin (e.g. saliva, urine, faeces, blood) does not provide any further information, as proteins are evenly distributed between blood, collagen, bone, cartilage or other tissue proteins (Supplementary table S8), and are not associated with any particular tissue or body fluid that might shed light on the rat’s aDNA and ancient proteins introduction in soil. The genome of the rat is clearly ancient, but whether it is contemporary with the burial is uncertain. Rat bones were not found near the body in the original excavation; due to their small size they are often missed and difficult to recover but given the care taken excavating human skeletons would probably have been identified if present, and there was no evidence of gnawing on the bones ^61–63^. The black rat went extinct in most of eastern England by the 18th century, due to competition from the brown rat ^64^. The finding of a black rat genome is also interesting as the Black Death spread to England from Europe. *Y. pestis* has been found in ancient rat remains ^65^; however, no trace was found in this soil. It is possible that this rat was not a carrier or infected with *Y. pestis*. A possible explanation for the presence of rat aDNA could be from faeces or urine deposited on or near the body before or during burial. The soil in the grave fill appears to be the material that was excavated to create the grave, as the site had been occupied for several centuries prior to becoming a cemetery it is possible that the rat aDNA predates the burial and relates to the earlier occupation. We cannot rule out leaching through the soil; however, the sample was taken from deep inside the petrous bone, making this scenario less likely. These findings have been corroborated by palaeoproteomics results validated by two independent tools, *novor.cloud* and *pFind*. To our knowledge, this represents the first reported ancient draft proteome of *R. rattus*, encompassing 42 partially covered proteins (>5%) (Supplementary table S8).

Additionally, the recovery of a partial genome of *R. rattus* with aDNA, along with the analysis of specific aDNA degradation patterns could be indicative that the identified *R. rattus* peptides are likely dated from the same Mediaeval period. This could constitute a remarkable and rare correspondence between aDNA and ancient proteins. These preliminary results demonstrate the feasibility of identifying a substantial number of ancient endogenous peptides from complex soil samples containing modern contaminants, environmental bacteria, and ancient endogenous proteins. This suggests that soil metaproteomics is a promising, albeit yet to be fully confirmed, alternative to the destructive analysis of valuable archaeological material offering insights into ancient environments, fauna, flora, and pathogens.

Even though the bones used in this study were cleaned at excavation years before sampling and were stored in various controlled and non-controlled environments, soil was still present and provided valuable biomolecular information. The human aDNA in bone-adhered soil varies greatly between samples and sample type; however, in certain circumstances enough coverage can be obtained to reveal valuable population genetics information including ancestry, genetic sex and even phenotypes. Furthermore, other individuals show enough endogenous DNA content to make them suitable candidates for target capture ^66^.

## Conclusion

Bone-adhered soil is a unique and valuable resource for obtaining ancient genomic and proteomic data in a non-destructive manner. This study highlights the potential of bone- adhered soil for studying the ‘death biome,’ scavenging of ancient bodies in a non-destructive manner. Due to the amount of shared biomolecular material between the burial and the soil, petrous soil should not be treated as a purely environmental control for metagenomic or proteomic analysis.

## Methods

All of the laboratory work was performed in dedicated ancient DNA facilities of the Institute of Genomics, University of Tartu. The library quantification and sequencing were performed at the Estonian Biocenter Core Laboratory. The main steps of the laboratory work are detailed below.

### Selection of archaeological samples

Soil from inside the ribs (samples GAM042B, JDS157B), vertebra (JDS123B) and petrous bones (all other samples) was removed from the bone using a dental scraper cleaned between uses with 6% v/w NaOCl and rinsed with double distilled water and 70% ethanol. Soil was collected into DNA lo-bind 1.5 ml tubes (Eppendorf).

### Extraction of ancient DNA and protein data

To each soil sample, 500 µl of 0.5M EDTA pH 8.0 was added and samples were rocked at room temperature overnight. From the extracts, 450 µl of supernatant were removed and purified according to manufacturer’s instructions using buffers from the Minelute^TM^ PCR Purification Kit (Qiagen) with changes as described in the online protocol ^67^ and the following project specific update: samples were eluted in 60 µl of EB buffer instead of 100 µl. One extraction was performed per sample for screening and 30μl used for libraries.

Library preparation was conducted using a protocol modified from the manufacturer’s instructions included in the NEBNext® Library Preparation Kit for 454 (E6070S, New England Biolabs, Ipswich, MA) as detailed in ^68^ except that libraries were single indexed and not split into two amplifications for PCR. DNA was amplified using the following PCR set up: 50μl DNA library, 1X PCR buffer, 2.5mM MgCl2, 1 mg/ml BSA, 0.2μM inPE1.0, 0.2mM dNTP each, 0.1U/μl HGS Taq Diamond and 0.2μM indexing primer. Cycling conditions were: 5’ at 94°C, followed by 18 cycles of 30 seconds each at 94°C, 60°C, and 68°C, with a final extension of 7 minutes at 72°C. Amplified products were purified using MinElute columns and eluted in 35 μl EB (Qiagen). Three verification steps were implemented to make sure library preparation was successful and to measure the concentration of dsDNA/sequencing libraries – fluorometric quantitation (Qubit, Thermo Fisher Scientific), parallel capillary electrophoresis (Tapestation, Agilent) and qPCR. Samples were shotgun-sequenced to a depth of approximately 20 million reads on the Illumina NextSeq500/550 using the High-Output single-end 75 base pair kit at the University of Tartu Institute of Genomics Core Facility. The ancient DNA data generation from the original skeletal elements was done as described in the relevant publications ^50,51^.

The pellet and remaining 50µl supernatant were used for protein isolation using a modified version of the SP3 protocol used in ^26^ and available on protocols.io ^36^. To the pellets, 150µl of 2M guanidine HCl was added, then 20 µl each of 100mM CAA and TCEP. Samples were incubated at 99C for 10 minutes then allowed to cool. To the sample, 10µl of a mix of Sera-Mag Carboxylate SpeedBeads (hydrophobic Cat. #45152105050250 and hydrophilic Cat. #65152105050250) was added along with 230 µl of 100% Ethanol and incubated at 24°C for 5 minutes at 500 rpm. Beads were pelleted using a magnetic rack and the supernatant removed. Beads were washed three times with 80% ethanol then 150µl of TEAB and 1µl trypsin were added to each tube. Samples were incubated overnight at 37°C at 500 rpm. In the morning, Pierce™ C18 Tips, 100 µL beds were used to immobilise the peptides. Tips were prepared with methanol, AT80 and 0.1% TFA. Samples were run through the tips twice and followed by two rounds of 0.1% TFA. Tips were stored at -20°C until delivered to the Proteomics Core Facility at the University of Tartu.

Samples were eluted from C18 StageTips and reconstituted in 21 ul of 0.5% TFA. LC-MS/MS analysis was carried out by loading the entire sample to a 0.3 × 5 mm trap-column (5 µm C18 particles, Dionex) using an Ultimate 3500 RSLCnano system (Dionex, California, USA). Peptides were eluted to an in-house packed (3 µm C18 particles, Dr Maisch, Ammerbuch, Germany) analytical 50 cm × 75 µm emitter-column (New Objective, Massachusetts, USA) and separated at 250 nl/min with an A to B 5-35% 1.5 h gradient (buffer A: 0.1% formic acid, buffer B: 80% acetonitrile + 0.1% formic acid). Both the trap- and analytical columns were operated at 40°C. Separated peptides were on-line electrosprayed to a Q Exactive Plus (Thermo Fisher Scientific) mass spectrometer via a nano-electrospray source (positive mode, spray voltage of 2.6 kV). The MS was operated with a top-10 data-dependent acquisition strategy. Briefly, one 350-1,400 m/z full MS scan at a resolution setting of R = 70,000 at 200 m/z was followed by higher-energy collisional dissociation fragmentation (normalised collision energy of 26) of the 10 most intense ions (z: +2 to +6) at R = 17,500. MS and MS/MS ion target values were 3,000,000 and 50,000 ions with 50 and 50 ms injection times, respectively. MS/MS isolation was carried out with 1.5 m/z isolation windows. Dynamic exclusion was limited to 30 s. Peptide match was set to “Preferred” to select isotopic features reminiscent of peptides for data-dependent scanning.

### Bioinformatic quality control and initial processing of sequences

Raw data was returned in the form of single FASTQ files. As a preliminary step, the sequences of adaptors and indexes and poly-G tails occurring due to the specifics of the NextSeq500/550 and Hiseq2500 technology were removed from the ends of DNA sequences using AdapterRemoval2 ^69^. In order to avoid spurious matches in the following analysis, we discarded all sequences shorter than 30bp and with a quality below 20. (For further information regarding all the software used and their versions see Supplementary Table S14).

### Metagenomic screening

To further reduce the spurious assignment of generated sequences to evolutionary conserved regions, we removed low complexity sequences using prinseq and a dust value of 7 ^70^. Finally, we collapsed all duplicated reads using bbmap tool suite ^71^. Following this, an initial metagenomic screening of the deduplicated sequences was performed, using *Kraken2* ^35^ against the *nt* database as for 11/29/2023 (available at https://benlangmead.github.io/aws-indexes/k2) against the newly generated libraries and their skeletal element counterpart (Supplementary Table S15).

For an in depth microbial metagenomic screening we started by purging the deduplicated sequences of human reads. To do this we mapped them against the human reference genome 37 using BWA *bactrack* ^72^, with edit distance and seeding tweaked to account for aDNA damage (-n 0.01, -o 2 and -l 10,000) ^73^. All sequences with mapping quality of 0 or higher were discarded, and the resultant reads were processed with KrakenUniq and the standard MicrobialDB as for 16/08/2020 ^39^. This database includes data belonging to *Bacteria, Archaea, Virus* and Eukaryotic pathogens. Additionally, it also includes the Human Reference genome 37, and a set of possible contaminant sequences.

For a more accurate detection of Eukaryotic sequences, a second screening was performed on the deduplicated sequences using KrakenUniq with a custom database. This included all published mitochondrial and plastid genomes, the human reference genome, and a dataset of known contaminants and synthetic sequences as for 08/02/2022. After selecting potential targets, we performed a series of competitive mappings against their respective reference genomes in order to avoid cross mapping between phylogenetically close species (See Supplementary Methods, Supplementary Figure S23). This resulted in a set of animal species filtered bam for each different sample analysed.

### Microbial metagenomic analysis

Raw KrakenUniq reports were filtered by using the E-value formula described by Guellil and Borry, derived from the proportion of *kmers* by read per genome coverage in each taxonomic level ^74^. Higher values of this statistic denote a higher distribution of the matching hits along the reference genome, and thus, the higher the probability that the hit is genuine. We have considered as valid those microbial species with an E-value of 7. To estimate the proportion of reads originating from various microbial sources, we have used Sourcetracker2, a source- prediction software with a Bayesian framework ^75^. The analysis with Sourcetracker was performed on normalised bacterial taxa reads abundance at the species level. We have used a custom metagenomic dataset of sources, including modern human dental calculus, human oral plaque, skin, soil, gut and domestic animal gut (see Supplementary Methods Section S1.1, Supplementary Figures S24-S26, Supplementary Table S16). We then merged each filtered sample report to the reference dataset, keeping species with at least 200 reads in the Reference samples, and 50 reads in the target sample (to account for lower abundance of microbial reads in ancient samples), and discarding species with an abundance below 0.02% in the entire dataset. Sourcetracker2 was then runned with our target sample as sink, using a refraction value of 100 for sinks and sources. We have used the same dataset to perform a non-Metric Dimensional Scaling (nMDS) from Bray-Curtis distances using R package *vegan* ^76^.

Damage per source was estimated by mapping sequences to species characteristic of each source as inferred by Sourcetracker2. To facilitate the process we extracted reads by taxonomy ID using krakentools ^77^ and mapping those same sequences to the respective reference genome with *BWA bactrack* with astringent settings (-e 0.1). Mapped sequences were then analysed for terminal cytosine deamination using MapDamage2 ^78^. We have calculated the average deamination ratio in the last 5 bp of the read in those species with more than 100 sequences mapped. Additionally we analysed GC content in the different microbial sequences retrieved (see Supplementary Methods Section S1.2, Supplementary Figures S27).

We extracted species assigned by sourcetracker2 to be characteristic of human and ruminant gut microbiomes from our samples’ KrakenUniq raw reports, and from the references raw reports. We applied an Evalue filter of >=7 and a read number of >=10. Once the individual reports were filtered, we merged all the reports into a merged dataset, normalising for sample library size, and discarding in the process species with a representation below 0.02% in the total of the dataset. This filtered dataset was further normalised using a *clr* transformation in the R package *compositions* ^79^, and a Principle Component Analysis (PCA) was computed using R package *mixOmics* ^80^. Resultant sample values and loadings were visualised using R. We performed similar analysis for Oral microbes found in samples CHRY038B, CHRY051B and TRM003B (see Supplementary Methods Section S1.3, Supplementary Figures S28).

Microbial species of interest were filtered from the raw KrakenUniq report using the same Evale formula. We then extracted them from the raw *fastq* using krakentools and validated using a combination of megablast and MEGAN6 ^81,82^.

### Modern and ancient contamination

We used schmutzi to detect the presence of modern human contamination in our mapped sequences. This method considers the endogenous deamination rates of the sequences ^83^. We applied schmutzi to both nuclear and mitochondrial human sequences. In addition to that, we also have used contamix to infer contamination at a mitochondrial level that could originate from modern sequences or as a result of the mixture of genuine ancient sequences.

For the retrived *Rattus rattus* sequences, we sought to determine if they were from a single individual, or from different specimens of black rat. To accomplish this, we generated callings for the *Rattus* mitochondria using GATK UnifiedGenotyper ^84^. Afterwards, we checked the heterozygosity ratio in biallelic positions with ≤ 5X depth, discarding those that could be originating from aDNA transition patterns (C↔T or G↔A).

### Uniparental markers and sex determination analysis

For human sequences, we called *rCRS* variants using *GATK* v.3.7 *UnifedGenotyper*, followed by creating a consensus *fasta* sequence with *bcftools* ^85^. We determined the mitochondrial haplogroup using haplogrep3 ^86^. To determine the genetic sex of the samples, we have used the ry_compute, considering ratio values below 0.016 as female, above 0.077 as male, and values in between as undeterminable ^87^.

For non-human animals, we computed the ratio of the normalised coverage in the X chromosome and the normalised coverage in the Y chromosome. Values above 1 are indicative of a male individual, while values close to 0 are indicative of a female.

### Rattus rattus dataset building and population genetics analysis

With the intention of studying the genetic background of the black rat sequences found in NMS022B, we started by creating a population genetics dataset. First, we downloaded and mapped publicly available data from *Rattus rattus* and *Rattus tanezumi* ^47,48,88,89^. For modern data, we trimmed the sequencing adapters, keeping sequences up to 30bp and merging paired end reads when necessary. Following this, we used *BWA mem* with default parameters to map the reads against the *Rattus rattus* reference genome *Rrattus_CSIRO.* Afterwards, duplicated sequences were removed by coordinate using *picard* and sequences with a mapping quality equal or above 30 were kept for downstream analysis. For ancient samples, we followed the same procedure, using *BWA backtrack* (-l 65536 -n -0.01 -o 2 -q 0) instead of the *mem* algorithm. Basic statistics were generated using Qualimap2 ^90^.

We selected a total of 52 individuals (including NMS022B) with more than 0.01X for population genetic analysis. First we generated genotype likelihoods using *angsd* ^91^ with the following options “*-uniqueOnly 1 -remove_bads 1 -trim 5 -C 50 -baq 1 -minInd 30 -nThreads 10 -skipTriallelic 1 -GL 2 -minMapQ 30 -doGlf 2 -doMajorMinor 1 -doMaf 2 -minMaf 0.05 - SNP_pval 1e-6”.* The resultant dataset contained 513,655 positions. We then computed a PCA using *PCAngsd* with 100,000 iterations ^92^. We also generated a dataset of pseudohaploid callings using *angsd* with the options “*-dohaplocall 1 -doCounts 1 -minMinor 0 -uniqueOnly 1 -remove_bads 1 -minMCQ 30 -minInd 40 -trim 5* ” resulting in 218,175 biallelic positions. We then runned the *f4* analysis, available in *AdmixtTools* as *qpDstat* ^49^. We tested the following tree (Rattus tanemuzi, NMS022B, Modern_Canada, X). Statistical significance was assessed using block jack-knife resampling.

### Rattus rattus mitochondrial phylogeny

We used GATK UnifiedGenotyper to generate callings for all published samples and NMS022B. For the mitochondrial phylogeny, we selected biallelic SNPs with a depth >=5X and a genotype calling quality of >=20. For heterozygous positions, we kept all positions with allele depth proportion of 1/9 as homozygous for the majority allele, discarding all the others in the process. Consensus mitochondrial sequences were generated for each sample with *bcftools,* using the filtered sets of SNPs and the *Rattus rattus* mitochondrial reference genome. To finalise the creation of the dataset, we masked in each sample those positions which did not meet the filtering criteria with an *N*.

A maximum-likelihood (ML) phylogenetic tree was created using RAxML ^93^. We used the GTR-GAMMA substitution model, with 1,000 bootstraps and *Rattus tanezumi* isolate RJPNAna02 as an outgroup. The resultant tree was visualised using *ggtree*.

### Human variant calling genetic imputation

Published human data from the Cherry Hinton site were mapped using *BWA bactkrack* as stated above ^50^. We then generated pseudohaploid callings of the published individuals and the filtered data of the sediment sample CHRY038B using *pileupcaller* (available at https://github.com/stschiff/sequenceTools) and the Human Origins (HO) dataset as calling panel ^94^. Dataset manipulation was done using PLINK ^95^.

We imputed genotypes in both CHRY038B (average genomic coverage 0,01x) and CHRY038 (average genomic coverage 0,06x) samples using QUILT, a tool developed to work with low-coverage whole-genome sequences ^96^, and the Haplotype Reference Consortium (HRC) reference panel ^53^. To speed up the process, each reference chromosome was split into the overlapping chunks (N=771 altogether), which were merged after imputation chromosome-wise using bcftools concat --ligate option ^85^. Post-imputation data processing included: a) correction of the conflicting genotypes (GT) by genotype probability (GP) (gp-to-get option in the bcftools tag2tag plugin); filtering out positions with minor allele frequency (MAF) less than 5% in the HRC reference panel. Additionally we kept only positions with genotype probability 98% and higher (GP >0,98); this filter was applied individually at each genome. This resulted in 24,469 and 224,460 positions in CHRY038B and CHRY038, respectively. (Supplementary Figure S29). As genotypes for both ancient samples were imputed from the genomes with sequence coverage (<0,1x), which is below the one indicated suitable for imputation ^97^, we consider our results as suggestive.

### Human population genetic analysis

A PCA using 442 modern European individuals from the HO dataset was computed using *smartpca,* with our samples CHRY038B and CHRY038 projected. We used options *shrinkmode: YES,* and *outliermode: 2.* Ellipses describing 95% confidence intervals for the projected samples were generated with *optionellconf: 0.95.* We repeated the process including CHRY038B and CHRY038 at different stages of imputation (Raw Pseudo-Haploid callings, GP>95 and GP>98).

We assessed relatedness (in this particular case, if CHRY038 and CHRY038B) were identical using READv2 with default parameters ^54^, with our samples at different stages of imputation and 20 additional samples from the Cherry Hinton site (Supplementary Table S11).

### Human phenotype prediction

Using the imputed genomes of CHRY038B and CHRY038 (the original sample), we extracted genotype calls for 39 of the 41 HIrisPlex-S ^98^ variants and 74 SNPs involved in diet and diseases, coding the allele information as the number of the effective allele (0, 1, 2) using plink (Supplementary Table S13A). The diet and disease set of 74 SNPs was selected starting from lists of variants previously analysed in aDNA studies ^99–101^, prioritising those with a role in the response to pathogenic infection. A table with the HIrisPlex-S SNP alleles per individual was uploaded on the HIrisPlex-S webtool (https://hirisplex.erasmusmc.nl/) to obtain probabilities values for eye, hair, and skin colour category. This output was then interpreted following the manual to obtain the final pigmentation prediction for the original skeletal element and for the genomes from soil (Supplementary Table S13B).

### Protein database search and result filtering

Raw mass spectrometry data were searched for each samples using *novor.cloud* ^102^ and *pFind* ^103,104^ against all proteins in SwissProt (version dated 2 may 2024) and cRAP databases (version dated of 4 march 2019) (https://www.thegpm.org/crap/). For both searches, fixed modification was set to include carbamidomethylation of cysteine and variable post-translational modifications (PTMs) were set to include proline hydroxylation, glutamine and asparagine deamidation, methionine oxidation, and pyroglutamate formation from glutamine and glutamic acid. Searches were conducted with trypsin full-specific digestion. Precursor mass tolerance was set to 15 ppm and fragment mass tolerance to 0.02 Da and the false-discovery rate of peptide spectrum matches equal ≤1.0%. *pFind* peptides were filtered based on their score. Only peptides with a score below or equal to 0.01 and at least two PSMs were considered. Protein matches were considered if they had at least two unique peptide matches in either *novor.cloud* or *pFind*. All contaminants from our samples were identified and removed using the peptide identified within our extraction blank and the cRAP database.

### Post-translational modifications analysis

For both *novor.cloud* and *pFind*, we measured the proportion of all identified post-translational modifications (PTMs) relative to the total number of amino acids. We quantified in particular methionine oxidation and glutamine/asparagine deamidation along with the peptide length distributions

### Rattus rattus peptide analysis

For the search of rat peptides in the NMS022B sample, we built an in-house database with protein sequences obtained from the reference genome of *Rattus rattus*_CSIRO and *Rattus norvegicus*_GRCr8 (The SwissProt database contains only reference protein sequences of *R. norvegicus*). Next, all filtered peptides were searched against this custom database and NCBI_nr using the same parameters as before. In order to identify specific-species peptides, we blasted all the identified peptides using BLASTP against the ncbi nr database ^105^. Filtered results (100% of identity and coverage) were parsed on MEGAN. We used species-specific sequences from ollagens, lactadherin, and glycoprotein proteins to assign the NMS022B sample peptides within the genera of European *Rattus*.

## Availability of data and material

The datasets generated and analysed during the current study are available in the ENA and ProteomeXchange. Genomic data from the original skeletal elements (previously published) is available in the ENA repository under the accession ID: PRJEB59976. and the data depository of the EBC (https://evolbio.ut.ee/). All data needed to evaluate the conclusions in the paper are present in the paper and/or the Supplementary Materials.

## Supporting information

Supplementary Methods Figures S1-S29 and Table description

## Acknowledgements

We thank Sarah A. Inskip and Ruoyun Hui for their insightful comments and suggestions. This work was supported by the European Union through the European Research Council Advanced Grant “Making Ancestors: the Politics of Death in European Prehistory” (No. 885137) (T.d.D.M, B.B., J.E.R., C.L.S.), the Wellcome Trust (Award no. 2000368/Z/15/Z) (S.A.I., J.D., C.C., J.E.R, C.L.S.), the European Regional Development Fund (Project No. 2014- 2020.4.01.16-0030) (C.L.S.), and by the Estonian Research Council grant (PSG492) (E.O.). We thank the support of Sergo Kasvandik and the University of Tartu Proteomics Core Facility, The Genotyping and Sequencing Core laboratory of EGCUT.

## Author Information

### Authors’ contributions

Conceptualization: CLS, TdDM Data curation: TdDM, AS

Formal Analysis: TdDM, BB, RB, LK, EDA, CLS Funding acquisition: JER, CW, EO

Investigation: CLS

Methodology: TdDM, BB, RB, CLS Project administration: CLS Resources: SAI, JD, CC, JER Supervision: CLS, CW Visualisation: TdDM, BB, RB

Writing – original draft: TdDM, BB, RB, CLS Writing – review & editing: All authors

## Ethics declarations

### Competing interests

The authors declare no competing interests.

## Notes

### Competing Interest Statement

The authors have declared no competing interest.

## References

1. Slon, V. et al. Neandertal and Denisovan DNA from Pleistocene sediments. Science 356, 605–608 (2017).

2. Zhang, D. et al. Denisovan DNA in Late Pleistocene sediments from Baishiya Karst Cave on the Tibetan Plateau. Science 370, 584–587 (2020).

3. Zavala, E. I. et al. Pleistocene sediment DNA reveals hominin and faunal turnovers at Denisova Cave. Nature 595, 399–403 (2021).

4. Massilani, D. et al. Microstratigraphic preservation of ancient faunal and hominin DNA in Pleistocene cave sediments. Proc. Natl. Acad. Sci. U. S. A. 119, (2022).

5. Pedersen, M. W. et al. Postglacial viability and colonization in North America’s ice-free corridor. Nature 537, 45–49 (2016).

6. Pedersen, M. W. et al. A comparative study of ancient environmental DNA to pollen and macrofossils from lake sediments reveals taxonomic overlap and additional plant taxa. Quat. Sci. Rev. 75, 161–168 (2013).

7. Alsos, I. G. et al. Sedimentary ancient DNA from Lake Skartjørna, Svalbard: Assessing the resilience of arctic flora to Holocene climate change. Holocene 26, 627–642 (2016).

8. Kjær, K. H. et al. A 2-million-year-old ecosystem in Greenland uncovered by environmental DNA. Nature 612, 283–291 (2022).

9. Gelabert, P. et al. Genome-scale sequencing and analysis of human, wolf, and bison DNA from 25,000-year-old sediment. Curr. Biol. 31, 3564–3574.e9 (2021).

10. Wang, Y. et al. Late Quaternary dynamics of Arctic biota from ancient environmental genomics. Nature 600, 86–92 (2021).

11. Fernandez-Guerra, A. et al. A 2-million-year-old microbial and viral communities from the Kap København Formation in North Greenland. bioRxiv 2023.06.10.544454 (2023) doi:10.1101/2023.06.10.544454.

12. Armbrecht, L. et al. Ancient marine sediment DNA reveals diatom transition in Antarctica. Nat. Commun. 13, 1–14 (2022).

13. Pérez, V., Liu, Y., Hengst, M. B. & Weyrich, L. S. A Case Study for the Recovery of Authentic Microbial Ancient DNA from Soil Samples. Microorganisms 10, (2022).

14. Dabney, J., Meyer, M. & Pääbo, S. Ancient DNA damage. Cold Spring Harb. Perspect. Biol. 5, (2013).

15. Slon, V. et al. Extended longevity of DNA preservation in Levantine Paleolithic sediments, Sefunim Cave, Israel. Sci. Rep. 12, 1–16 (2022).

16. Le Meillour, L. et al. Identification of degraded bone and tooth splinters from arid environments using palaeoproteomics. Palaeogeogr. Palaeoclimatol. Palaeoecol. 511, 472–482 (2018).

17. Hendy, J. et al. A guide to ancient protein studies. Nat Ecol Evol 2, 791–799 (2018).

18. Demarchi, B. et al. Protein sequences bound to mineral surfaces persist into deep time. Elife 5, (2016).

19. Welker, F. et al. Enamel proteome shows that Gigantopithecus was an early diverging pongine. *Nature* 576, 262–265 (2019).

20. Cappellini, E. et al. Early Pleistocene enamel proteome from Dmanisi resolves Stephanorhinus phylogeny. Nature 574, 103–107 (2019).

21. Welker, F. et al. The dental proteome of Homo antecessor. Nature 580, 235–238 (2020).

22. Chen, F. et al. A late Middle Pleistocene Denisovan mandible from the Tibetan Plateau. Nature 569, 409–412 (2019).

23. Horn, I. R. et al. Palaeoproteomics of bird bones for taxonomic classification. Zool. J. Linn. Soc. 186, 650–665 (2019).

24. Buckley, M. Paleoproteomics: An Introduction to the Analysis of Ancient Proteins by Soft Ionisation Mass Spectrometry. in Paleogenomics: Genome-Scale Analysis of Ancient DNA (eds. Lindqvist, C. & Rajora, O. P.) 31–52 (Springer International Publishing, Cham, 2019).

25. Rao, H. et al. Palaeoproteomic analysis of Pleistocene cave hyenas from east Asia. Sci. Rep. 10, 16674 (2020).

26. Bleasdale, M. et al. Ancient proteins provide evidence of dairy consumption in eastern Africa. Nat. Commun. 12, 632 (2021).

27. Runge, A. K. W. et al. Palaeoproteomic analyses of dog palaeofaeces reveal a preserved dietary and host digestive proteome. Proc. Biol. Sci. 288, 20210020 (2021).

28. Kanerva, S., Smolander, A., Kitunen, V., Ketola, R. A. & Kotiaho, T. Comparison of extractants and applicability of MALDI–TOF-MS in the analysis of soil proteinaceous material from different types of soil. Org. Geochem. 56, 1–9 (2013).

29. Solazzo, C., Scibè, C. & Eng-Wilmot, K. Proteomics characterization of‘ organic’ metal threads- First results and future directions. *England*, May 13–18, *2019 …* (2019).

30. Giuffrida, M. G., Mazzoli, R. & Pessione, E. Back to the past: deciphering cultural heritage secrets by protein identification. Appl. Microbiol. Biotechnol. 102, 5445–5455 (2018).

31. Li, L., Zhu, L. & Xie, Y. Proteomics analysis of the soil textile imprints from tomb M6043 of the Dahekou Cemetery site in Yicheng County, Shanxi Province, China. Archaeol. Anthropol. Sci. 13, 7 (2021).

32. Damgaard, P. B. et al. Improving access to endogenous DNA in ancient bones and teeth. Sci. Rep. 5, 11184 (2015).

33. Pinhasi, R., Fernandes, D. M., Sirak, K. & Cheronet, O. Isolating the human cochlea to generate bone powder for ancient DNA analysis. Nat. Protoc. 14, 1194–1205 (2019).

34. Kazarina, A. et al. Analysis of the bacterial communities in ancient human bones and burial soil samples: Tracing the impact of environmental bacteria. J. Archaeol. Sci. 109, 104989 (2019).

35. Wood, D. E., Lu, J. & Langmead, B. Improved metagenomic analysis with Kraken 2. Genome Biol. 20, 257 (2019).

36. Wilkin, S., et al. SP3 (Single-Pot, Solid-Phase, Sample-Preperation) Protein Extraction for Dental Calculus. (2021).

37. Cleland, T. P. Human Bone Paleoproteomics Utilizing the Single-Pot, Solid-Phase-Enhanced Sample Preparation Method to Maximize Detected Proteins and Reduce Humics. J. Proteome Res. 17, 3976–3983 (2018).

38. Boutet, E., Lieberherr, D., Tognolli, M., Schneider, M. & Bairoch, A. UniProtKB/Swiss-Prot. Methods Mol. Biol. 406, 89–112 (2007).

39. Breitwieser, F. P., Baker, D. N. & Salzberg, S. L. KrakenUniq: confident and fast metagenomics classification using unique k -mer counts. Genome Biol. 19, 1–10 (2018).

40. Pockrandt, C., Zimin, A. V. & Salzberg, S. L. Metagenomic classification with KrakenUniq on low-memory computers. bioRxiv 2022.06.01.494344 (2022) doi:10.1101/2022.06.01.494344.

41. McGhee, J. J. et al. Meta-SourceTracker: application of Bayesian source tracking to shotgun metagenomics. PeerJ 8, e8783 (2020).

42. Tisza, M. J. & Buck, C. B. A catalog of tens of thousands of viruses from human metagenomes reveals hidden associations with chronic diseases. Proceedings of the National Academy of Sciences 118, e2023202118 (2021).

43. Stewart, R. D. et al. Compendium of 4,941 rumen metagenome-assembled genomes for rumen microbiome biology and enzyme discovery. Nat. Biotechnol. 37, 953–961 (2019).

44. Su, M. et al. Metagenomic Analysis Revealed Differences in Composition and Function Between Liquid-Associated and Solid-Associated Microorganisms of Sheep Rumen. Front. Microbiol. 13, 851567 (2022).

45. Rampelli, S. et al. Metagenome Sequencing of the Hadza Hunter-Gatherer Gut Microbiota. Curr. Biol. 25, 1682–1693 (2015).

46. O’Leary, N. A. et al. Reference sequence (RefSeq) database at NCBI: current status, taxonomic expansion, and functional annotation. Nucleic Acids Res. 44, D733–45 (2016).

47. Yu, H. et al. Palaeogenomic analysis of black rat (Rattus rattus) reveals multiple European introductions associated with human economic history. Nat. Commun. 13, 2399 (2022).

48. Puckett, E. E. et al. Global population divergence and admixture of the brown rat (Rattus norvegicus). Proc. Biol. Sci. 283, (2016).

49. Patterson, N. et al. Ancient admixture in human history. Genetics 192, 1065–1093 (2012).

50. Hui, R. et al. Genetic history of Cambridgeshire before and after the Black Death. Sci Adv 10, eadi5903 (2024).

51. Scheib, C. L. et al. Local population structure in Cambridgeshire during the Roman occupation. bioRxiv 2023.07.31.551265 (2023) doi:10.1101/2023.07.31.551265.

52. Lazaridis, I. et al. Genomic insights into the origin of farming in the ancient Near East. Nature 536, 419–424 (2016).

53. McCarthy, S. et al. A reference panel of 64,976 haplotypes for genotype imputation. Nat. Genet. 48, 1279–1283 (2016).

54. Alaçamlı, E. et al. READv2: Advanced and user-friendly detection of biological relatedness in archaeogenomics. bioRxiv 2024.01.23.576660 (2024) doi:10.1101/2024.01.23.576660.

55. Wibowo, M. C. et al. Reconstruction of ancient microbial genomes from the human gut. Nature 594, 234–239 (2021).

56. Payen, G., Rimoux, L., Gueux, M. & Lery, N. Body putrefaction in ‘air tight’ burials. IV. Microbiologic findings and environment. Acta Med. Leg. Soc. 38, 153–163 (1988).

57. Anastasiou, E., Papathanasiou, A., Schepartz, L. A. & Mitchell, P. D. Infectious disease in the ancient Aegean: Intestinal parasitic worms in the Neolithic to Roman Period inhabitants of Kea, Greece. Journal of Archaeological Science: Reports 17, 860–864 (2018).

58. Gildner, T. E. & Casana, J. Intestinal parasitic infection within a wealthy nineteenth century household from rural New England: Evidence from Dartmouth College, New Hampshire. Journal of Archaeological Science: Reports 37, 102990 (2021).

59. Cessford, C. et al. Beyond Plague Pits: Using Genetics to Identify Responses to Plague in Medieval Cambridgeshire. European Journal of Archaeology 1–23 (2021).

60. Spyrou, M. A. et al. Phylogeography of the second plague pandemic revealed through analysis of historical Yersinia pestis genomes. Nature Communications vol. 10 Preprint at 10.1038/s41467-019-12154-0 (2019).

61. Cessford, C. et al. Buried with their Buckles On: Clothed Burial at the Augustinian Friary, Cambridge. Mediev. Archaeol. 66, 151–187 (2022).

62. Cessford, C., Samuel, M., Herring, V., Holder, N. & Mills, P. THE ARCHITECTURE OF THE AUGUSTINIAN FRIARY, CAMBRIDGE. The Antiquaries Journal 1–33 (2023).

63. Cessford, C. & Neil, B. The people of the Cambridge Austin friars. Archaeological Journal 179, 385–446 (2022).

64. Hedrich, H. Taxonomy and Stocks and Strains. The Laboratory Rat (2020) doi:10.1016/B978-012074903-4/50006-6.

65. Morozova, I. et al. New ancient Eastern European Yersinia pestis genomes illuminate the dispersal of plague in Europe. Philos. Trans. R. Soc. Lond. B Biol. Sci. 375, 20190569 (2020).

66. Villalba-Mouco, V. et al. A 23,000-year-old southern Iberian individual links human groups that lived in Western Europe before and after the Last Glacial Maximum. Nat Ecol Evol 7, 597–609 (2023).

67. Keller, M. & Scheib, C. L. Ancient DNA extract purification (chunk samples/high volume). (2023).

68. Keller, M., Scheib, C. L. & Bonucci, B. Library preparation (dsDNA double indexing, non-UDG, 2x split). (2023).

69. Schubert, M., Lindgreen, S. & Orlando, L. AdapterRemoval v2: rapid adapter trimming, identification, and read merging. BMC Res. Notes 9, 88 (2016).

70. Schmieder, R. & Edwards, R. Quality control and preprocessing of metagenomic datasets. Bioinformatics 27, 863–864 (2011).

71. Bushnell, B. BBMap. Preprint at (2015).

72. Li, H. & Durbin, R. Fast and accurate short read alignment with Burrows-Wheeler transform. Bioinformatics 25, 1754–1760 (2009).

73. Schubert, M. et al. Improving ancient DNA read mapping against modern reference genomes. BMC Genomics 13, 178 (2012).

74. Guellil, M. et al. Genomic blueprint of a relapsing fever pathogen in 15th century Scandinavia. Proc. Natl. Acad. Sci. U. S. A. 115, 10422–10427 (2018).

75. Knights, D. et al. Bayesian community-wide culture-independent microbial source tracking. Nat. Methods 8, 761–763 (2011).

76. Dixon, P. VEGAN, a package of R functions for community ecology. J. Veg. Sci. 14, 927–930 (2003).

77. Lu, J. et al. Metagenome analysis using the Kraken software suite. Nat. Protoc. 17, 2815–2839 (2022).

78. Jónsson, H., Ginolhac, A., Schubert, M., Johnson, P. L. F. & Orlando, L. mapDamage2.0: fast approximate Bayesian estimates of ancient DNA damage parameters. Bioinformatics 29, 1682– 1684 (2013).

79. van den Boogaart, K. G. & Tolosana-Delgado, R. ‘compositions’: A unified R package to analyze compositional data. Comput. Geosci. 34, 320–338 (2008).

80. Rohart, F., Gautier, B., Singh, A. & Lê Cao, K.-A. mixOmics: An R package for ’omics feature selection and multiple data integration. PLoS Comput. Biol. 13, e1005752 (2017).

81. Chen, Y., Ye, W., Zhang, Y. & Xu, Y. High speed BLASTN: an accelerated MegaBLAST search tool. Nucleic Acids Res. 43, 7762–7768 (2015).

82. Beier, S., Tappu, R. & Huson, D. H. Functional Analysis in Metagenomics Using MEGAN 6. in Functional Metagenomics: Tools and Applications (eds. Charles, T. C., Liles, M. R. & Sessitsch, A.) 65–74 (Springer International Publishing, Cham, 2017).

83. Renaud, G., Slon, V., Duggan, A. T. & Kelso, J. Schmutzi : contamination estimate and endogenous mitochondrial consensus calling for ancient DNA. Additional le 1. (2015).

84. McKenna, A. et al. The Genome Analysis Toolkit: a MapReduce framework for analyzing next-generation DNA sequencing data. Genome Res. 20, 1297–1303 (2010).

85. Danecek, P. et al. Twelve years of SAMtools and BCFtools. Gigascience 10, (2021).

86. Schönherr, S., Weissensteiner, H., Kronenberg, F. & Forer, L. Haplogrep 3 - an interactive haplogroup classification and analysis platform. Nucleic Acids Res. 51, W263–W268 (2023).

87. Skoglund, P., Storå, J., Götherström, A. & Jakobsson, M. Accurate sex identification of ancient human remains using DNA shotgun sequencing. J. Archaeol. Sci. 40, 4477–4482 (2013).

88. Massini Espino, M., Mychajliw, A. M., Almonte, J. N., Allentoft, M. E. & Van Dam, A. R. Raptor roosts as invasion archives: insights from the first black rat mitochondrial genome sequenced from the Caribbean. Biol. Invasions 24, 17–25 (2022).

89. Teng, H. et al. Whole-Genome Sequencing Reveals Genetic Variation in the Asian House Rat. *G3* 6, 1969–1977 (2016).

90. Okonechnikov, K., Conesa, A. & García-Alcalde, F. Qualimap 2: advanced multi-sample quality control for high-throughput sequencing data. Bioinformatics 32, 292–294 (2016).

91. Korneliussen, T. S., Albrechtsen, A. & Nielsen, R. ANGSD: Analysis of Next Generation Sequencing Data. BMC Bioinformatics 15, 356 (2014).

92. Meisner, J. & Albrechtsen, A. Inferring Population Structure and Admixture Proportions in Low-Depth NGS Data. Genetics 210, 719–731 (2018).

93. Stamatakis, A. RAxML version 8: a tool for phylogenetic analysis and post-analysis of large phylogenies. Bioinformatics 30, 1312–1313 (2014).

94. Lazaridis, I. et al. Ancient human genomes suggest three ancestral populations for present-day Europeans. Nature 513, 409–413 (2014).

95. Purcell, S. et al. PLINK: a tool set for whole-genome association and population-based linkage analyses. Am. J. Hum. Genet. 81, 559–575 (2007).

96. Davies, R. W. et al. Rapid genotype imputation from sequence with reference panels. Nat. Genet. 53, 1104–1111 (2021).

97. Hui, R., D’Atanasio, E., Cassidy, L. M., Scheib, C. L. & Kivisild, T. Evaluating genotype imputation pipeline for ultra-low coverage ancient genomes. Sci. Rep. 10, 18542 (2020).

98. Chaitanya, L. et al. The HIrisPlex-S system for eye, hair and skin colour prediction from DNA: Introduction and forensic developmental validation. Forensic Sci. Int. Genet. 35, 123–135 (2018).

99. Allentoft, M. E. et al. Population genomics of Bronze Age Eurasia. Nature 522, 167–172 (2015).

100. Günther, T. et al. Population genomics of Mesolithic Scandinavia: Investigating early postglacial migration routes and high-latitude adaptation. PLoS Biol. 16, e2003703 (2018).

101. Olalde, I. et al. Derived immune and ancestral pigmentation alleles in a 7,000-year-old Mesolithic European. Nature 507, 225–228 (2014).

102. novor.cloud. https://app.novor.cloud/ https://app.novor.cloud/ (2024).

103. Shao, G. et al. How to use open-pFind in deep proteomics data analysis?- A protocol for rigorous identification and quantitation of peptides and proteins from mass spectrometry data. Biophys Rep 7, 207–226 (2021).

104. Chi, H. et al. Comprehensive identification of peptides in tandem mass spectra using an efficient open search engine. Nat. Biotechnol. (2018) doi:10.1038/nbt.4236.

105. Altschul, S. F., Gish, W., Miller, W., Myers, E. W. & Lipman, D. J. Basic local alignment search tool. J. Mol. Biol. 215, 403–410 (1990).

