## Supplementary Methods Figures S1-S29 and Table description for "Bone adhered soil as a source of target and environmental DNA and proteins"

#### S1 Microbial analysis

##### S1.1 Reference dataset creation

We downloaded raw fastq sequences from ENA ^1–9^ (Supplementary Table S16). Adapter sequences were trimmed and read pairs were merged using AdapterRemoval2 , with a minimum length of 30, minimum overlap of 11 and minimum sequencing quality of 20. Low complexity sequences using prinseq and a dust value of 7 ^10^ and duplicated reads were collapsed using bbmap ^11^. Deduplicated and high quality sequences were mapped against the Human Reference Genome 37 using BWA mem ^12^. Human free sequences were then classified against a Microbial Database (MicrobialDB as for 16/08/2020) using KrakenUniq ^13,14^. Raw KrakenUniq were filtered using the Evalue formula described by Borry and Guellil ^15,16^. We accepted a minimum Evalue of 5 and 200 sequences as minimum values to consider a hit as true down to the taxonomic level of species.

After the initial bioinformatic processing of the individual samples, we proceeded to merge all the filtered reports in one unique dataset. To achieve this we use a custom python script, discarding all species that were represented with under the 0.02% of the total sequences of the whole dataset.

To test the behaviour of this reference dataset, we used Sourcetracker2 strategy *leave-one-out* (loo) ^17^*.* In brief, this feature iteratively compares each different sample in the dataset against all the samples (Suppl Figure S24). We further validate these results by performing a hierarchical cluster analysis of the Bray Curtis distances of the samples’ microbial abundances using R package *cluster* *^18^* (Suppl Figure S25). We then proceed to discard the following samples due to their low levels of similitude to other samples in their clusters: *skin25, CowRumen-10678_0001, CowRumen-10678_0006, CowRumen-10678_0007,* and *CowRumen-10678_0013.*

Once the source dataset was properly delimited, we generated a set of source-specific species. Those were characterised using Sourcetracker2 option “*per_sink_feature_assignments*”, and were estimated to human gut (HuGt=49; ruralGut and urbanGut), ruminant gut (RmGt=10; CowRumen and Sheep Rumen), skin (Skin=45), oral (Oral=64; ModernCalculus, Subplaque and Supplaque), and soil (Soil=14; Soil and FarmlandSoil) (Supplementary Table S2). Although for the most part, species are unique to a source, some level of species overlap is found between human gut and rumen (HuRm=10), soil and skin (SlSk=9), and oral and skin (OrSk=12) (Supplementary Figure S26).

##### S1.2 GC content analysis

Analysis of the GC content of each source reveals a weak tendency for GC bias in short human gut and soil microbial sequences (HuGt, HuRm, Soil, SlSk p < 0.01), but not in oral, ruminant gut, or skin microbiomes (RmGt p= 0.375; Oral p = 0.629; OrSk p = 0.316; Skin p = 0.176) (Supplementary Figure S27).

##### S1.3 Oral Microbiome analysis

We extracted species corresponding to different microbial datasets samples CHR038B, CHR051B and TRM003B raw krakenUniq reports. Two different datasets were used for PCA analyses, one including the Oral Microbiome inferred by Sourcetracker2 (76 oral species), and the *Human Oral Microbiome Database* (HOMD) ^19^. We also included 68 published ancient and modern oral metagenomes. For each metagenome, species were extracted from the raw KrakenUniq report if they had an Evaule >=7, and at least 10 reads. After this initial filtering, we merged the filtered metagenomes for each of the 2 species datasets. Once merged, we normalised each sample in each for library size, and discarded species with representation below 0.02%. The resultant datasets were then renormalised using a *clr* transformation in the R package *transformations* *^20^*, and a Principle Component Analysis (PCA) was computed using R package *mixOmics* *^21^**.* Resultant sample values and loadings were visualised using R. . The results show that our samples cluster with other ancient published oral metagenomes ^22^, but closer to modern plaque than modern calculus ^8,9,22^. In particular, they fall with other samples containing low microbial species coverage, indicating that this factor could bias the results (Supplementary Figure S28).

#### S2 Eukaryotes Screening

##### S2.1 KrakenUniq dataset creation

We downloaded all refseq sequences corresponding to plastids and mitochondria as of 02/11/2022. Additionally we added the whole Human Reference Genome 38 and the contaminants database using KrakenUniq built-in function “*--download-library*”. We then proceed to build the KrakenUniq database using default settings (*--kmer-len 35, --minimizer-len 31*). This operation was run using 500GB of RAM memory and 32 cpus.

In addition to that, we also build a set of reference genomes for each different genus and species. To do that we selected all the representative genomes for each different species according to NCBI. We then created the individual reference genome for the species, and proceeded to obtain taxonomic information using taxonomy_ranks available in ETE toolkit ^23^. We then concatenated all species genomes inside a genus into a fasta, and then converted into a reference.

##### S2.2 Screening

##### We ran the preprocessed data (see Methods) against the aforementioned custom Organelle KrakenUniq database. In order to validate the hits against animal mitochondria we used the E-value formula described by Guellil and Borry on the raw KrakenUniq reports. In this case, we used an E-value of 7 at a genus level and a minimum of 3 reads per hit. The reasoning behind this is that three reads matches a read length of 50bp, representing around 1% of the mitochondrial genome for an animal. We used sheep bone samples Z1460, Z1461,Z1463, and Z1464 as controls (Supplementary Figure S9).

Once target species were selected, we created a reference by concatenating all species found inside the target genus into a unique reference. This step was purely done to speed up downstream analysis. We mapped the preprocessed fastq against this multi-genus reference genome with the common ancient DNA ^24^ mapping settings in BWA *backtrack* *^25^*; edit distance of 0.01 (-e 0.01), gap open penalty of 2 (-o 2) and seeding disabled (-l 10,000). All reads with a mapping quality of 0 or higher were converted back into fastq using the command “*samtools fastq”**^26^**,* keeping the original qualities found in the read*.* Afterwards, we started a set of parallel mappings, using those filtered fastq, and all the different genus references. We reclassified mapping reads according to lowest Edit Distance to each genus, discarding shared reads with equal edit distance in the process, using a custom-build python script. This was followed by a step of BAM to fastq reconversion by genus, and a second step of mapping, this time for individual species inside the analysed genus. Reads were reclassified by Edit Distance, obtaining individual BAMs for each different species. Basic statistics were performed for those BAMs using Qualimap2 ^27^. We then selected the best candidates for species by the number of mapped reads, and in cases of an equal number of mapped sequences, according to average edit distance. Finally, we mapped the original pre-processed fastq against each different individual target reference genome and reclassified the sequences (Supplementary Figure S10). We generated basic mapping statistics with Qualimap2 and aDNA damage patterns with MapDamage2 ^28^. We also calculated the -Δ statistic (differential of edit distance) ^29^ (Supplementary Figure S13). We assumed a species was present if at least 3 reads mapped to the species. Whole genome references were then created for those species to map against.

The reasoning behind this initial screening and iterative steps of mapping and reclassification is to overcome the computational limitations of the cluster. Metagenomic classification software such as Kraken2 or Centrifuge, despite being extremely fast and relatively accurate, do not allow for proper result validation ^30,31^. KrakenUniq alone could be a good candidate ^13^, unfortunately even with new memory saving settings present in the latest release of the software ^14^, building a database including all whole genomes from Eukaryotes requires a substantial amount of memory. Finally, MALT needs a prohibitive amount of RAM memory to run ^29^. We have also tested *euka* *^32^**,* specifically designed for organelles, but the taxonomic resolution is somewhat lower and the running time it also longer. Following the pipeline above we manage to run our samples (10-20 million reads), in less than 2 hours using only 20GB of RAM memory and 8 cpus. All directories and files associated with this pipeline including KrakenUniq database, reference genomes and metadata files have an approximate disk space of less than 3GB.

##### S2.3 Whole genome mapping and quality control

We downloaded whole genomes for the target species and created references from them. We mapped using *BWA backtrack* as previously specified and reclassified the resultant mapped reads according to their edit distance, in order to avoid spurious mappings between evolutionary close organisms (Supplementary Figure S23). To finalise the authentication process, we validated the resultant sequences according to the presence of aDNA deamination patterns, read length distribution, edit distance distribution decay, and normalised coverage distribution across the genome.

### Supplementary Figures

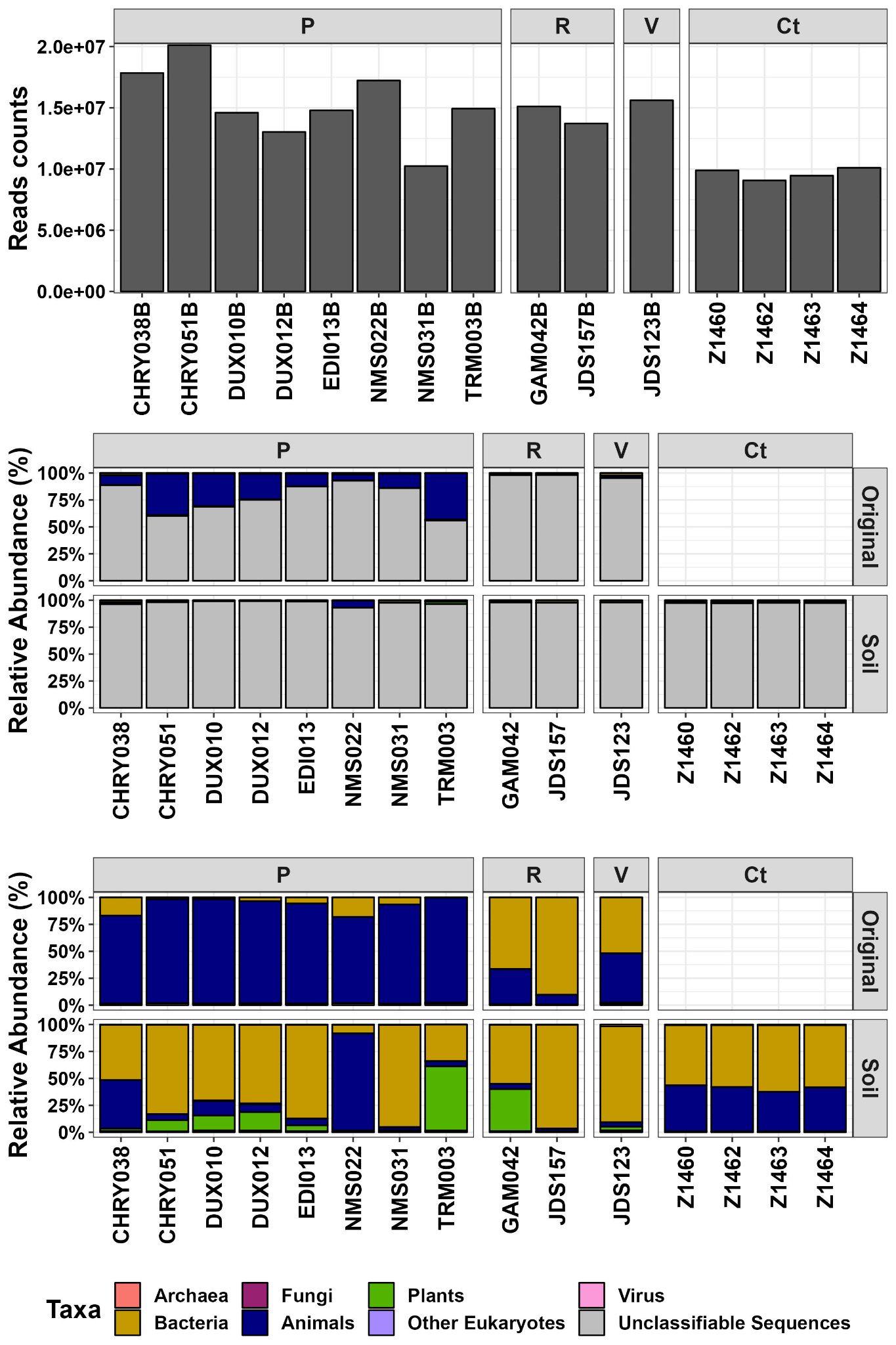

**Figure S1.** **Sequences’ abundances.** Barplot displaying the number of generated sequences per sample (A). Stacked plot of the relative abundance of sequences assigned by Kraken2 to the different domains of life (Archaea, Bacteria, Eukaryota [Fungi, Animals, Plants, Other], Virus), with (B) and without (C) taking into account unclassifiable sequences for both Soil and Original Skeletal element. Samples are grouped by their sampling origin ( P = Petrous Bone Soil, R = Rib Soil, V = Vertebrae Soil, Ct = Control).

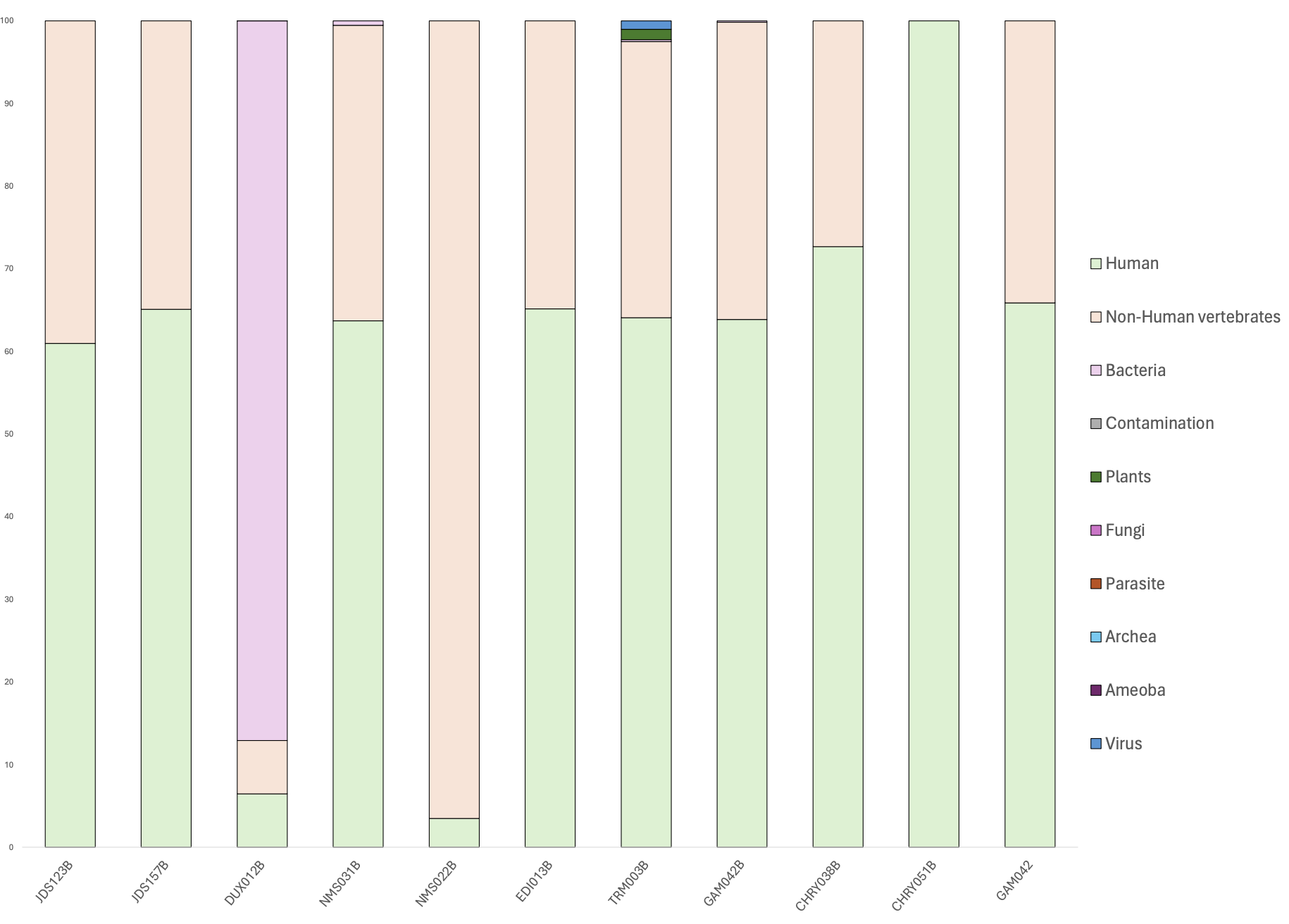

**Figure S2. Relative proportions of peptide origin.** Stacked bar plot showed the relative abundance for peptide origin based on novor.cloud output.

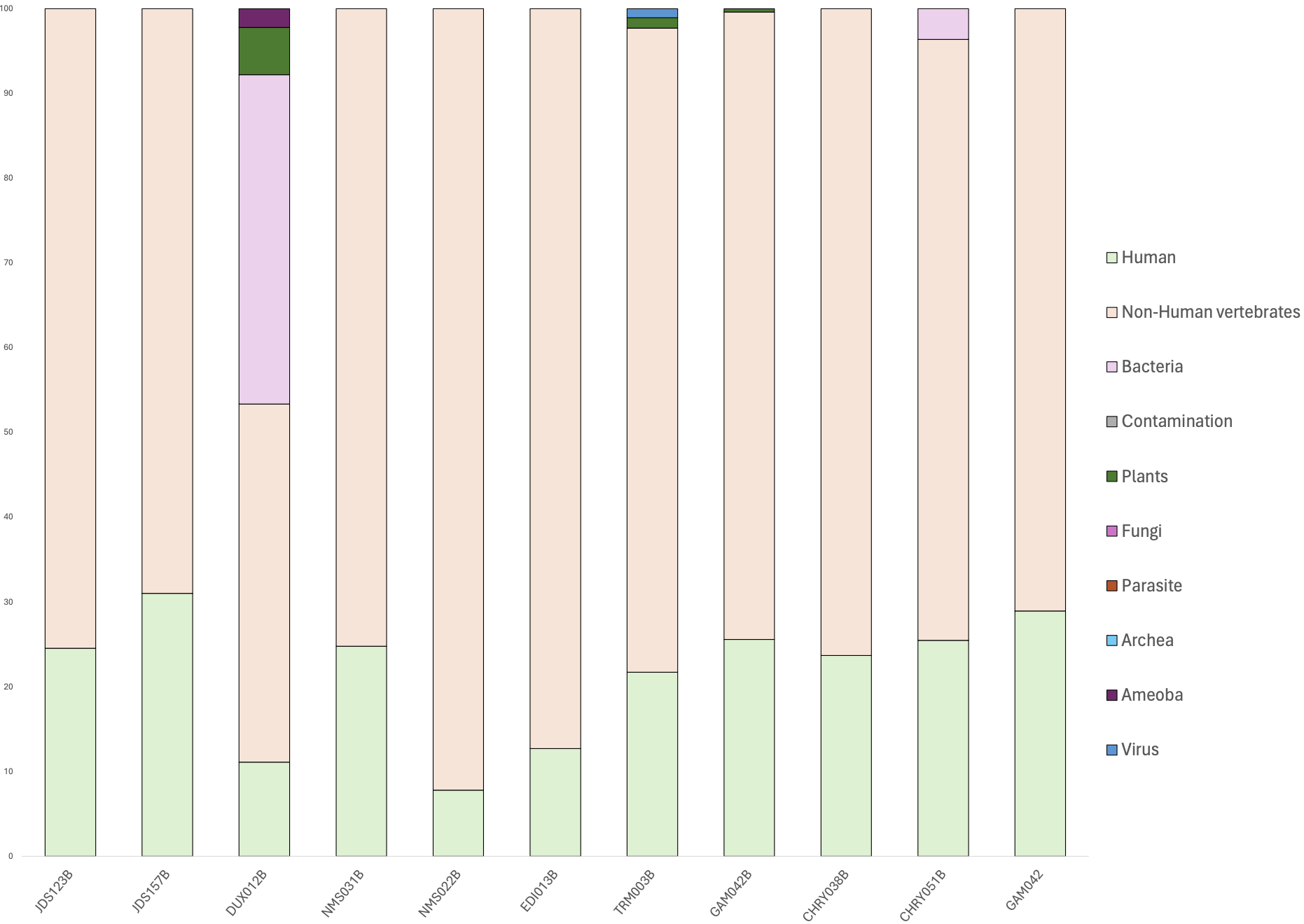

**Figure S3. Relative proportions of peptide origin.** Stacked bar plot showed the relative abundance for peptide origin based on pFind output.

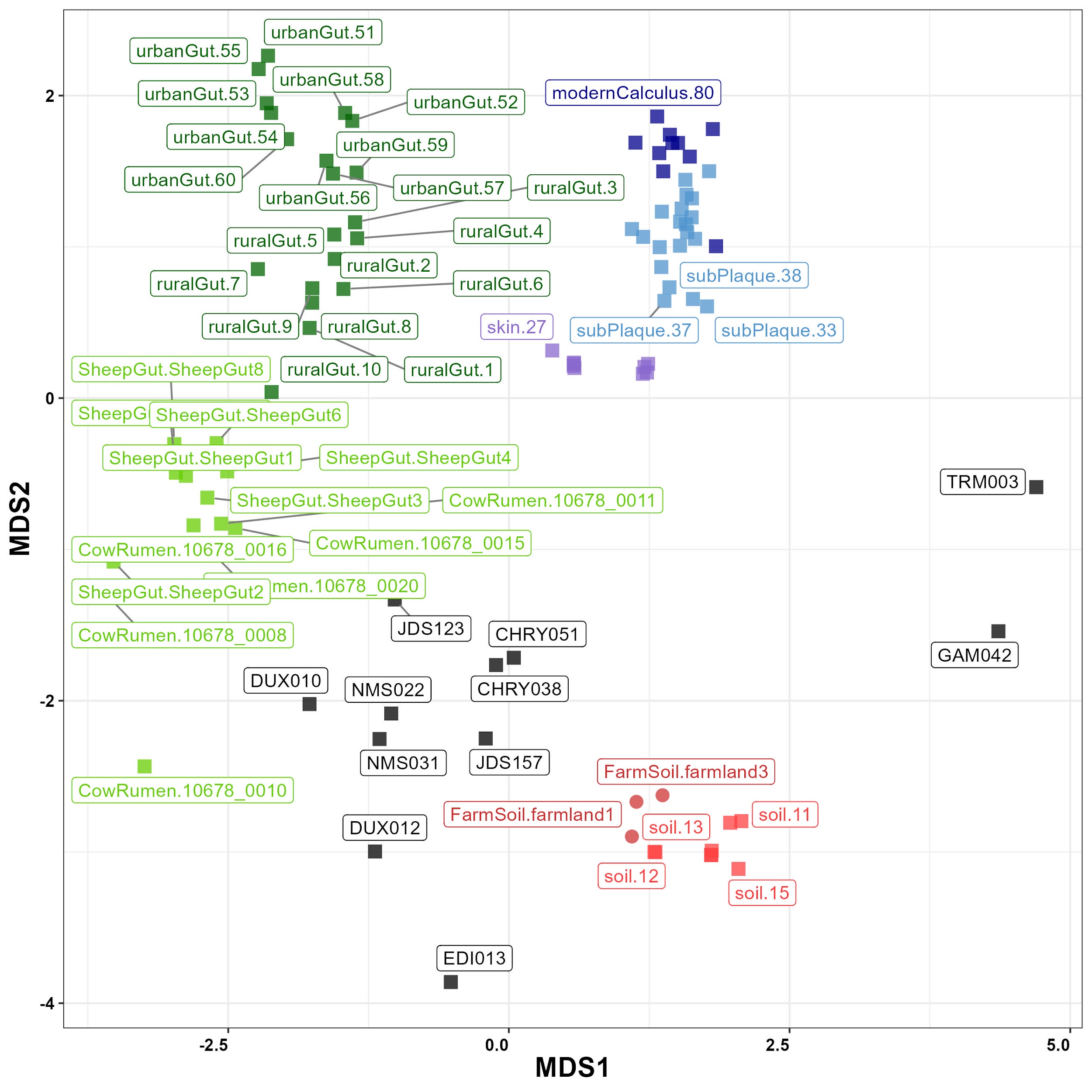

### Figure S4. Non-metric dimensional scaling plot using Bray-Curtis distances of analysed ancient soil samples and a set of modern references metagenomic profiles obtained with KrakenUniq. Modern microbiome references include soil (Soil), farmland soil (FarmSoil), human non-industrialised gut microbiome (ruralGut), human industrialised gut microbiome (urbanGut), ruminant gut microbiome (CowRumen and SheepRumen), human skin microbiome (skin), human calculus (modernCalculus), human supragingival plaque (supPlaque) and human subgingival plaque (subPlaque).

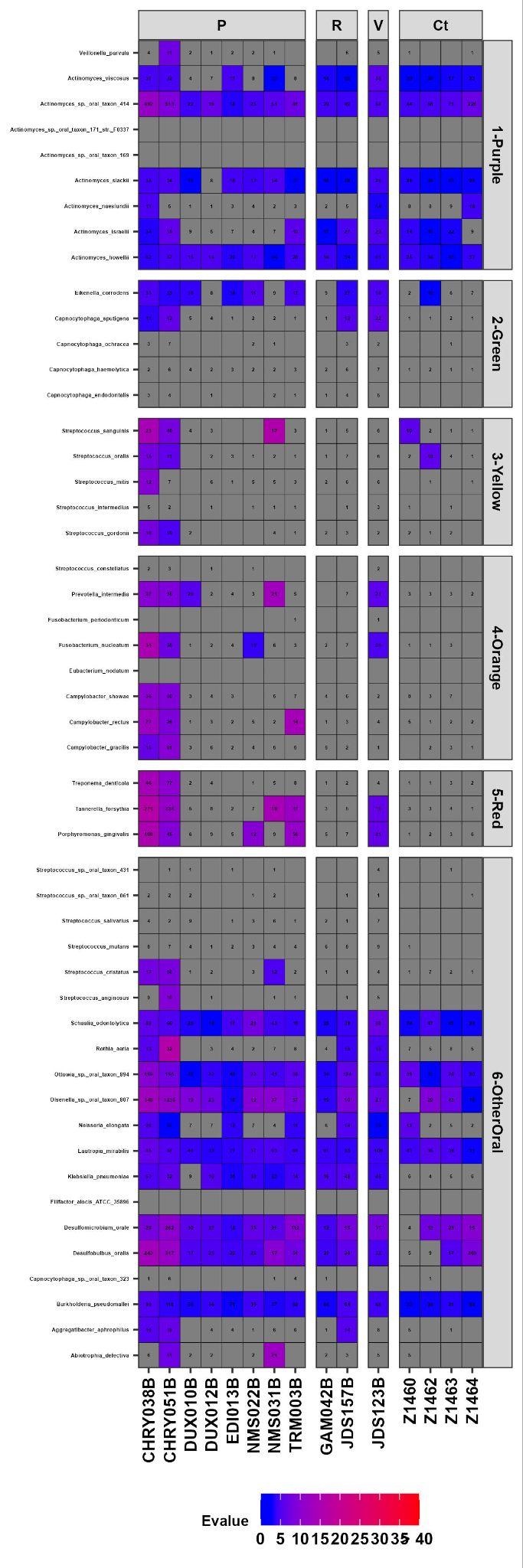

**Figure S5.** Oral Bacteria presence by complex (Purple, Green, Yellow, Orange, Red and Other Oral Bacteria). Cells’ filling denote Evalue (>=7 is considered as true positive, purple to red filling). Only hits with more than 10 reads are displayed.

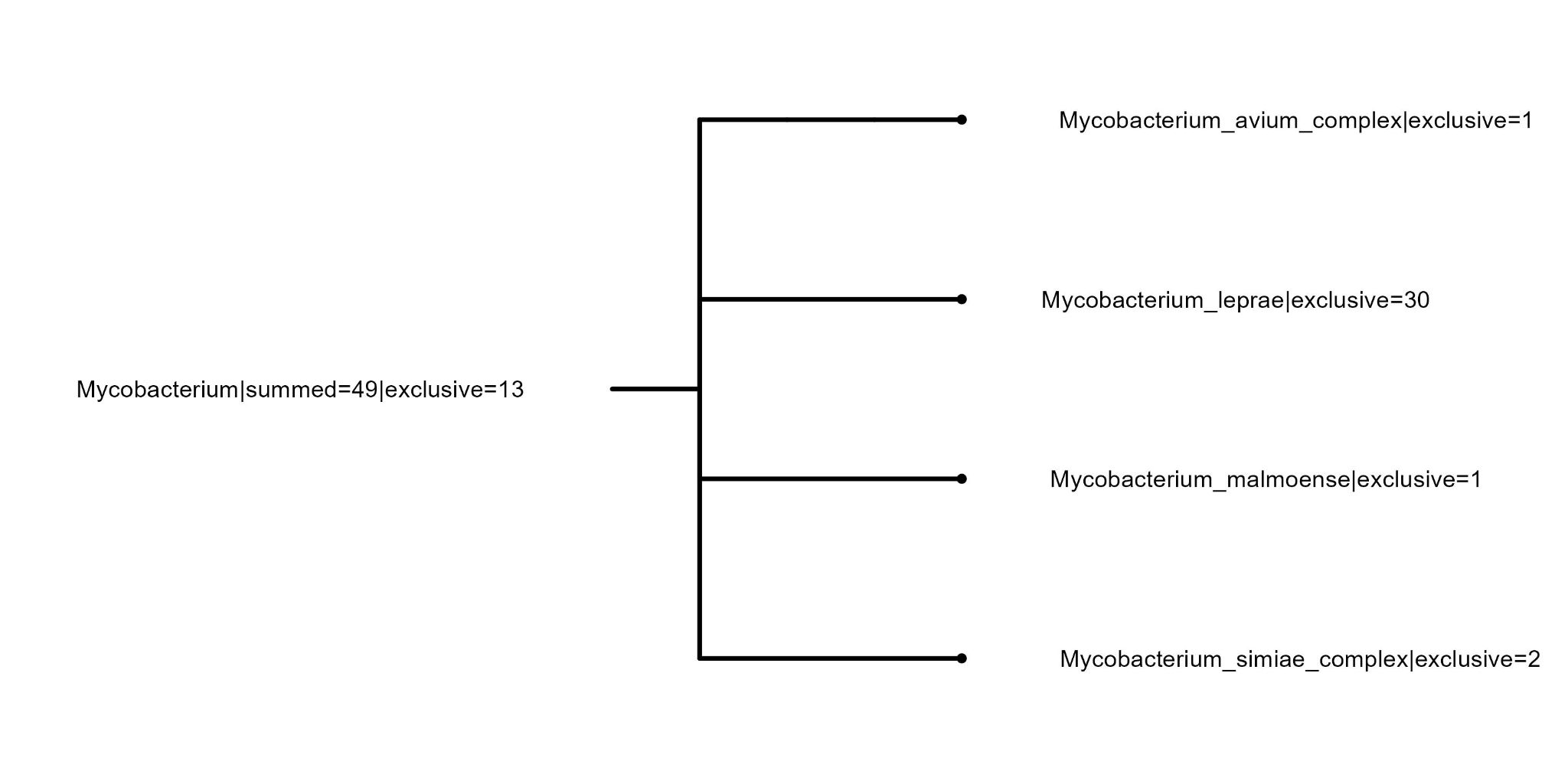

**Figure S6.** Blastn results of reads assigned by KrakenUniq (51) to *Mycobacterium leprae* in sample CHRY051B. From the initial number of sequences, 30 are exclusive of *M. leprae,* while 13 are common to the *Mycobacterium genus.*

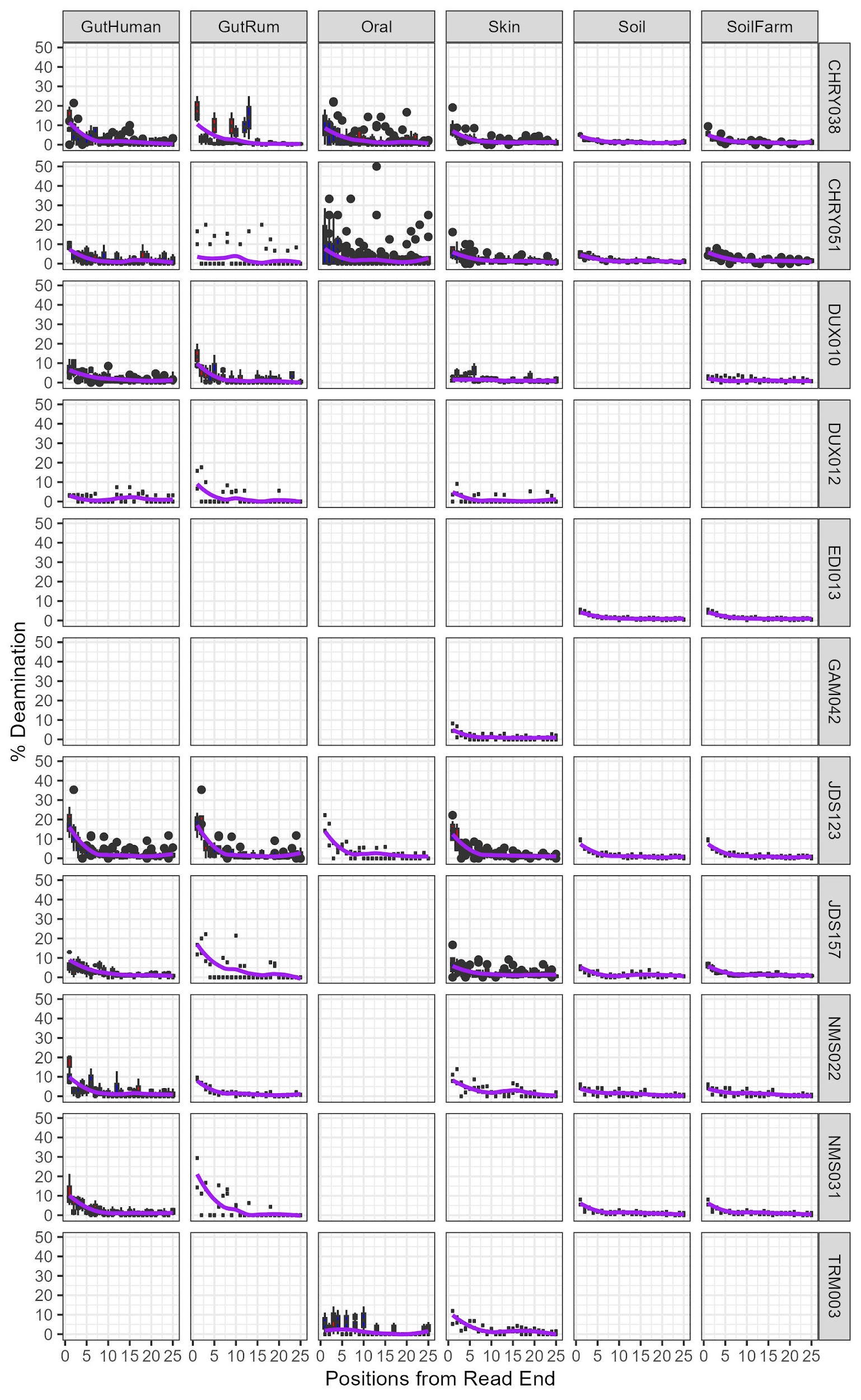

### Figure S7. Combined deamination rates at the forward (red boxes) and reverse (blue boxes) DNA strands of the microbial species assigned to different metagenomic sources. Only species with more than 100 mapped sequences with strict settings (quality 37 and ED of 0.1) are represented. Average deamination rates between 5’ and 3’ ends are displayed as a purple line.

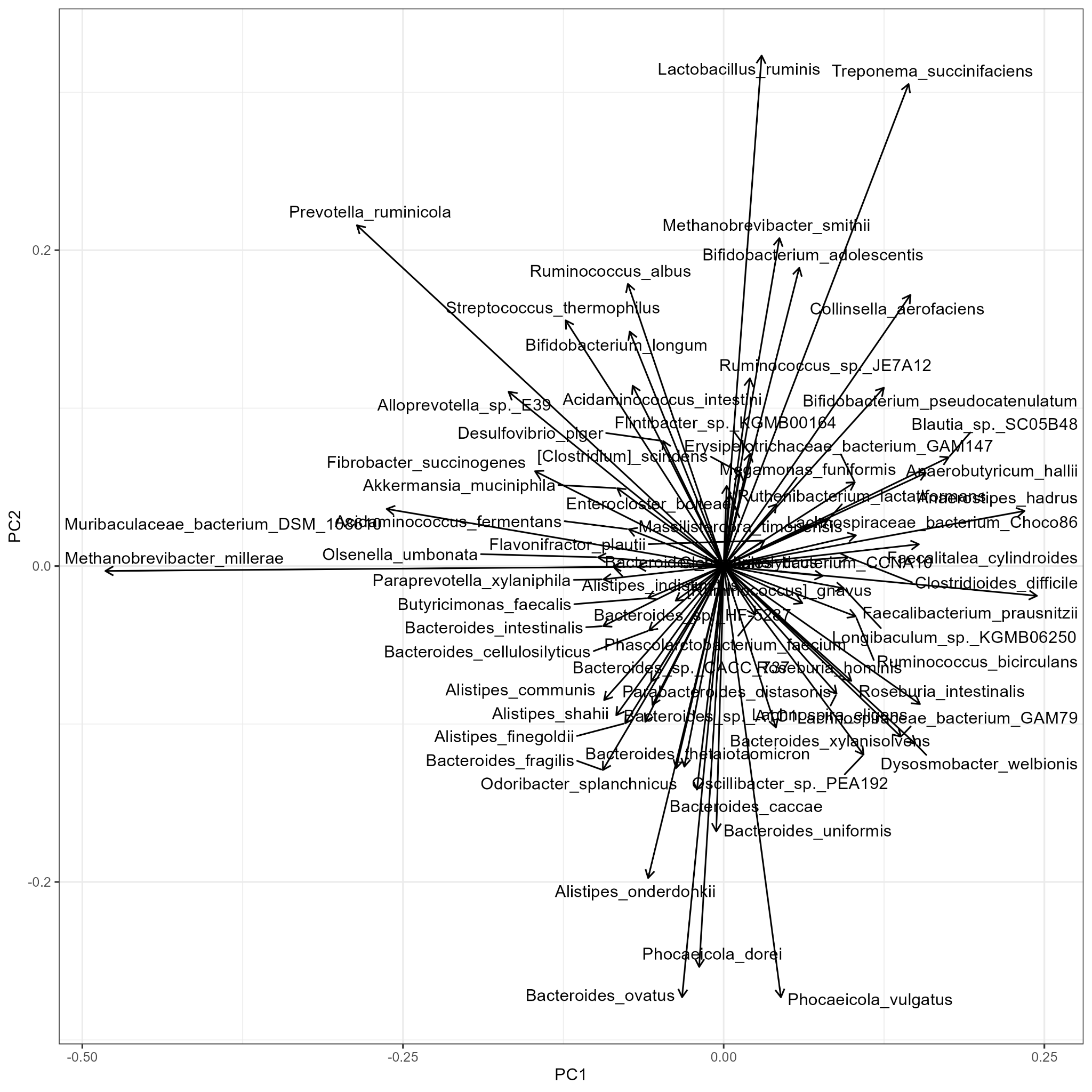

### Figure S8. PCA loadings for the gut microbiome PCA with the Human and Ruminant Gut microbial species assigned by sourctracker2.

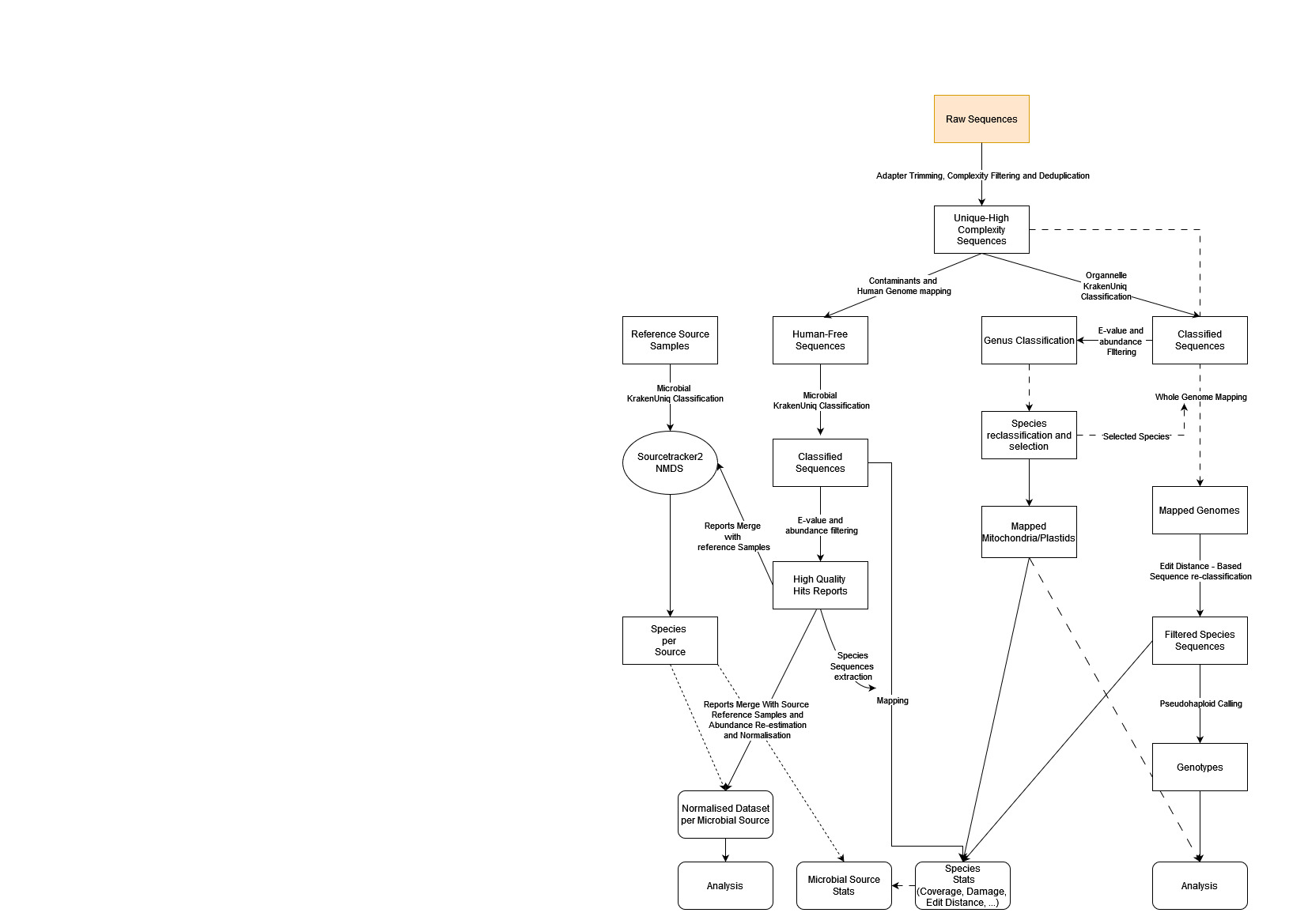

**Figure S9.** Diagram of custom pipeline developed for this study.

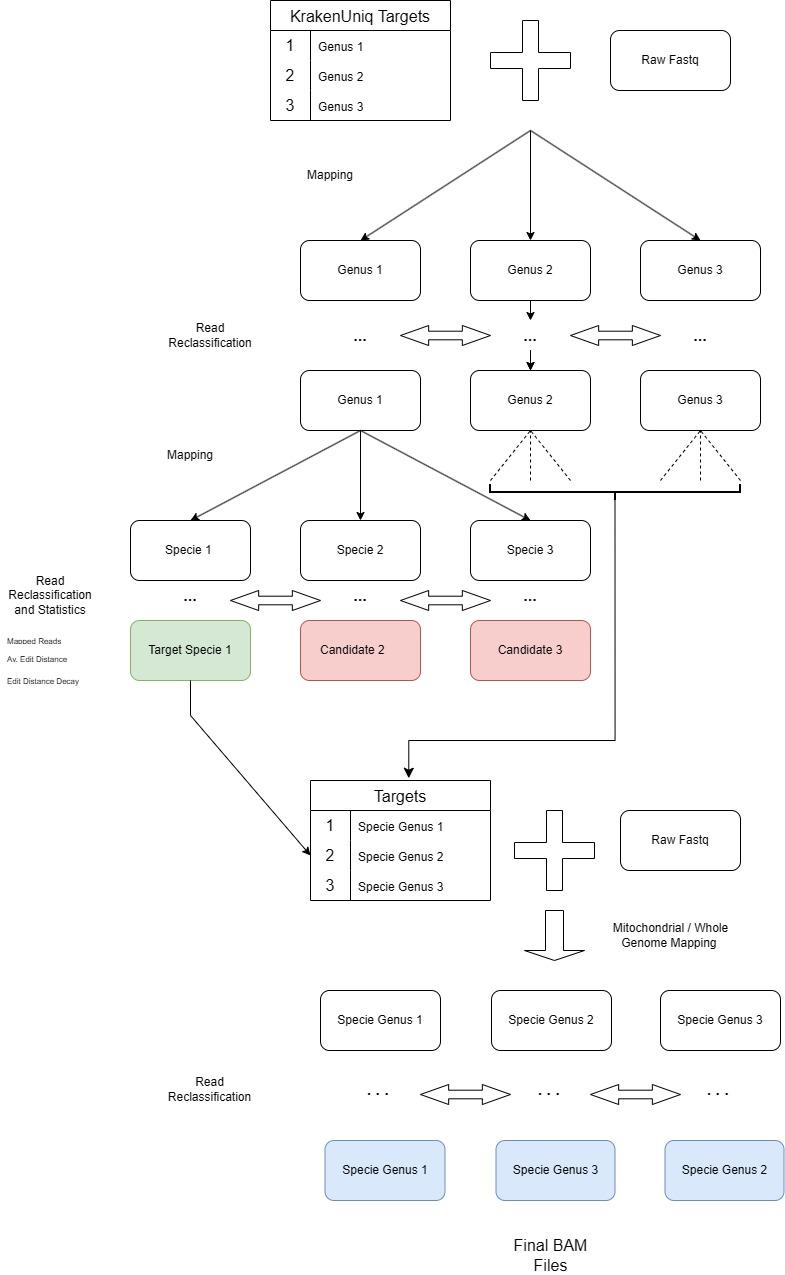

**Figure S10. Figure S9.** Diagram of the approximation used to identify Eukaryotic species.

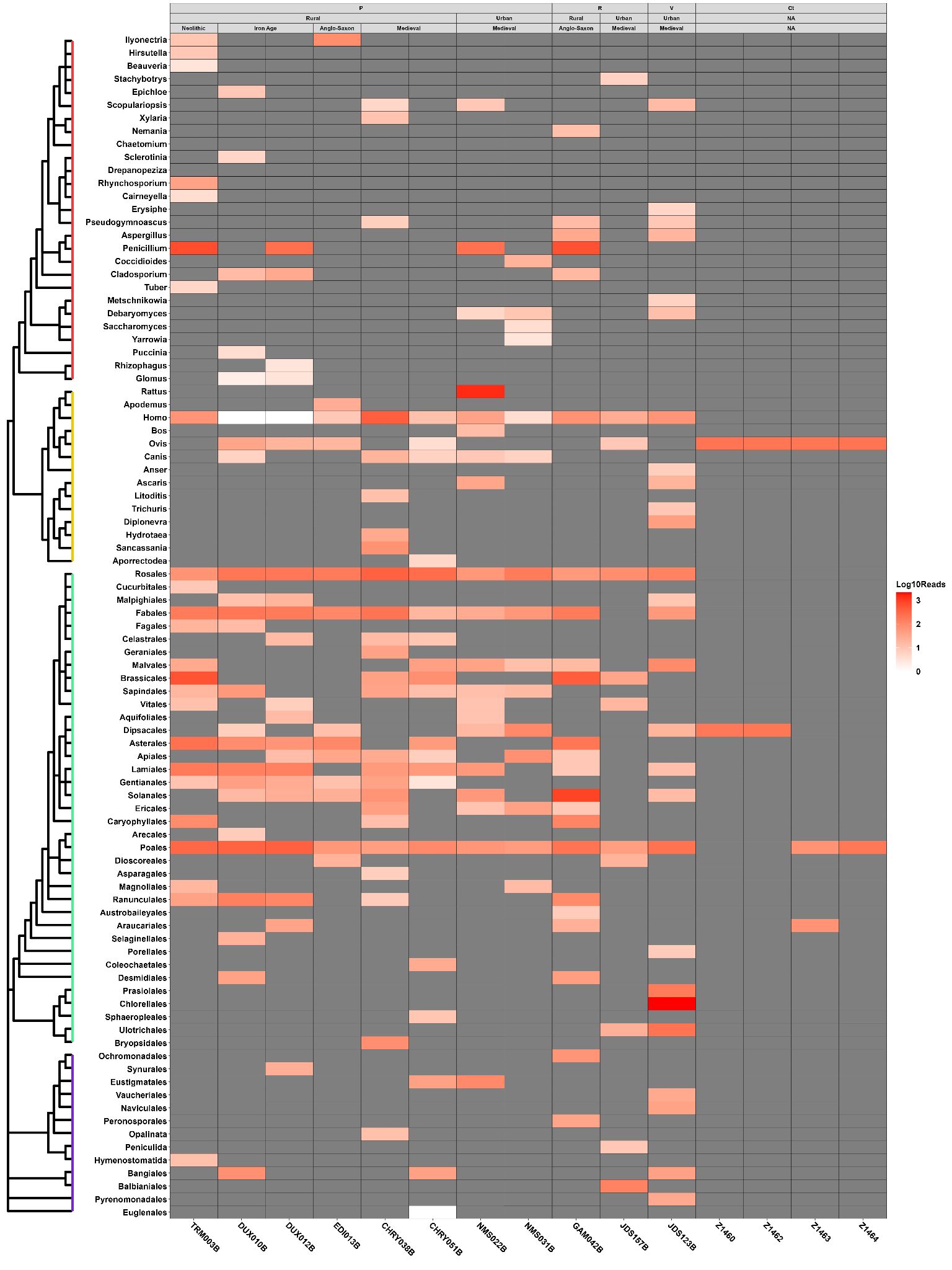

**Figure S11.** Heatmap of Eukaryotic Organelle DNA found in the samples (including controls). Vertical axis is ordered by the taxonomic location of detected species, with colours denoting the clades of interest (Red=Fungi, Yellow=Animals, Green=Viridiplantae, Purple=Protist). Fungi and Animals are classified down to genus level, while Plants and Protist to order level. Samples are grouped by their sampling origin ( P = Petrous Bone Soil, R = Rib Soil, V = Vertebrae Soil, Ct = Control), context of site (Rural vs Urban), and site datation (Neolithic, Iron Age, Anglo-Saxon and Medieval). Cells are filled with read numbers normalised to a log10 scale, with higher numbers displaying a darker red.

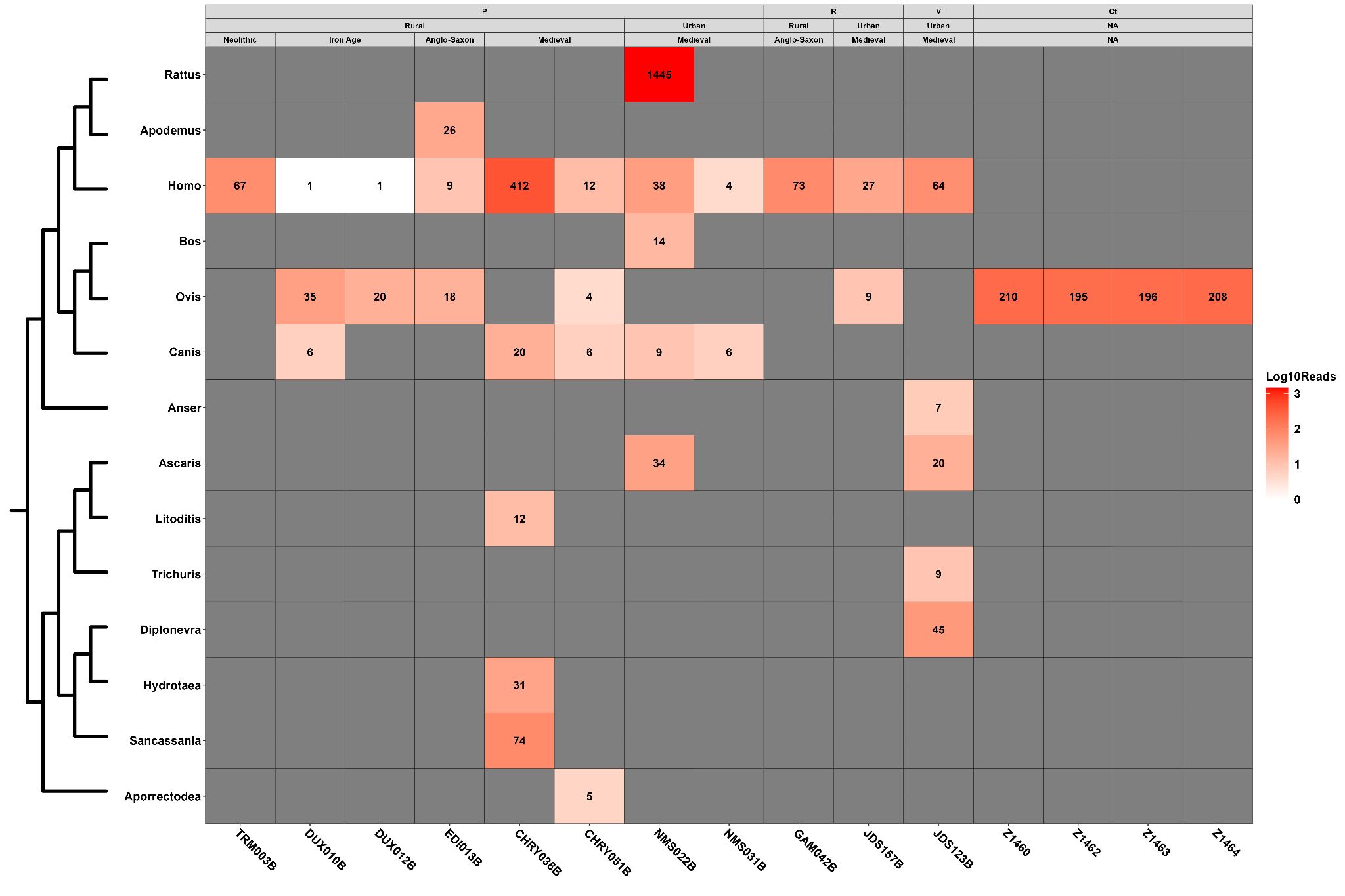

### Figure S12. Zoom in of Animal mitochondrial sequences found in the samples (including controls). The number of mapped reads is displayed inside each cell. The vertical axis is ordered by the taxonomic location of detected species. Samples are grouped by their sampling origin ( P = Petrous Bone Soil, R = Rib Soil, V = Vertebrae Soil, Ct = Control), context of site (Rural vs Urban), and site datation (Neolithic, Iron Age, Anglo-Saxon and Medieval). Read numbers are in a log10 scale with higher numbers displayed in a darker red.

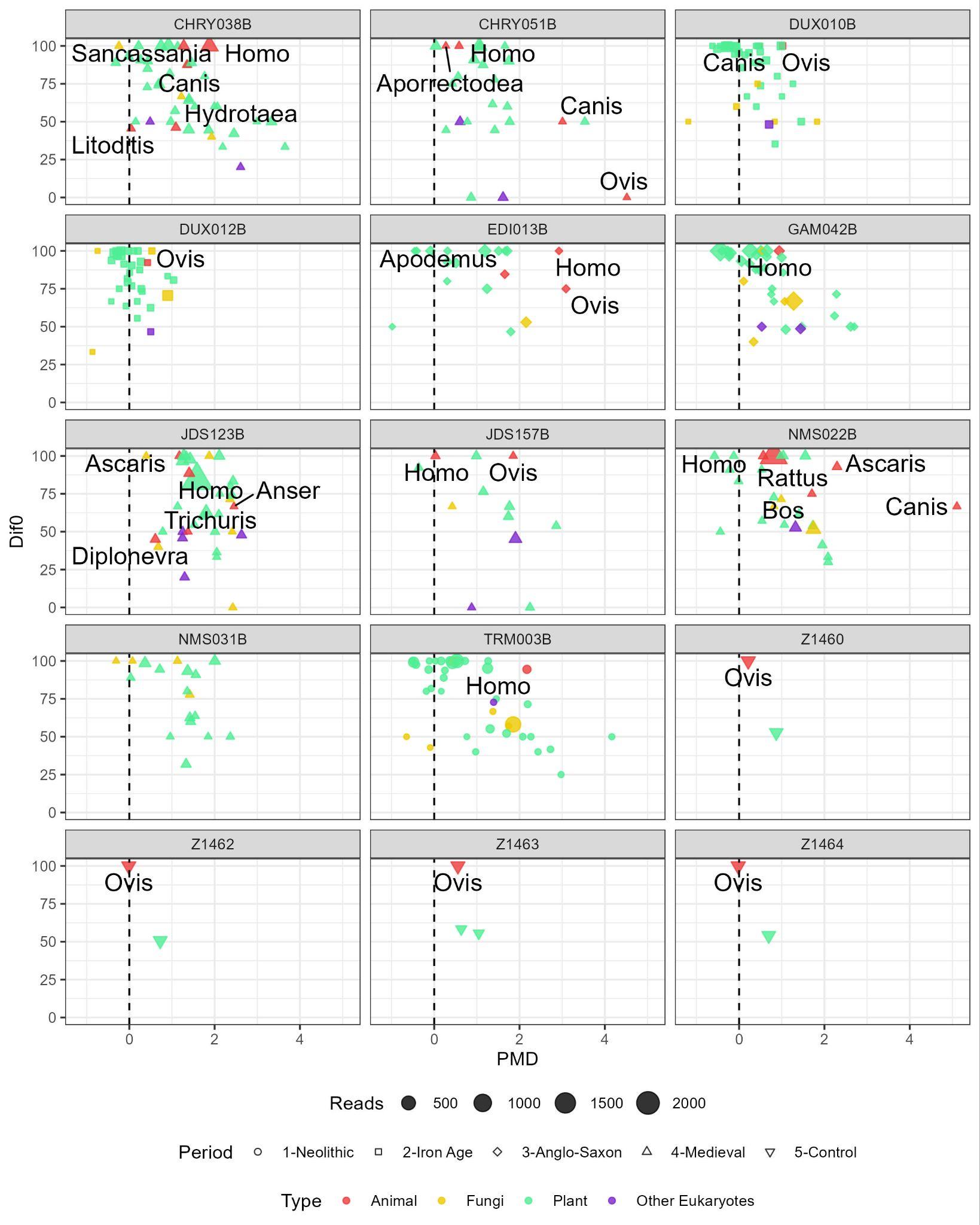

### Figure S13. Edit Distance differential against PMD score in retrieved organelle sequences per genus. Size depicts number of sequences and colour taxonomic classification. Animal species are annotated in the plot. Higher PMD value denotes higher damage in the sequences, while higher differential indicates better affinity to the reference genome.

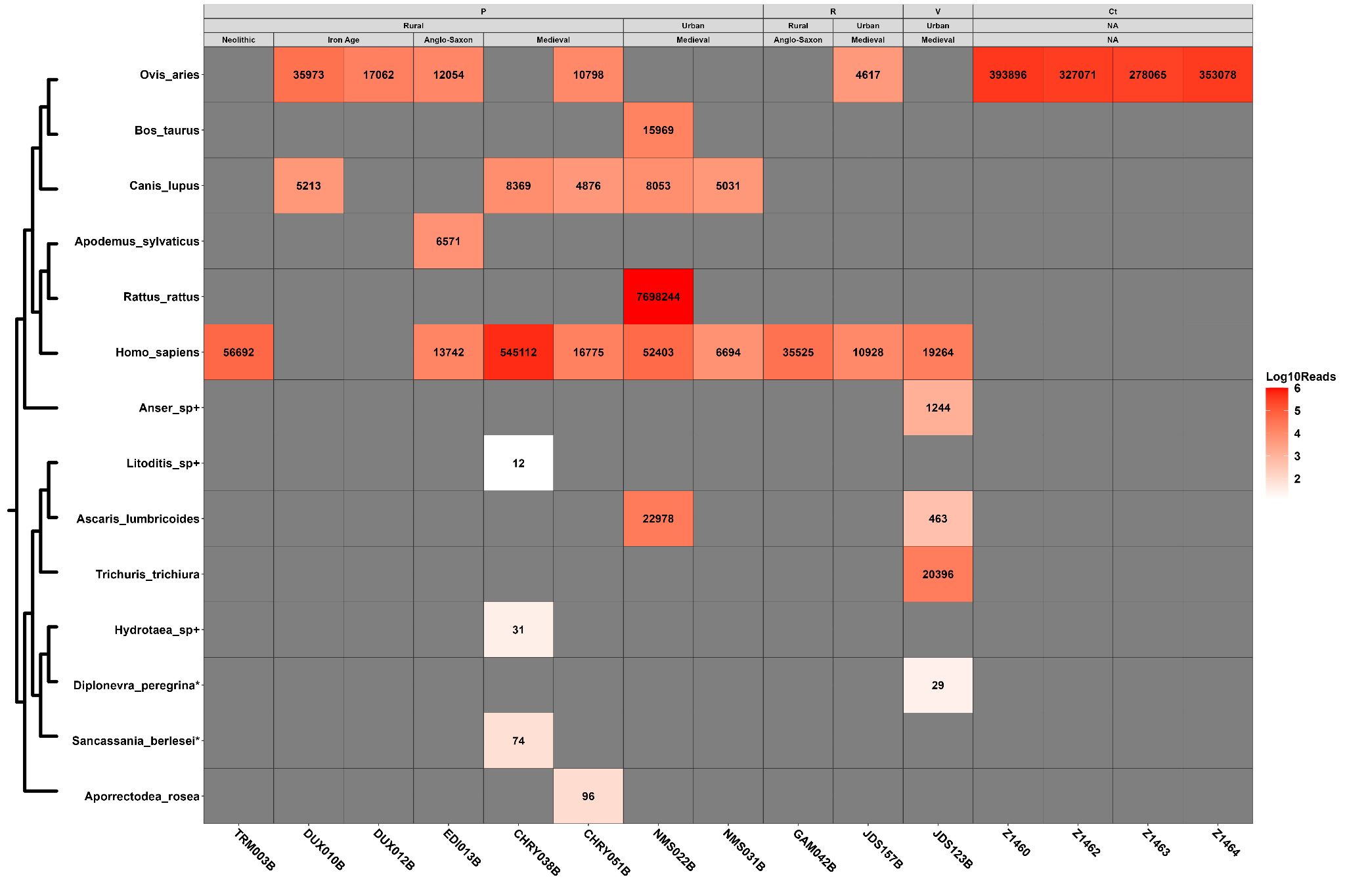
**Figure S14.** Number of reads from a select panel of animal species present in the samples and controls. The vertical axis is ordered by the taxonomic location of detected species, with some species marked by * (Only mitochondrial reference available) and + (Taxonomic level could not be determined below genus level). Horizontal axis is the sample. Samples are grouped by their sampling origin ( P = Petrous Bone Soil, R = Rib Soil, V = Vertebrae Soil, Ct = Control), context of site (Rural vs Urban), and site datation (Neolithic, Iron Age, Anglo-Saxon and Medieval). Read numbers are in a log10 scale with higher numbers displayed in a darker red.

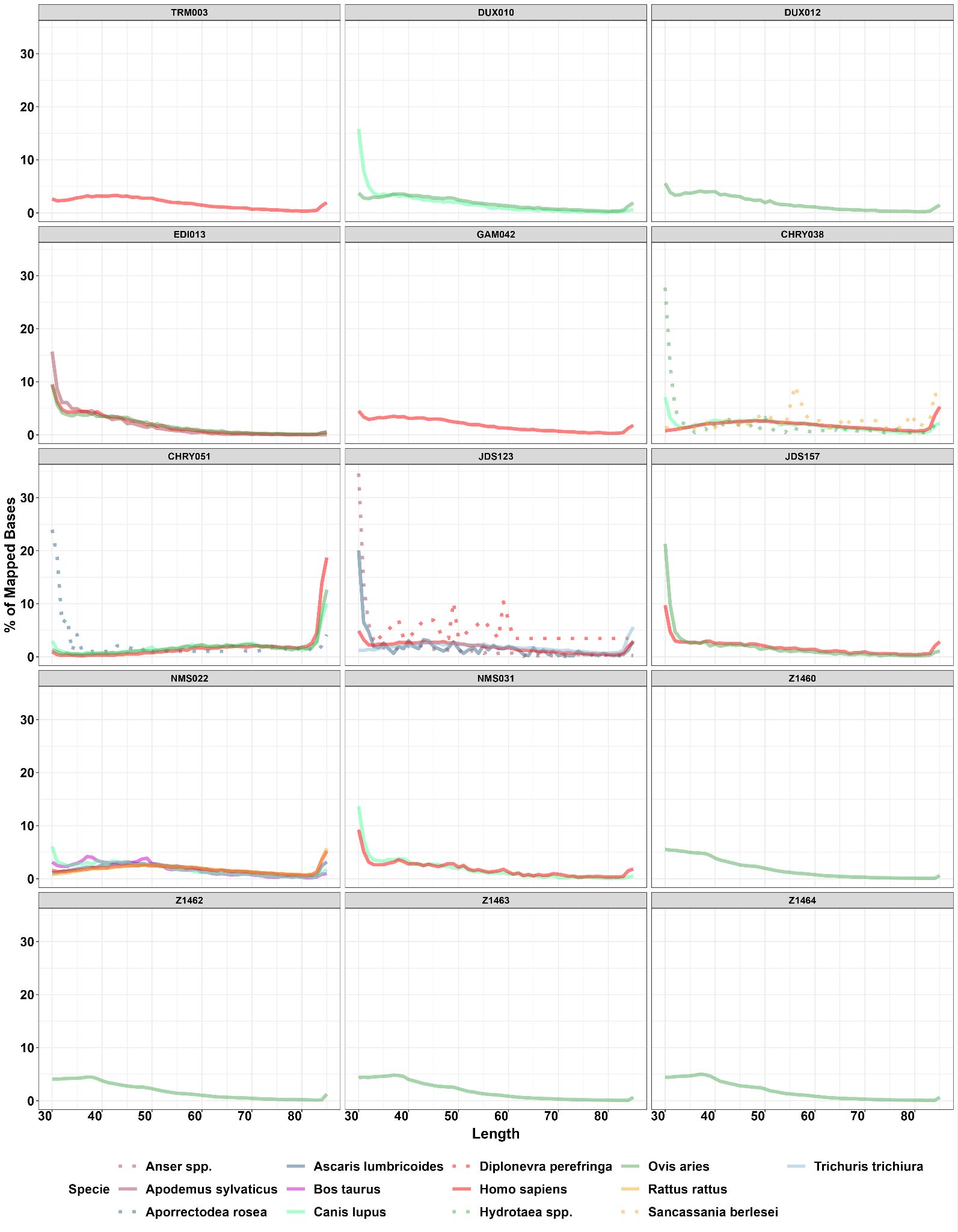

### Figure S15. Read length distributions for each species per sample.

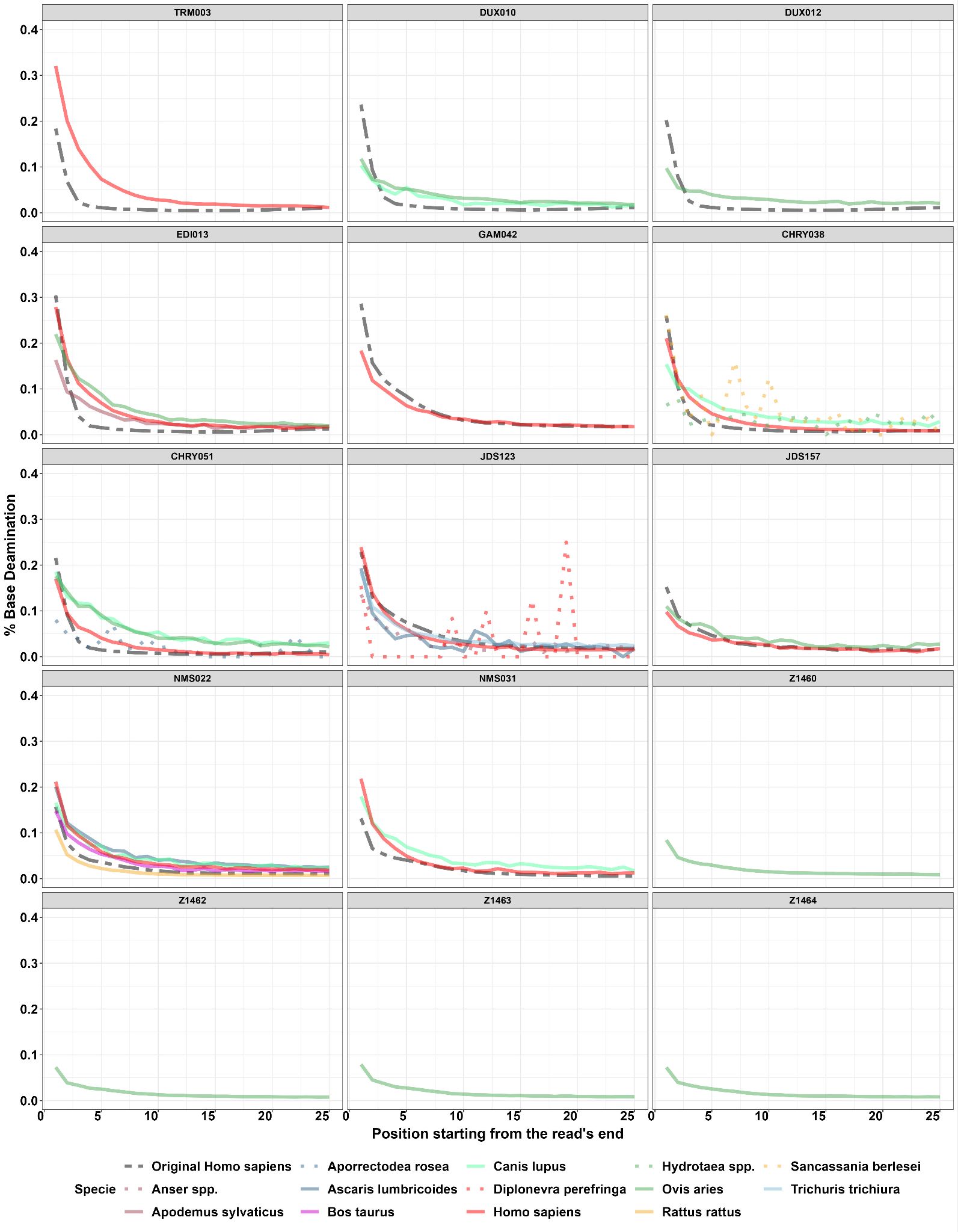

### Figure S16. Damage patterns from each species in each sample.

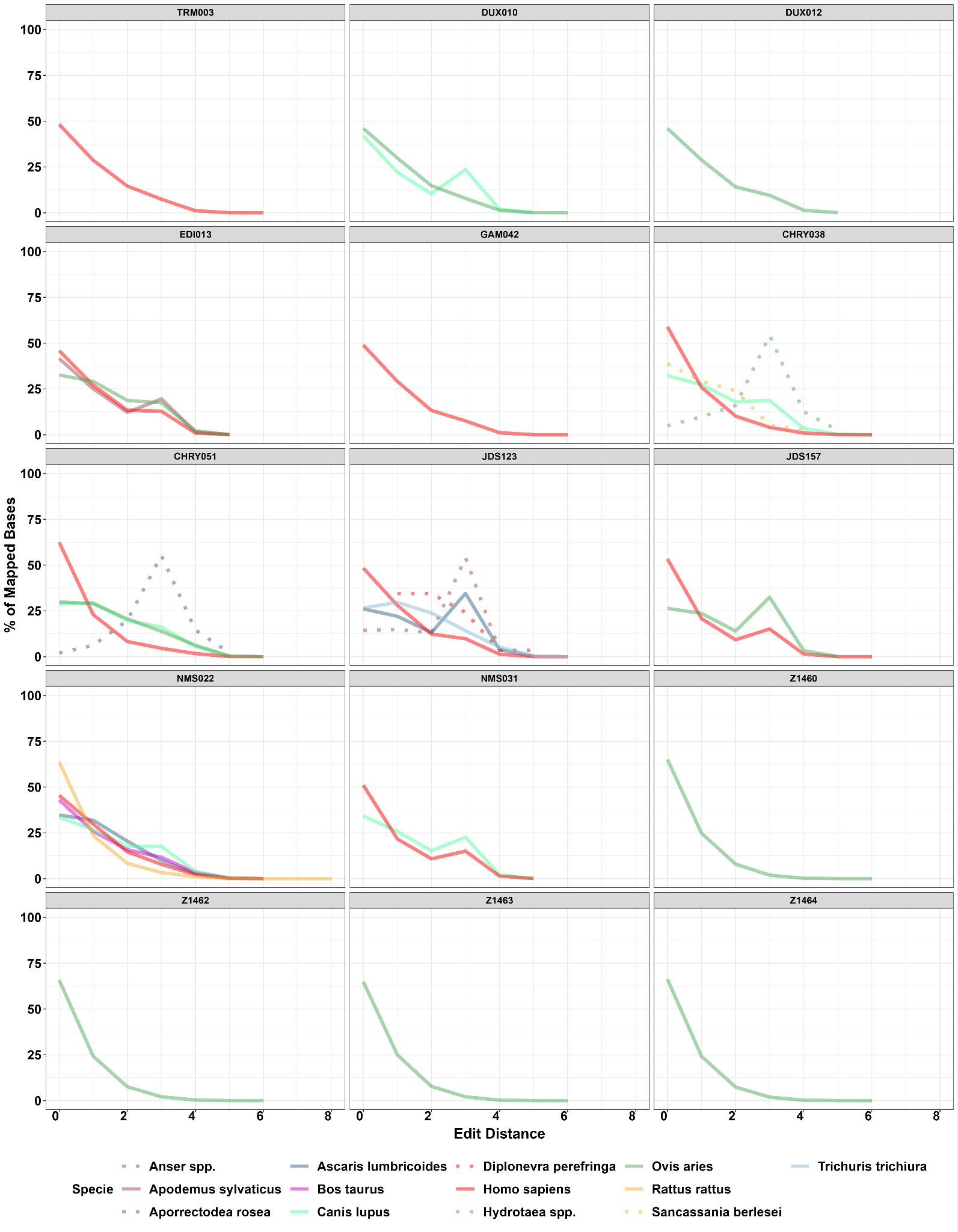

### Figure S17. Edit distances from each species in each sample.

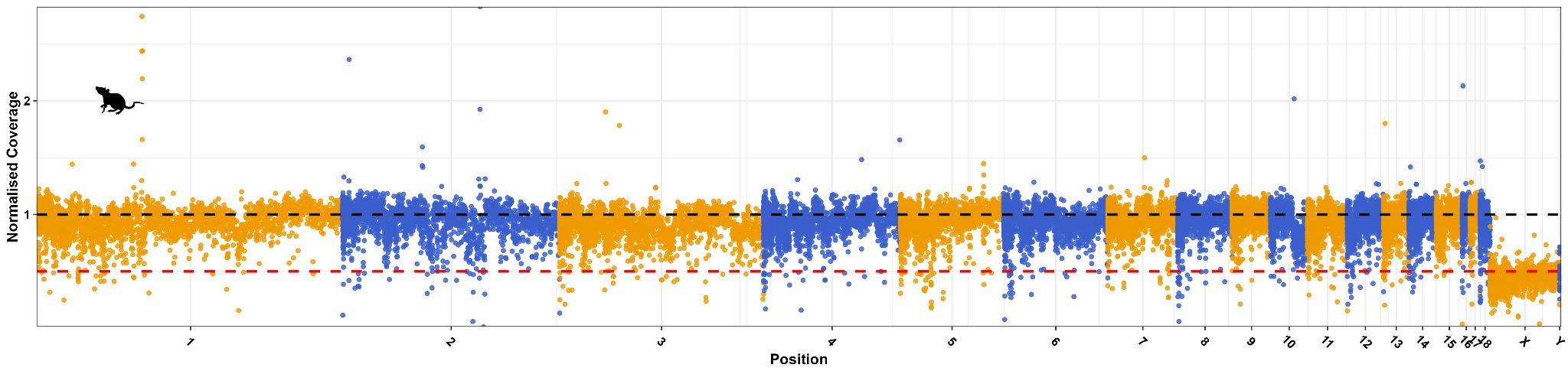

### Figure S18. Normalised coverage distribution for *Rattus rattus* genome. Chromosome X and Y present a normalised coverage of approximately 0.5, indicating that the individual is most probably a male.

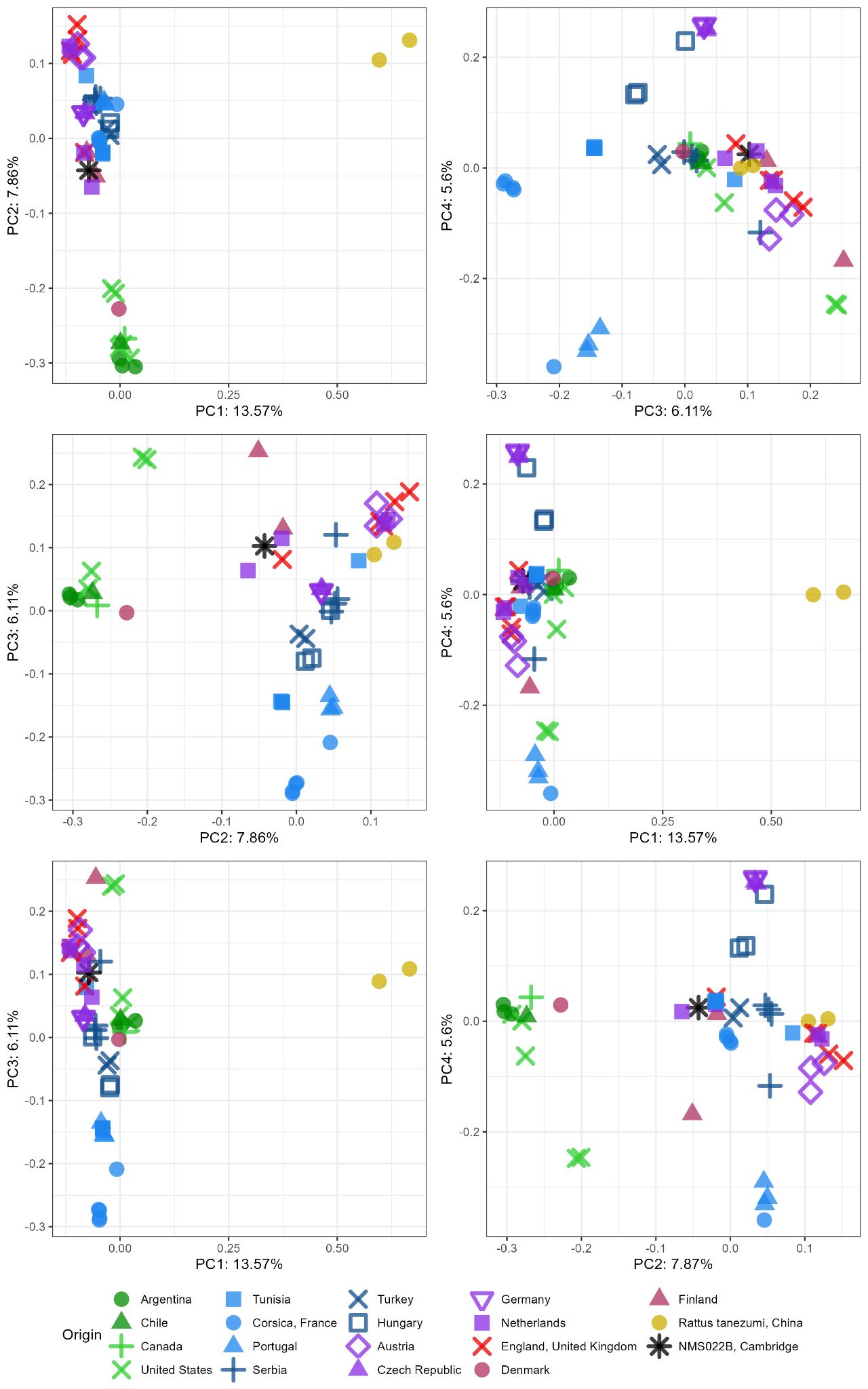

### Figure S19. Major components in the *Rattus rattus* PCA, including PC1, PC2, PC3 and PC4.

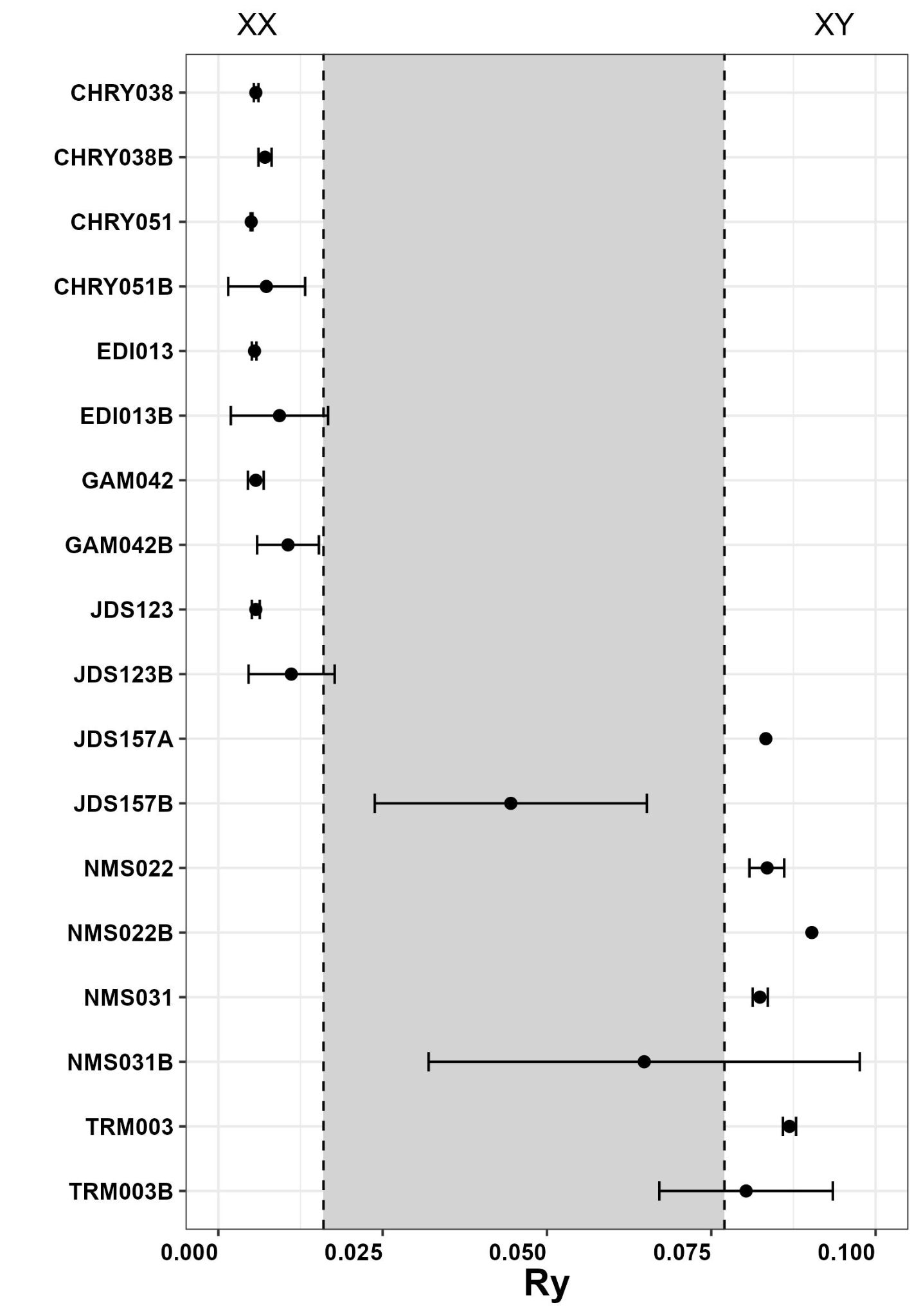

### Figure S20. Sex determination for human sequences in each sample. Sample ID without letter B is the original skeletal sample as published in (Hui et al. 2024, Keller et al. 2019, Scheib et al. 2019, or Scheib et al. *in preparation*).

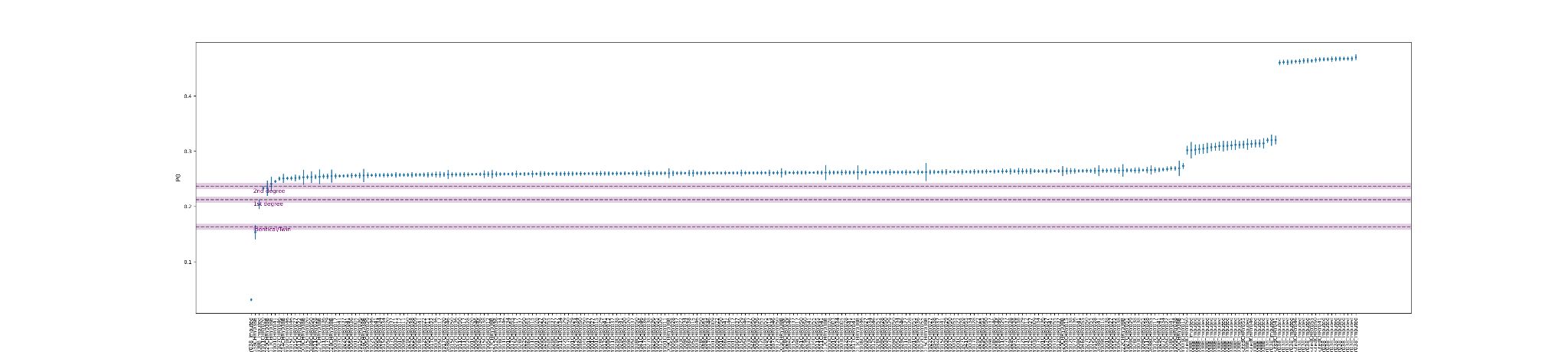

### **Figure S**21**.** Degree of relatedness determined by READv2 between CHRY038, CHRY038B, and other published contemporary individuals from the Cherry Hinton site. CHRY038 and CHRY038B are designed as the same individual/identical twins before (blue) and after (red) imputation.

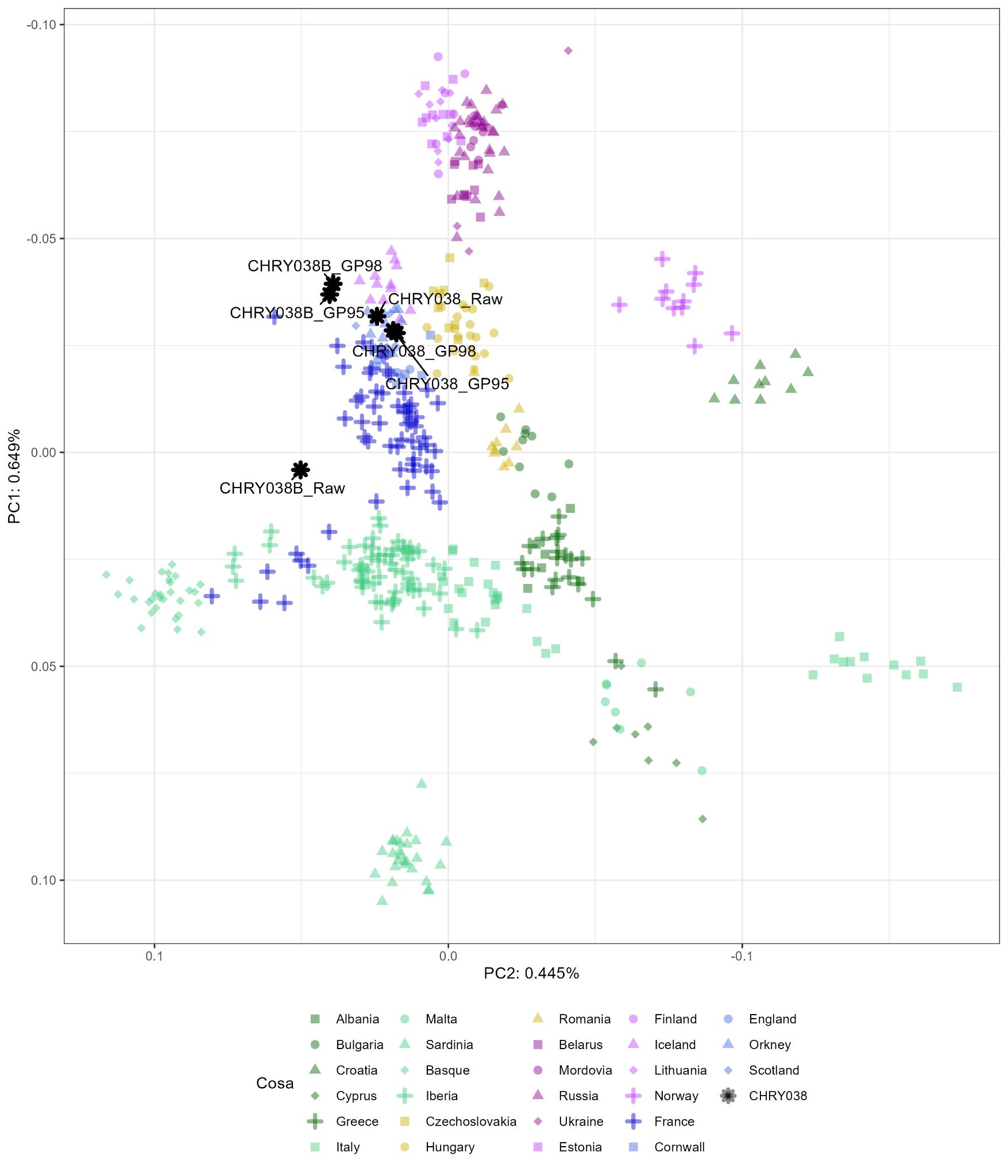

### Figure S22. CHRY038 human sequences PCA, including both sequences retrieved from the original skeletal element (CHRY038) and soil (CHRY038B), with different levels of imputation; No Imputation (Raw), Genotype Probability of 95 (GP95), and Genotype Probability of 98 (GP98).

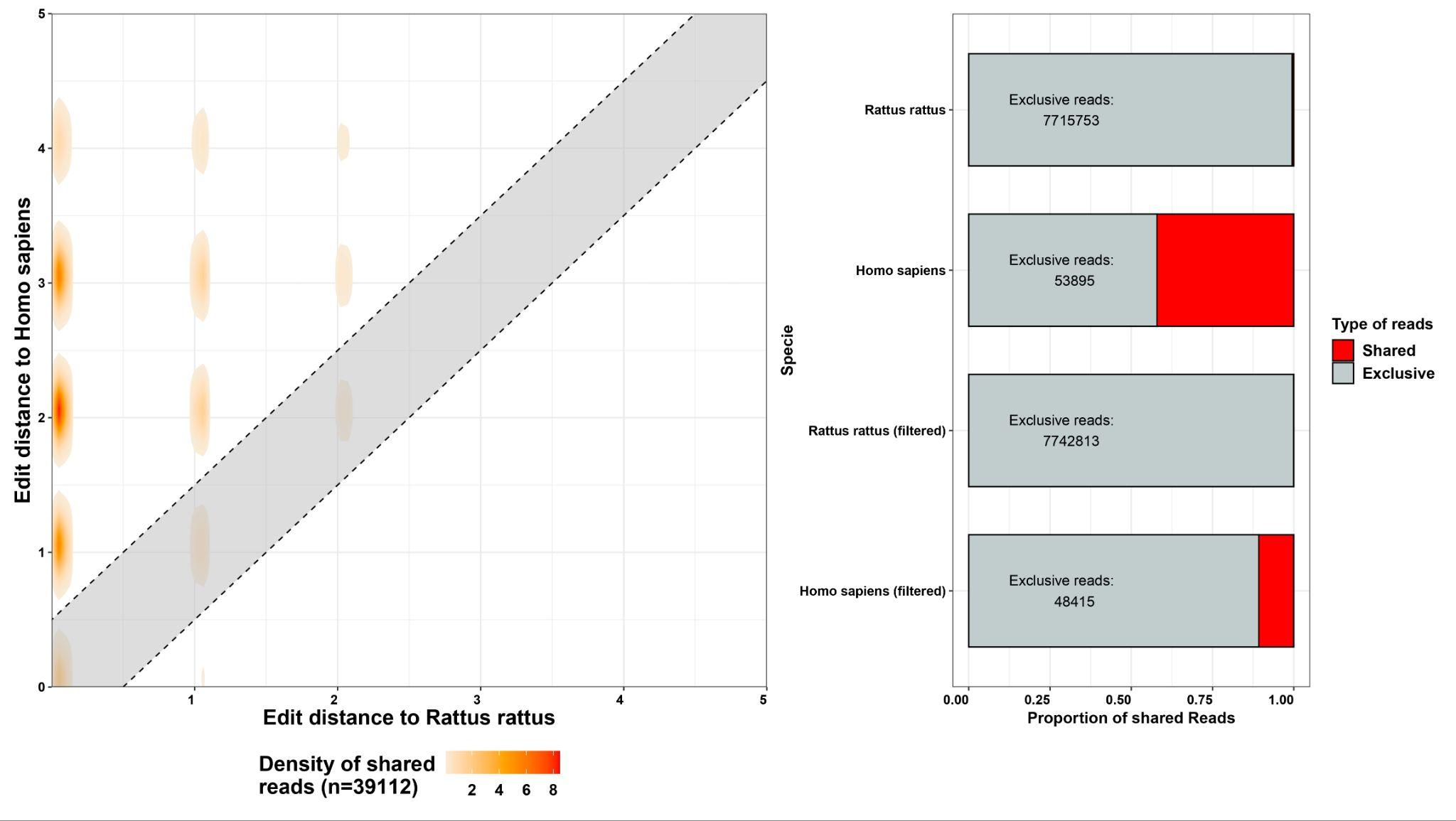

**Figure S23.** Example of cross mapping between 2 evolutionary close species, *Homo sapiens* and *Rattus rattus.* Approximately 40% of the sequences mapping to *H. sapiens* also map against *R. rattus.* Applying a filter based on edit distance to the reference genome, we can reduce this value to 10%.

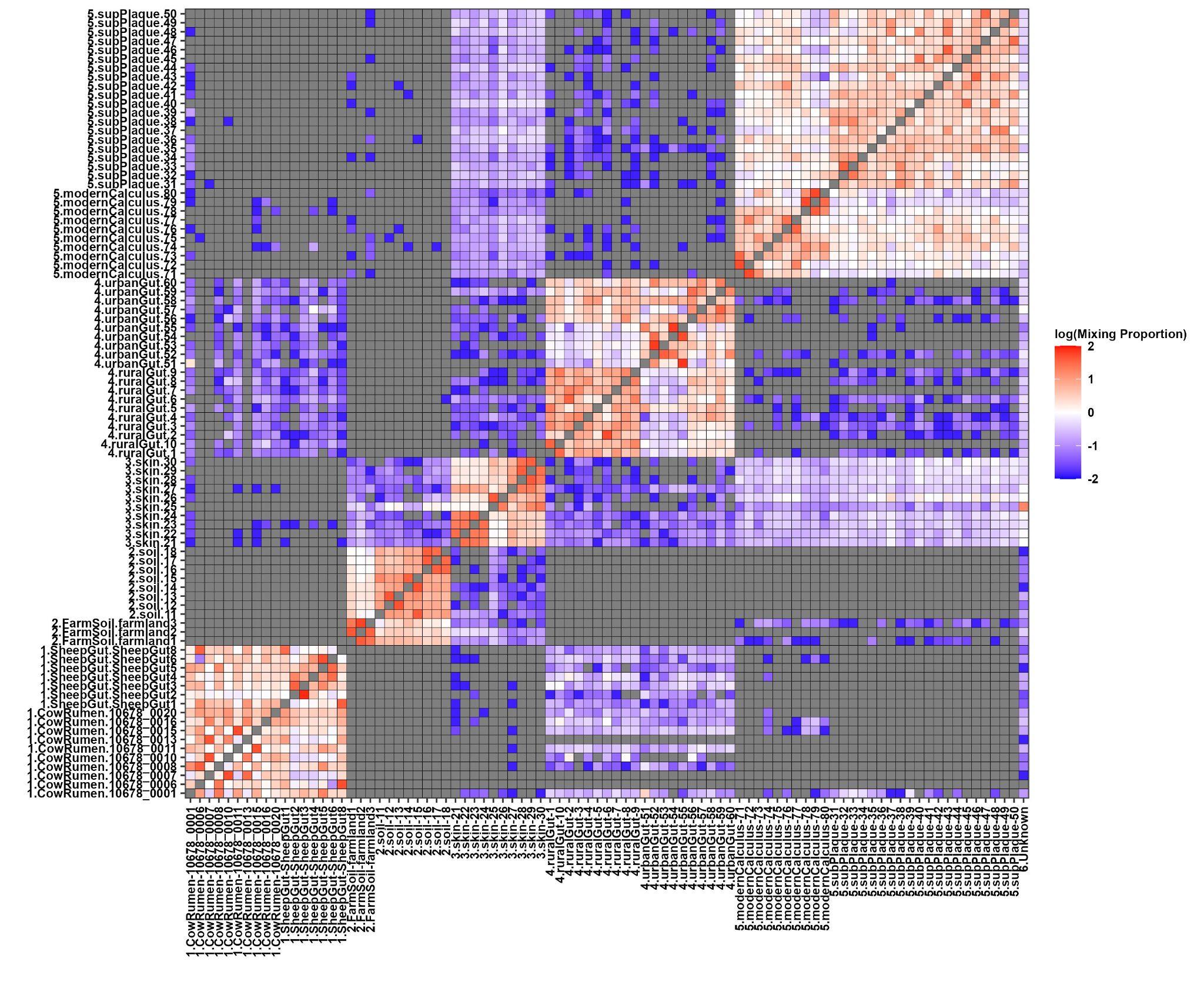

**Figure S24.** Microbial abundances affinity as estimated by sourcetracker2 “*leave one out”* function in the different reference microbiome sources.

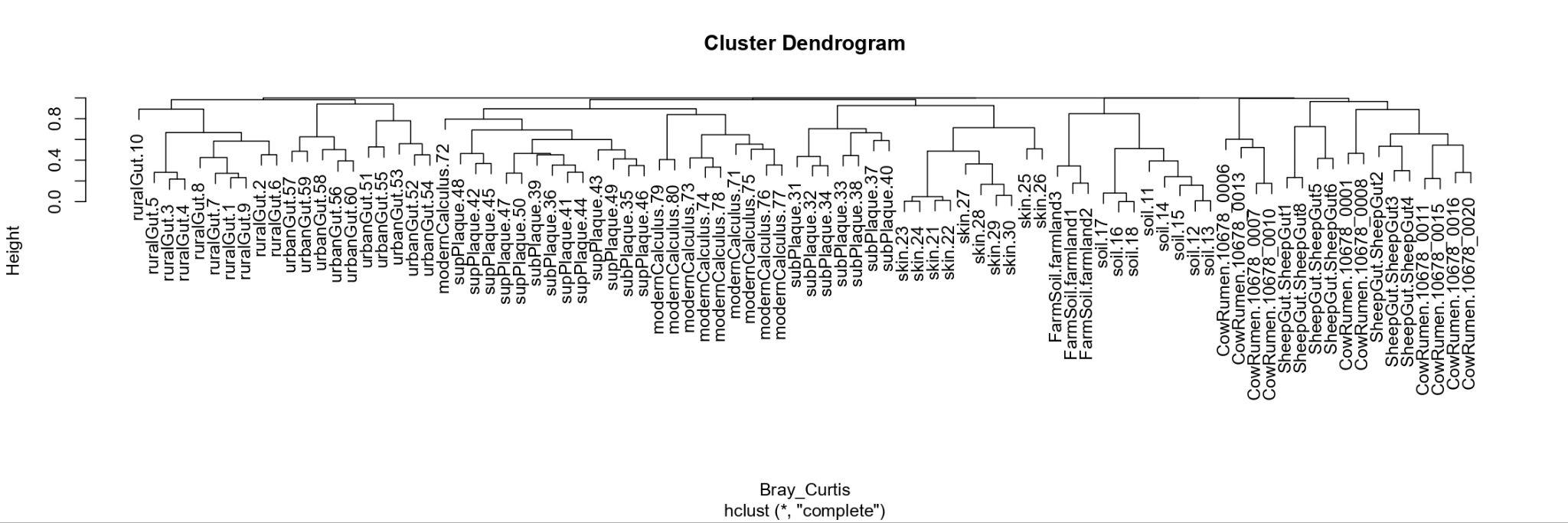

**Figure S25.** Hierarchical clustering analysis of the Bray-Curtis distances from the different microbial abundances in the Sourcetracker2 reference sources.

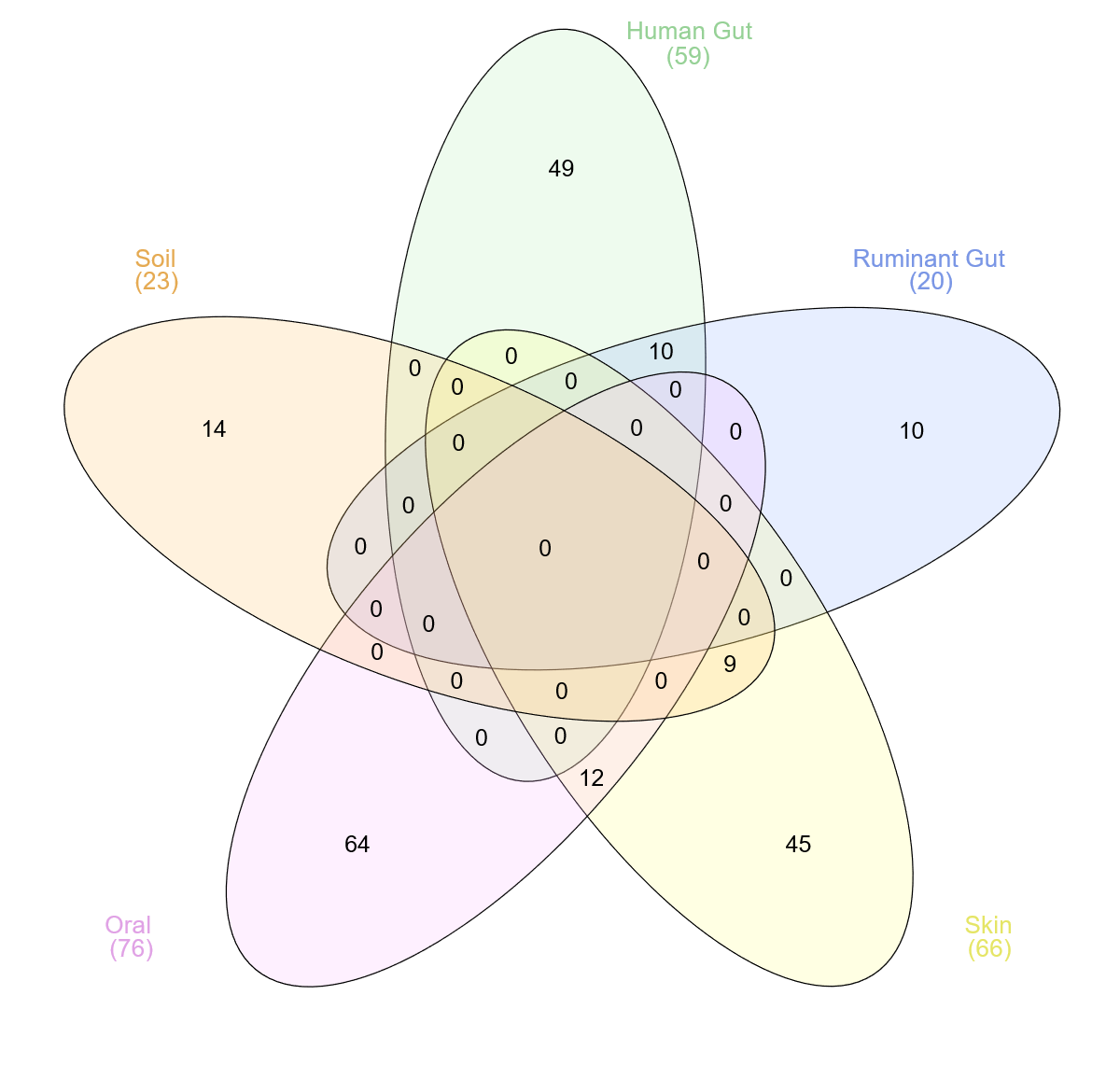

**Figure S26.** Number of characteristic microbial species determined by sourcetracker2. Species overlap is found between human gut and rumen (HuRm=10), soil and skin (SlSk=9), and oral and skin (OrSk=12).

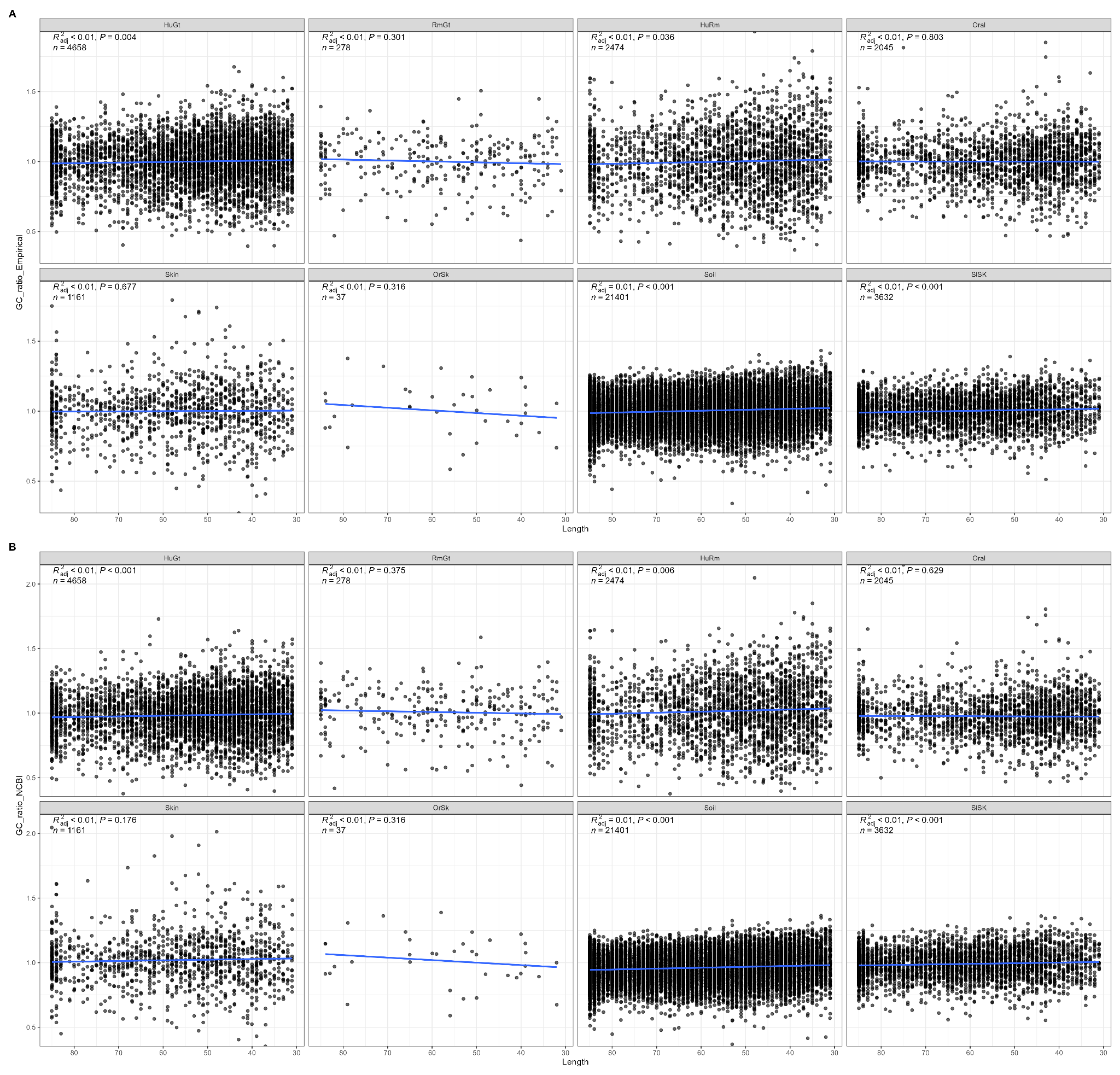

**Figure S27.** GC content distribution across read length of source-specific microbial species with more than 10 mapped reads after KrakenUniq E-value filtering above 7. GC content bias associated was tested using a linear regression in R. Panel A) shows GC content normalised to its empirical observed value while B) shows GC content normalised to the assembly GC ratio stated in NCBI.

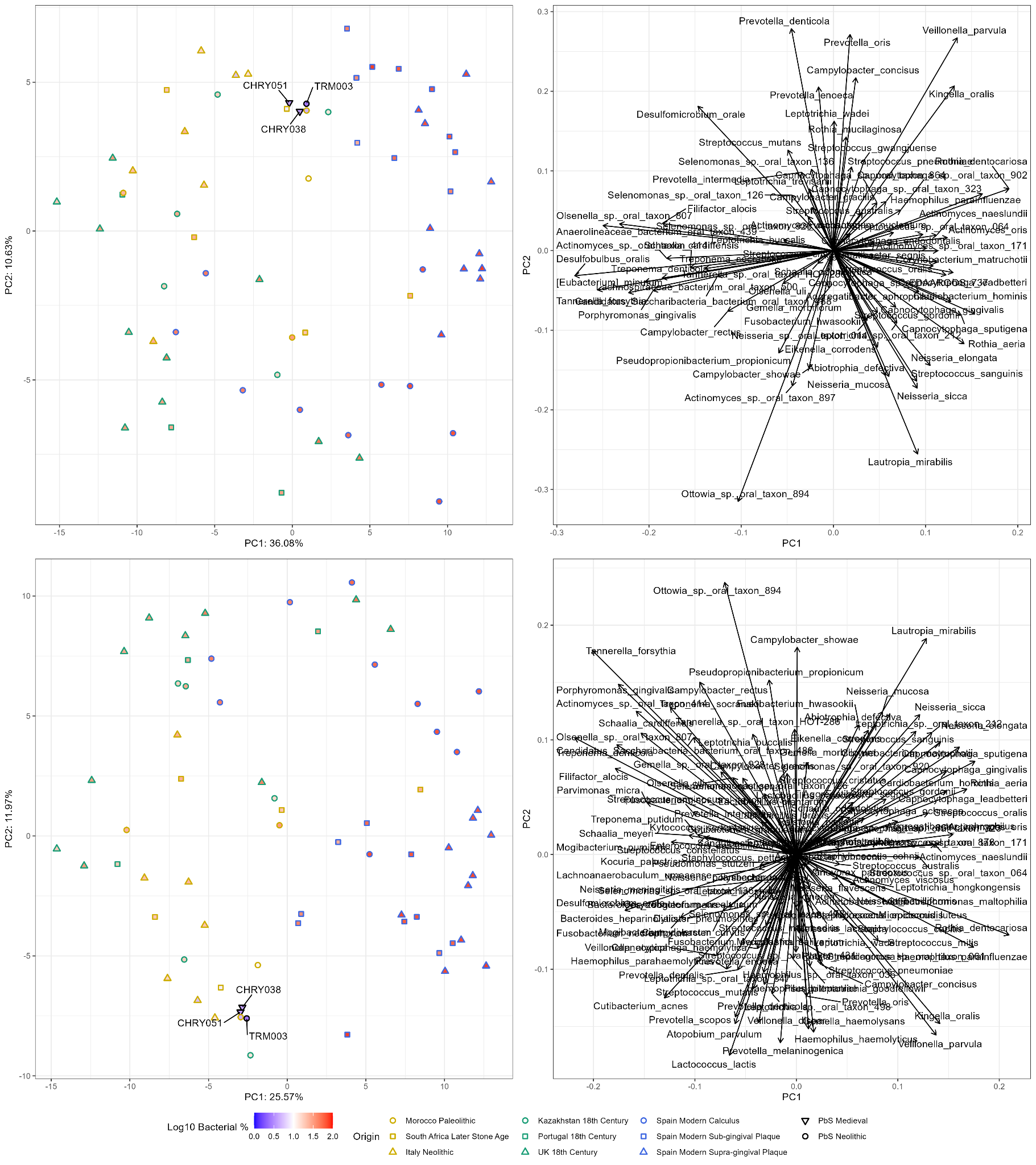

**Figure S28.** PCA and PCA loadings for species characterised as oral by sourcetracker2 (upper panels) and by the Human Oral Microbiome Database (HOMD) (lower panels). We used samples with higher numbers of Oral hits (CHRY038B, CHRY051B, TRM003B), and published oral microbiomes spanning from the Paleolithic to the Present. Dots are filled with the log10 normalised value of the percentages of microbial sequences in each sample.

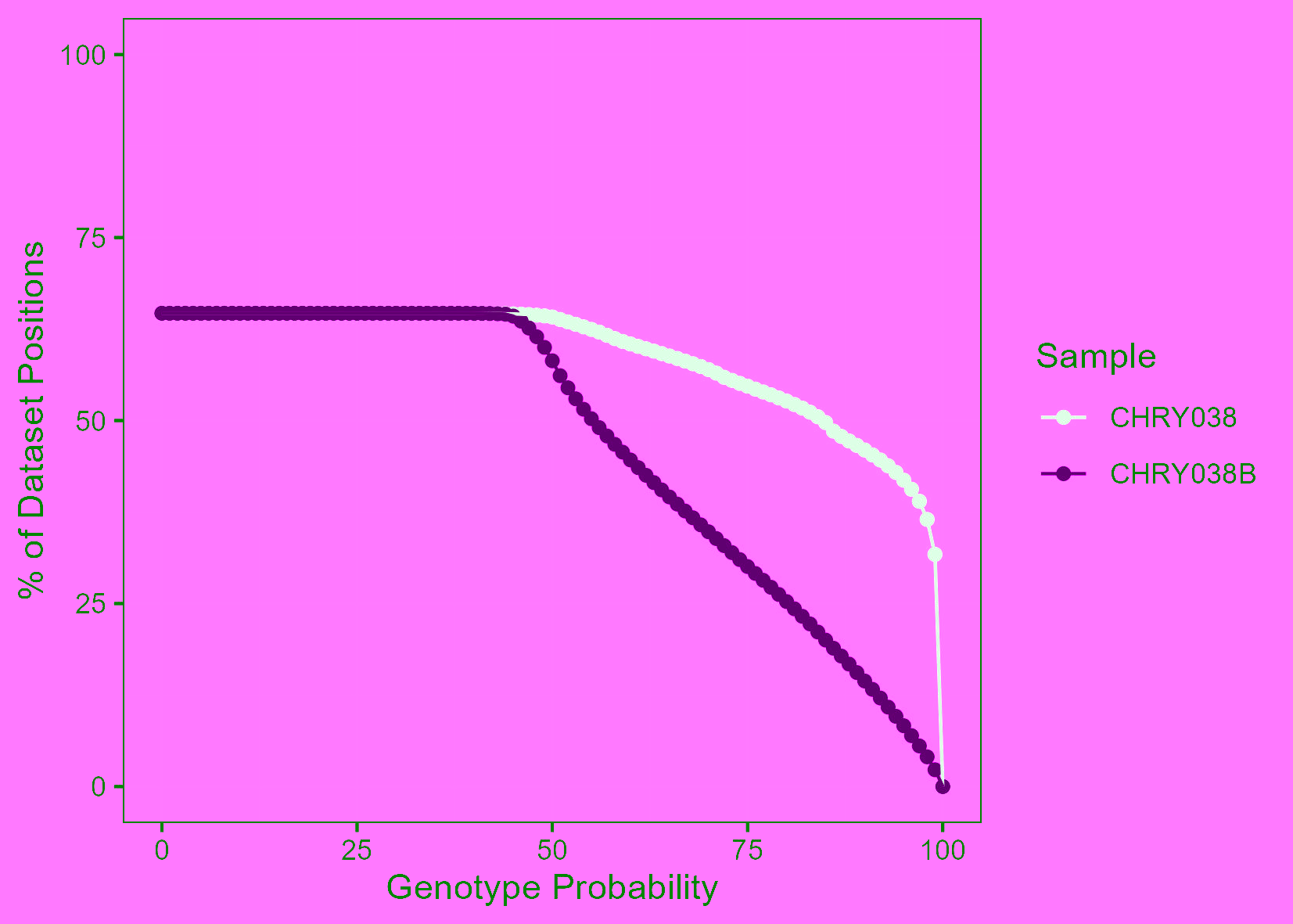

**Figure S29.** Percentage of the HO dataset SNPs covered after imputation in the original petrous bone sequenced (CHRY038) and in the adhered soil (CHRY038B) in relation to the genotype probability. The number of covered SNPs in CHRY038B rapidly decay with more stringent filtering values, most probably caused by the low number of initial SNPs (8,580).

### Supplementary Tables

**Table S1.** Raw sequences generated for each sample before and after prefiltering. All pre-filtered reads are at least 30bp and quality 30 as stated in the main methods section

| **Sample** | **Raw Reads** | **Unique High Complexity Reads** |
| --- | --- | --- |
| CHRY038B | 19,964,480 | 17,88,2745 |
| CHRY051B | 21,743,979 | 20,161,737 |
| DUX010B | 17,455,2580 | 14,704,196 |
| DUX012B | 15,229,7970 | 13,046,691 |
| EDI013B | 19,470,0280 | 14,815,381 |
| GAM042B | 19,753,5280 | 15,138,649 |
| JDS123B | 18,537,8100 | 15,643,459 |
| JDS157B | 17,856,6780 | 13,749,513 |
| NMS022B | 20,149,396 | 17,253,437 |
| NMS031B | 16,004,4520 | 10,258,581 |
| TRM003B | 17,596,592 | 14,956,668 |
| GAM042 | 32,248,147 | 18,431,240 |
| JDS157A | 16,405,420 | 4,477,441 |
| Z1460 ( Control-1) | 19,823,619 | 9,997,561 |
| Z1462 ( Control-2) | 18,134,387 | 9,165,025 |
| Z1463 ( Control-3) | 18,938,113 | 9,557,912 |
| Z1464 ( Control-4) | 20,088,743 | 10,202,713 |

**Table S2.** Proteomics screening results

**Table S3.** Sourcetracker2 results; bacterial species abundances.

**Table S4.** KrakenUniq classified Genus’ sequences of microbial Pathogens. Sequences were extracted and validated by blastn.

| **Sample** | **KrUn Reads** | **Genus** | **Species** | **Blast Reads** |
| --- | --- | --- | --- | --- |
| JDS175B | 236 | *Yersinia* | *Yersinia enterocolitica* | 10 |
| NMS022B | 91 | *Acanthamoeba* | *Acanthamoeba castellanii* | 5 |
| TRMO03B | 20 | *Entamoeba* | *-* | 0 |
| JDS123B | 10 | *Giardia* | *-* | 2 |
| CHRY051B | 53 | *Mycobacterium* | *Mycobacterium leprae* | 30 |

**Table S5.** Bacterial damages raw data.

**Table S6.** Eukaryotic screening reference genomes.

**Table S7.** *Rattus* population genetics dataset.

**Table S8.** *Rattus rattus* proteins identified from the NMS0022B sample with more than 5% coverage. The identification of tissues in which the proteins are expressed in rats or mice was done using the Uniprot database.

| **Species** | **Protein** | **Coverage with *novor cloud*** | **coverage with *Pfind*** | **Expressed in** |
| --- | --- | --- | --- | --- |
| *Rattus rattus* | alpha-2-Hs-glycoprotein isoform XI | 70.2% | 73.6% | Blood |
| *Rattus rattus* | collagen alpha-2(I) chain | 74.1% | 73.1% | Tissues |
| *Rattus rattus* | collagen alpha-1(I) chain | 71.4% | 72.5% | Tissues |
| *Rattus rattus* | alpha-2-Hs-glycoprotein isoform | NA | 72% | Blood |
| *Rattus rattus* | collagen alpha-1(II) chain | 67.1% | 70.6% | Chondrocytes |
| *Rattus rattus* | biglycan | 46.1% | 50.7% | Tissues |
| *Rattus rattus* | chondroadherin | 35.9% | 33.9% | Cartilage/bone/bone marrow |
| *Rattus rattus* | secreted phosphoprotein | 28.6% | 28.6% | Tissues |
| *Rattus rattus* | collagen alpha-3(VI) chain | 24% | 27.4% | NA |
| *Rattus rattus* | matrix Gla protein | 24.3% | 27.2% | NA |
| *Rattus rattus* | collagen alpha-1(IX) chain | 22.5% | 26.3% | NA |
| *Rattus rattus* | pigment epithelium-derived factor | 16.7% | 22.5% | Tissues |
| *Rattus rattus* | hyaluroan and proteoglycan link protein 1 | 21.5% | 21.5% | Tissues |
| *Rattus rattus* | osteopotin | 5.7% | 20.8% | NA |
| *Rattus rattus* | fibromodulin | 20.5% | 20.5% | NA |
| *Rattus rattus* | SPARC | 24.3% | 19.9% | Tissues |
| *Rattus rattus* | collagen alpha-3(IX) chain | 19% | 19% | NA |
| *Rattus rattus* | cartilage oliogomeric matrix protein | 24.9% | 18.5% | NA |
| *Rattus rattus* | collagen alpha-2(XI) chain | 12.2% | 16.4% | Tissues |
| *Rattus rattus* | C-type lectin domain family 3 member A | 16.3% | 16.3% | NA |
| *Rattus rattus* | collagen alpha-2(IX) chain | 10.5% | 16% | NA |
| *Rattus rattus* | collagen alpha-1(XII) chain | 16.4% | 15.1% | NA |
| *Rattus rattus* | lactadherin | 11% | 14.7% | Tissues |
| *Rattus rattus* | phosphoethanolamine/phosphocholine phosphatase isoform X2 | 21% | 13.1% | NA |
| *Rattus rattus* | cartilage intermediate layer protein 2 | 8% | 13% | Tissues |
| *Rattus rattus* | matrilin-3 | NA | 12.9% | Tissues |
| *Rattus rattus* | serum albumin | 11.7% | 12.8% | Blood |
| *Rattus rattus* | osteomodulin | 12.5% | 12.5% | Tissues |
| *Rattus rattus* | phosphoethanolamine/phosphocholine phosphatase isoform X1 | NA | 12.1% | NA |
| *Rattus rattus* | coagulation factor VII | 9.4% | 11.9% | Blood |
| *Rattus rattus* | fibronectin isoform X10 | 8.5% | 11.9% | NA |
| *Rattus rattus* | collagen-alpha-1(III) chain | 10.7% | 10.9% | Tissues |
| *Rattus rattus* | collagen alpha-1(XI) chain | 6.9% | 10.6% | Tissues |
| *Rattus rattus* | vitronectin | 11.3% | 9.9% | Tissues |
| *Rattus rattus* | collagen alpha-2(V) chain | 4.7% | 8.9% | Tissues |
| *Rattus rattus* | annexin A1 | 8.7% | 8.7% | Blood |
| *Rattus rattus* | prolargin | 9.3% | 7.2% | Tissues |
| *Rattus rattus* | EMILIN-1 isoform X2 | 2.3% | 6.9% | NA |
| *Rattus rattus* | collagen alpha-1(X) chain | 10.7% | 6.3% | Tissues |
| *Rattus rattus* | basement membrane-specific heparan sulfate proteoglycan core protein | 5.3% | 5.7% | NA |
| *Rattus rattus* | aggrecan core protein | 5.5% | 5.7% | NA |
| *Rattus rattus* | periostin isoform X4 | 2.8% | 5.7% | NA |

**Table S9.** Human mapping statistics and sex determination.

**Table S10.** Readv2 results.

**Table S11**. Cherry Hinton samples used for kinship analysis.

| **Individual** | **Dataset Coverage** |
| --- | --- |
| CHRY001 | 0.2593 |
| CHRY003 | 0.0895 |
| CHRY005 | 0.0871 |
| CHRY009 | 0.2418 |
| CHRY011 | 0.0983 |
| CHRY012 | 0.0933 |
| CHRY015 | 0.0631 |
| CHRY017 | 0.0926 |
| CHRY020 | 0.2031 |
| CHRY022 | 0.1441 |
| CHRY027 | 0.0948 |
| CHRY028 | 0.1263 |
| CHRY029 | 0.088 |
| CHRY034 | 0.2306 |
| CHRY036 | 0.112 |
| CHRY037 | 0.2912 |
| CHRY041 | 0.1821 |
| CHRY046 | 0.2544 |
| CHRY050 | 0.1558 |
| CHRY055 | 0.266 |
| CHRY038 | 0.0693 |
| CHRY038-Imputed | 0.3642 |
| CHRY038B | 0.012 |
| CHRY038B-Imputed | 0.0397 |

**Table S12.** Human population genetics dataset.

**Table S13.** CHRY038 Original and Soil phenotypic comparison.

**Table S14.** Information of the software used

| **Software** | **Version** | **Reference** |
| --- | --- | --- |
| AdapterRemoval2 | 2.3.3 | ^33^ |
| prinseq | 0.20.04 | ^10^ |
| bbmap | 38.18 | ^11^ |
| Kraken2 | 2.1.2 | ^30^ |
| BWA | 0.7.17-r1188 | ^12,25^ |
| KrakenUniq | 1.0.4 | ^13,14^ |
| Sourcetracker2 | 2.0.1.dev0 | ^17^ |
| vegan | vegan 2.6-4 | ^34^ |
| krakentools | 1.2 | ^35^ |
| MapDamage2 | 2.2.1 | ^28^ |
| Qualimap2 | 2.2.2-dev | ^27^ |
| pmdtools | 0.60 | ^36^ |
| compositions | 2.0-8 | ^20^ |
| mixOmics | 6.23.4 | ^21^ |
| BLAST | 2.15.0 | ^37^ |
| MEGAN6 | 6.25.1 | ^38^ |
| schmutzi |  | ^39^ |
| GATK | 3.7 | ^40^ |
| RAxML | 8.2.12 | ^41^ |
| haplogrep3 | 3.2.1 | ^42^ |
| bcftools | 1.8 | ^26^ |
| ry_compute | 0.4 | ^43^ |
| angsd | 0.940-dirty | ^44^ |
| pcangsd | 0.940-dirty | ^45^ |
| AdmixtTools | 7.0.2 | ^46^ |
| sequenceTools | 1.5.2 | ^47^ |
| PLINK | 1.90b6.21 | ^48^ |
| QUILT | 1.0.2 | ^49^ |
| Eigensoft (smartpca) | 8.0.0 | ^50^ |
| READv2 | 2.01 | ^51^ |
| pFind | 3.2.0 | ^52^ |

**Table S15.** Original skeletal element samples.

| **Sample ID** | **Alternative ID** | **Skeletal Element** | **Reference** |
| --- | --- | --- | --- |
| CHRY038 | PSN947 | Petrous bone | ^53^ |
| CHRY051 | PSN1155 | Petrous bone | ^53^ |
| DUX010 | PSN458 | Petrous bone | ^54^ |
| DUX012 | PSN460 | Petrous bone | ^54^ |
| EDI013 | PSN559 | Petrous bone | ^55^ |
| GAM042 | PSN842 | Rib | This article |
| JDS123 | J-07 / PSN223 | Vertebrae | ^53^ |
| JDS157A | PSN403 | Rib | This article |
| NMS022 | PSN518 | Petrous bone | ^53^ |
| NMS031 | PSN509 | Petrous bone | ^53^ |
| TRM003 | PSN616 | Petrous bone | ^54^ |

**Table S16.** Metagenomic sources used for microbial source-tracking analysis.

| **SampleID** | **Source** | **Environment** | **SRA** | **Publication** |
| --- | --- | --- | --- | --- |
| modernCalculus-71 | Calculus | Oral | ERS3395764 | Velsko 2019 |
| modernCalculus-72 | Calculus | Oral | ERS3395765 | Velsko 2019 |
| modernCalculus-73 | Calculus | Oral | ERS3395766 | Velsko 2019 |
| modernCalculus-74 | Calculus | Oral | ERS3395767 | Velsko 2019 |
| modernCalculus-75 | Calculus | Oral | ERS3395768 | Velsko 2019 |
| modernCalculus-76 | Calculus | Oral | ERS3395769 | Velsko 2019 |
| modernCalculus-77 | Calculus | Oral | ERS3395770 | Velsko 2019 |
| modernCalculus-78 | Calculus | Oral | ERS3395771 | Velsko 2019 |
| modernCalculus-79 | Calculus | Oral | ERS3395772 | Velsko 2019 |
| modernCalculus-80 | Calculus | Oral | ERS3395773 | Velsko 2019 |
| ruralGut-10 | Human_ruralGut | Human gut | SRR1930145 | Rampelli 2015 |
| ruralGut-1 | Human_ruralGut | Human gut | SRR1761698 | Rampelli 2015 |
| ruralGut-2 | Human_ruralGut | Human gut | SRR1761705 | Rampelli 2015 |
| ruralGut-3 | Human_ruralGut | Human gut | SRR1761710 | Rampelli 2015 |
| ruralGut-4 | Human_ruralGut | Human gut | SRR1761718 | Rampelli 2015 |
| ruralGut-5 | Human_ruralGut | Human gut | SRR1761721 | Rampelli 2015 |
| ruralGut-6 | Human_ruralGut | Human gut | SRR1929408 | Rampelli 2015 |
| ruralGut-7 | Human_ruralGut | Human gut | SRR1930121 | Rampelli 2015 |
| ruralGut-8 | Human_ruralGut | Human gut | SRR1930123 | Rampelli 2015 |
| ruralGut-9 | Human_ruralGut | Human gut | SRR1930141 | Rampelli 2015 |
| skin-21 | Skin | Skin | SRR1631060 | Oh 2016 |
| skin-22 | Skin | Skin | SRR1631061 | Oh 2016 |
| skin-23 | Skin | Skin | SRR1631063 | Oh 2016 |
| skin-24 | Skin | Skin | SRR1631064 | Oh 2016 |
| skin-26 | Skin | Skin | SRR3184100 | Oh 2016 |
| skin-27 | Skin | Skin | SRR3184876 | Oh 2016 |
| skin-28 | Skin | Skin | SRR3189411 | Oh 2016 |
| skin-29 | Skin | Skin | SRR3189416 | Oh 2016 |
| skin-30 | Skin | Skin | SRR3189418 | Oh 2016 |
| soil-11 | Soil | Soil | ERR671927 | Supratim 2023 |
| soil-12 | Soil | Soil | ERR671931 | Supratim 2023 |
| soil-13 | Soil | Soil | ERR671933 | Supratim 2023 |
| soil-14 | Soil | Soil | ERR671934 | Supratim 2023 |
| soil-15 | Soil | Soil | ERR671935 | Supratim 2023 |
| soil-16 | Soil | Soil | ERR671936 | Supratim 2023 |
| soil-17 | Soil | Soil | ERR671938 | Supratim 2023 |
| soil-18 | Soil | Soil | ERR687883 | Supratim 2023 |
| subPlaque-31 | SubPlaque | Oral | SRR061294 | HMP 2012 |
| subPlaque-32 | SubPlaque | Oral | SRR062298 | HMP 2012 |
| subPlaque-33 | SubPlaque | Oral | SRR062299 | HMP 2012 |
| subPlaque-34 | SubPlaque | Oral | SRR513165 | HMP 2012 |
| subPlaque-35 | SubPlaque | Oral | SRR513768 | HMP 2012 |
| subPlaque-36 | SubPlaque | Oral | SRR513775 | HMP 2012 |
| subPlaque-37 | SubPlaque | Oral | SRR514202 | HMP 2012 |
| subPlaque-38 | SubPlaque | Oral | SRR514239 | HMP 2012 |
| subPlaque-39 | SubPlaque | Oral | SRR514306 | HMP 2012 |
| subPlaque-40 | SubPlaque | Oral | SRR514329 | HMP 2012 |
| supPlaque-41 | SupPlaque | Oral | SRR061192 | HMP 2012 |
| supPlaque-42 | SupPlaque | Oral | SRR061320 | HMP 2012 |
| supPlaque-43 | SupPlaque | Oral | SRR061365 | HMP 2012 |
| supPlaque-44 | SupPlaque | Oral | SRR061562 | HMP 2012 |
| supPlaque-45 | SupPlaque | Oral | SRR062083 | HMP 2012 |
| supPlaque-46 | SupPlaque | Oral | SRR063517 | HMP 2012 |
| supPlaque-47 | SupPlaque | Oral | SRR1804664 | HMP 2012 |
| supPlaque-48 | SupPlaque | Oral | SRR1804823 | HMP 2012 |
| supPlaque-49 | SupPlaque | Oral | SRR512767 | HMP 2012 |
| supPlaque-50 | SupPlaque | Oral | SRR513828 | HMP 2012 |
| urbanGut-51 | Human_urbanGut | Oral | SRR059389 | HMP 2012 |
| urbanGut-52 | Human_urbanGut | Oral | SRR059425 | HMP 2012 |
| urbanGut-53 | Human_urbanGut | Oral | SRR059455 | HMP 2012 |
| urbanGut-54 | Human_urbanGut | Oral | SRR059917 | HMP 2012 |
| urbanGut-55 | Human_urbanGut | Oral | SRR060358 | HMP 2012 |
| urbanGut-56 | Human_urbanGut | Oral | SRR1761677 | HMP 2012 |
| urbanGut-57 | Human_urbanGut | Oral | SRR1761682 | HMP 2012 |
| urbanGut-58 | Human_urbanGut | Oral | SRR1761688 | HMP 2012 |
| urbanGut-59 | Human_urbanGut | Oral | SRR1761692 | HMP 2012 |
| urbanGut-60 | Human_urbanGut | Oral | SRR1761697 | HMP 2012 |
| CowRumen-10678_0008 | Ruminant_Gut | Ruminant Gut | ERR3201382 | Stewart 2019 |
| CowRumen-10678_0010 | Ruminant_Gut | Ruminant Gut | ERR3201384 | Stewart 2019 |
| CowRumen-10678_0011 | Ruminant_Gut | Ruminant Gut | ERR3201385 | Stewart 2019 |
| CowRumen-10678_0015 | Ruminant_Gut | Ruminant Gut | ERR3201388 | Stewart 2019 |
| CowRumen-10678_0016 | Ruminant_Gut | Ruminant Gut | ERR3201389 | Stewart 2019 |
| CowRumen-10678_0020 | Ruminant_Gut | Ruminant Gut | ERR3201390 | Stewart 2019 |
| FarmSoil-farmland1 | Farm_Soil | Soil | SRR22318749 |  |
| FarmSoil-farmland2 | Farm_Soil | Soil | SRR22318750 |  |
| FarmSoil-farmland3 | Farm_Soil | Soil | SRR22318751 |  |
| SheepGut-SheepGut1 | Ruminant_Gut | Ruminant Gut | SRR17509605 | Su 2022 |
| SheepGut-SheepGut2 | Ruminant_Gut | Ruminant Gut | SRR17509606 | Su 2022 |
| SheepGut-SheepGut3 | Ruminant_Gut | Ruminant Gut | SRR17509607 | Su 2022 |
| SheepGut-SheepGut4 | Ruminant_Gut | Ruminant Gut | SRR17509608 | Su 2022 |
| SheepGut-SheepGut5 | Ruminant_Gut | Ruminant Gut | SRR17509609 | Su 2022 |
| SheepGut-SheepGut6 | Ruminant_Gut | Ruminant Gut | SRR17509611 | Su 2022 |
| SheepGut-SheepGut8 | Ruminant_Gut | Ruminant Gut | SRR17509620 | Su 2022 |

Supplementary Bibliography

1.	Velsko, I. M. *et al.* Microbial differences between dental plaque and historic dental calculus are related to oral biofilm maturation stage. *Microbiome* **7**, 102 (2019).

2.	Rampelli, S. *et al.* Metagenome Sequencing of the Hadza Hunter-Gatherer Gut Microbiota. *Curr. Biol.* **25**, 1682–1693 (2015).

3.	Oh, J. *et al.* Temporal Stability of the Human Skin Microbiome. *Cell* **165**, 854–866 (2016).

4.	Mukherjee, S. *et al.* Twenty-five years of Genomes OnLine Database (GOLD): data updates and new features in v.9. *Nucleic Acids Res.* **51**, D957–D963 (2023).

5.	Human Microbiome Project Consortium. Structure, function and diversity of the healthy human microbiome. *Nature* **486**, 207–214 (2012).

6.	Stewart, R. D. *et al.* Compendium of 4,941 rumen metagenome-assembled genomes for rumen microbiome biology and enzyme discovery. *Nat. Biotechnol.* **37**, 953–961 (2019).

7.	Su, M. *et al.* Metagenomic Analysis Revealed Differences in Composition and Function Between Liquid-Associated and Solid-Associated Microorganisms of Sheep Rumen. *Front. Microbiol.* **13**, 851567 (2022).

8.	Tisza, M. J. & Buck, C. B. A catalog of tens of thousands of viruses from human metagenomes reveals hidden associations with chronic diseases. *Proceedings of the National Academy of Sciences* **118**, e2023202118 (2021).

9.	Lloyd-Price, J. *et al.* Strains, functions and dynamics in the expanded Human Microbiome Project. *Nature* **550**, 61–66 (2017).

10.	Schmieder, R. & Edwards, R. Quality control and preprocessing of metagenomic datasets. *Bioinformatics* **27**, 863–864 (2011).

11.	Bushnell, B. BBMap. Preprint at (2015).

12.	[Li, H. Aligning sequence reads, clone sequences and assembly contigs with BWA-MEM. *arXiv [q-bio.GN]* (2013).](http://paperpile.com/b/pp3oSU/uWoe)

13.	Breitwieser, F. P., Baker, D. N. & Salzberg, S. L. KrakenUniq: confident and fast metagenomics classification using unique k -mer counts. *Genome Biol.* **19**, 1–10 (2018).

14.	Pockrandt, C., Zimin, A. V. & Salzberg, S. L. Metagenomic classification with KrakenUniq on low-memory computers. *bioRxiv* 2022.06.01.494344 (2022) doi:[10.1101/2022.06.01.494344](http://dx.doi.org/10.1101/2022.06.01.494344).

15.	A new E-score for KrakenUniq. *Maxime Borry* <https://maximeborry.com/post/kraken-uniq/> (2022).

16.	Guellil, M. *et al.* Genomic blueprint of a relapsing fever pathogen in 15th century Scandinavia. *Proc. Natl. Acad. Sci. U. S. A.* **115**, 10422–10427 (2018).

17.	Knights, D. *et al.* Bayesian community-wide culture-independent microbial source tracking. *Nat. Methods* **8**, 761–763 (2011).

18.	[‘Finding Groups in Data’: Cluster Analysis Extended Rousseeuw et al. [R package cluster version 2.1.6]. (2023).](http://paperpile.com/b/pp3oSU/geXl)

19.	Chen, T. *et al.* The Human Oral Microbiome Database: a web accessible resource for investigating oral microbe taxonomic and genomic information. *Database* **2010**, baq013 (2010).

20.	van den Boogaart, K. G. & Tolosana-Delgado, R. ‘compositions’: A unified R package to analyze compositional data. *Comput. Geosci.* **34**, 320–338 (2008).

21.	Rohart, F., Gautier, B., Singh, A. & Lê Cao, K.-A. mixOmics: An R package for ’omics feature selection and multiple data integration. *PLoS Comput. Biol.* **13**, e1005752 (2017).

22.	Fellows Yates, J. A. *et al.* The evolution and changing ecology of the African hominid oral microbiome. *Proc. Natl. Acad. Sci. U. S. A.* **118**, (2021).

23.	Meng, G., Li, Y., Yang, C. & Liu, S. MitoZ: a toolkit for animal mitochondrial genome assembly, annotation and visualization. *Nucleic Acids Res.* **47**, e63 (2019).

24.	Schubert, M. *et al.* Improving ancient DNA read mapping against modern reference genomes. *BMC Genomics* **13**, 178 (2012).

25.	Li, H. & Durbin, R. Fast and accurate short read alignment with Burrows-Wheeler transform. *Bioinformatics* **25**, 1754–1760 (2009).

26.	Danecek, P. *et al.* Twelve years of SAMtools and BCFtools. *Gigascience* **10**, (2021).

27.	Okonechnikov, K., Conesa, A. & García-Alcalde, F. Qualimap 2: advanced multi-sample quality control for high-throughput sequencing data. *Bioinformatics* **32**, 292–294 (2016).

28.	Jónsson, H., Ginolhac, A., Schubert, M., Johnson, P. L. F. & Orlando, L. mapDamage2.0: fast approximate Bayesian estimates of ancient DNA damage parameters. *Bioinformatics* **29**, 1682–1684 (2013).

29.	Herbig, A. *et al.* MALT: Fast alignment and analysis of metagenomic DNA sequence data applied to the Tyrolean Iceman. *bioRxiv* 050559 (2016) doi:[10.1101/050559](http://dx.doi.org/10.1101/050559).

30.	Wood, D. E., Lu, J. & Langmead, B. Improved metagenomic analysis with Kraken 2. *Genome Biol.* **20**, 257 (2019).

31.	Kim, D., Song, L., Breitwieser, F. P. & Salzberg, S. L. Centrifuge: rapid and sensitive classification of metagenomic sequences. *Genome Res.* **26**, 1721–1729 (2016).

32.	Vogel, N. A. *et al.* euka: Robust tetrapodic and arthropodic taxa detection from modern and ancient environmental DNA using pangenomic reference graphs. *Methods Ecol. Evol.* **14**, 2717–2727 (2023).

33.	Schubert, M., Lindgreen, S. & Orlando, L. AdapterRemoval v2: rapid adapter trimming, identification, and read merging. *BMC Res. Notes* **9**, 88 (2016).

34.	Dixon, P. VEGAN, a package of R functions for community ecology. *J. Veg. Sci.* **14**, 927–930 (2003).

35.	Lu, J. *et al.* Metagenome analysis using the Kraken software suite. *Nat. Protoc.* **17**, 2815–2839 (2022).

36.	Skoglund, P. *et al.* Separating endogenous ancient DNA from modern day contamination in a Siberian Neandertal. *Proc. Natl. Acad. Sci. U. S. A.* **111**, 2229–2234 (2014).

37.	Altschul, S. F., Gish, W., Miller, W., Myers, E. W. & Lipman, D. J. Basic local alignment search tool. *J. Mol. Biol.* **215**, 403–410 (1990).

38.	Beier, S., Tappu, R. & Huson, D. H. Functional Analysis in Metagenomics Using MEGAN 6. in *Functional Metagenomics: Tools and Applications* (eds. Charles, T. C., Liles, M. R. & Sessitsch, A.) 65–74 (Springer International Publishing, Cham, 2017).

39.	Renaud, G., Slon, V., Duggan, A. T. & Kelso, J. Schmutzi: estimation of contamination and endogenous mitochondrial consensus calling for ancient DNA. *Genome Biol.* **16**, 224 (2015).

40.	McKenna, A. *et al.* The Genome Analysis Toolkit: a MapReduce framework for analyzing next-generation DNA sequencing data. *Genome Res.* **20**, 1297–1303 (2010).

41.	Stamatakis, A. RAxML version 8: a tool for phylogenetic analysis and post-analysis of large phylogenies. *Bioinformatics* **30**, 1312–1313 (2014).

42.	Schönherr, S., Weissensteiner, H., Kronenberg, F. & Forer, L. Haplogrep 3 - an interactive haplogroup classification and analysis platform. *Nucleic Acids Res.* **51**, W263–W268 (2023).

43.	Skoglund, P., Storå, J., Götherström, A. & Jakobsson, M. Accurate sex identification of ancient human remains using DNA shotgun sequencing. *J. Archaeol. Sci.* **40**, 4477–4482 (2013).

44.	Korneliussen, T. S., Albrechtsen, A. & Nielsen, R. ANGSD: Analysis of Next Generation Sequencing Data. *BMC Bioinformatics* **15**, 356 (2014).

45.	Meisner, J. & Albrechtsen, A. Inferring Population Structure and Admixture Proportions in Low-Depth NGS Data. *Genetics* **210**, 719–731 (2018).

46.	Patterson, N. *et al.* Ancient admixture in human history. *Genetics* **192**, 1065–1093 (2012).

47.	Schiffels, S. *sequenceTools*. (Github).

48.	Purcell, S. *et al.* PLINK: a tool set for whole-genome association and population-based linkage analyses. *Am. J. Hum. Genet.* **81**, 559–575 (2007).

49.	Davies, R. W. *et al.* Rapid genotype imputation from sequence with reference panels. *Nat. Genet.* **53**, 1104–1111 (2021).

50.	Price, A. L. *et al.* Principal components analysis corrects for stratification in genome-wide association studies. *Nat. Genet.* **38**, 904–909 (2006).

51.	Alaçamlı, E. *et al.* READv2: Advanced and user-friendly detection of biological relatedness in archaeogenomics. *bioRxiv* 2024.01.23.576660 (2024) doi:[10.1101/2024.01.23.576660](http://dx.doi.org/10.1101/2024.01.23.576660).

52.	Wang, L.-H. *et al.* pFind 2.0: a software package for peptide and protein identification via tandem mass spectrometry. *Rapid Commun. Mass Spectrom.* **21**, 2985–2991 (2007).

53.	Hui, R. *et al.* Genetic history of Cambridgeshire before and after the Black Death. *Sci Adv* **10**, eadi5903 (2024).

54.	Scheib, C. L. *et al.* Local population structure in Cambridgeshire during the Roman occupation. *bioRxiv* 2023.07.31.551265 (2023) doi:[10.1101/2023.07.31.551265](http://dx.doi.org/10.1101/2023.07.31.551265).

55.	Keller, M. *et al.* Ancient Yersinia pestis genomes from across Western Europe reveal early diversification during the First Pandemic (541-750). *Proc. Natl. Acad. Sci. U. S. A.* **116**, 12363–12372 (2019).
